# Retinal nerve fibre layer thickness reflects characteristics of brain grey and white matter

**DOI:** 10.1101/2025.05.13.653685

**Authors:** N. Ayyıldız, K. Mueller, S. Hardikar, F. Beyer, C. Enzenbach, R. Baber, K. Wirkner, S. Zachariae, J. Girbardt, J. D. Hassett, A. Anwander, T. Elze, M. Wang, A. V. Witte, F. G. Rauscher, A. Villringer

## Abstract

The retina is a relatively accessible part of the central nervous system compared to the brain. Using high resolution optical imaging we investigated the relationship between retinal thickness, obtained with optical coherence tomography, and structural features of the brain obtained with magnetic resonance imaging. In a population-based sample of over 500 subjects, we hypothesized: (i) that there are structural associations between circumpapillary retinal nerve fibre layer thickness and brain grey matter density and white matter microstructural properties in visual information processing areas, and specifically contralateral associations for nasal retinal fibers, and (ii) that retinal findings reflect broader changes in brain grey and white matter related to cardiovascular risk factors. In support of the first hypothesis, we showed associations of circumpapillary retinal nerve fibre layer thickness with visual cortex grey matter density and with optic radiation fractional anisotropy. These correlations were stronger for the right eye, possibly reflecting right ocular dominancy. Regarding the second hypothesis, while we confirmed the broad impact of cardiovascular risk factors such as body mass index, diabetes, and hypertension on brain structure, we didn’t find (adequate) significant partial correlations between circumpapillary retinal nerve fibre layer thickness and cardiovascular risk factors to support the hypothesis. As such, we couldn’t confirm that circumpapillary retinal nerve fibre layer thickness is associated with the impact of cardiovascular risk factors on the brain structure. However, when the effects of cardiovascular risk factors were accounted for statistically, circumpapillary retinal nerve fibre layer thickness (particularly on the right side) was associated with fractional anisotropy of limbic system tracts, i.e., the fornix and stria terminalis including hippocampus and amygdala. To further explore the structural associations between eye and brain, in terms of a possible common underlying pathology related to cardiovascular risk factors and progressive neurodegenerative diseases on the central nervous system, longitudinal and interventional studies are necessary.

## INTRODUCTION

The retina develops embryonically from the neural plate, together with the brain and spinal cord, to form the central nervous system (CNS). As part of the CNS, the retina and brain share common features of anatomy and function (London et al., 2013; Yap et al., 2019), and have been shown to share pathologies associated with neurodegenerative diseases (Casciano et al., 2024; Jabbehdari et al., 2024; Levin et al., 2022; Wolfrum et al., 2022). In contrast to the brain, the retina has a more regular neural architecture and is accessible for direct and *noninvasive* optical examination; in particular, optical coherence tomography (OCT) allows visualization of the retinal neural architecture with microscopic resolution *in vivo* (Doustar et al., 2017; Moons & De Groef, 2022; Vujosevic et al., 2022; Xie et al., 2022). Here, we investigate whether the retinal information obtained with OCT can serve as an indicator of the brain structure measured with magnetic resonance imaging (MRI).

There are two potential connections by which the retina may reflect the state of the brain. First, the retina is directly connected to the brain via the optic nerve, which projects to the occipital cortices via the optic radiation. Previous studies have suggested an association between circumpapillary retinal nerve fiber layer thickness (RNFLT, around optic disc) and structural properties in bilateral occipital regions (Mutlu et al., 2018; Ueda et al., 2022). Given the decussating fibers in the optic chiasm, coming solely from areas of the retina nearest to the nose (Girbardt et al., 2021), one would furthermore expect that the changes in left and right nasal RNFLT would be associated specifically with grey and white matter changes in contralateral visual brain areas.

The second possible connection between the retina and the brain is indirect, via other biological systems, resulting in joint sensitivity to various pathophysiological processes related to e.g., cardiovascular risk factors (CVRF) or neurodegeneration. CVRF have been suggested to have analogous effects on various brain and eye conditions (Barreiro-Gonzalez et al., 2020; Brain et al., 2023; Casson et al., 2021; Chrysou et al., 2019; Ebrahimi et al., 2023; Hainsworth et al., 2024; Jabbehdari et al., 2024; Langner et al., 2022; Lee et al., 2022; Ravi Teja et al., 2017; Yap et al., 2019; Zhou et al., 2021). CVRF such as higher body mass index (BMI), lower physical activity, arterial hypertension, smoking and diabetes mellitus have been shown to be associated with thinning of the retinal nerve fiber layer (Chen et al., 2023; Colijn et al., 2019; Li et al., 2020; Majithia et al., 2022; Mauschitz et al., 2018; Nousome et al., 2021; Rauscher et al., 2021; Wang et al., 2013). Likewise, it has been well established that CVRF are associated with a reduction in grey and white matter density throughout the brain, including frontal (e.g., (pre)frontal, precentral, supplementary motor, orbitofrontal, and anterior cingulate cortices), temporal (e.g., hippocampal, para-hippocampal, entorhinal, insular, and olfactory regions), occipito-parietal (e.g., primary visual cortex, cuneus, and precuneus) and subcortical (e.g., striatum and amygdala) regions as well as the cerebellum (Beyer, Garcia-Garcia, et al., 2019; Beyer, Kharabian Masouleh, et al., 2019; Elbejjani et al., 2019; Erickson et al., 2014; Erickson et al., 2010; Fritz et al., 2014; Gianaros et al., 2006; Jackow-Nowicka et al., 2021; Jennings et al., 2012; Kharabian Masouleh et al., 2016; Raz et al., 2003; Raz, Rodrigue, & Haacke, 2007; Raz, Rodrigue, Kennedy, et al., 2007; Roy et al., 2020; Schaare et al., 2019; Ward et al., 2010; Yu et al., 2021; Zhang et al., 2018).

In addition to the potential impact of CVRF on both retinal and brain structure, specific (neuro)pathologies related to major neurodegenerative diseases have been detected in primate and non-primate retina. These include amyloid-beta and/or phosphorylated-tau aggregation in patients with Alzheimer’s disease (Arouche-Delaperche et al., 2023; den Haan et al., 2018; Koronyo et al., 2017; Lam et al., 2021) and alpha-synuclein aggregation and dopaminergic cell loss in patients with Parkinson’s disease (Hart de Ruyter et al., 2023; La Morgia et al., 2013; Ortuno-Lizaran et al., 2020). Given such potential joint pathologies, various population (Barrett-Young et al., 2023; Chua et al., 2021; Lima Reboucas et al., 2023; Mauschitz et al., 2022; Mendez-Gomez et al., 2018; Mutlu et al., 2017; Mutlu et al., 2018; Ong et al., 2015; Ueda et al., 2022; van der Heide et al., 2024) as well as non-population studies (Byun et al., 2021; Casaletto et al., 2017; Lopez-de-Eguileta et al., 2022; Mejia-Vergara et al., 2021; Mendez-Gomez et al., 2018; Moran et al., 2022; Shi et al., 2020; Shi et al., 2019; Uchida et al., 2020) have suggested associations between the retina and specific brain regions including the limbic system and basal ganglia. Taken together, the above evidence suggests there may be more widespread parallels between the retina and neural grey and white matter, beyond just anatomically connected regions.

Thanks to the population-based LIFE-Adult-Study (Loeffler et al., 2015), we were able to work with RNFLT from OCT scans at microscopic resolution (um) as well as high-resolution T1-weighted (mm) and diffusion-weighted MRI for the detailed evaluation of brain grey and white matter. This multimodal approach allowed for integrative analysis of retinal and brain structure within the same individuals. The LIFE-Adult-Study also included CVRF, and notably, no prior studies have examined the concurrent impact of CVRF on whole-brain structure and retinal measures at a population level. Sectoral RNFLT data further enabled the assessment of nasal retinal fiber associations with contralateral visual brain regions for the first time. We aimed to confirm known anatomical connections and investigate how including and excluding the effects of CVRF influences retina-brain structural correlations at whole-brain level.

Based on the above considerations, we tested two hypotheses: (i) There are associations between RNFLT and grey matter density and white matter microstructural properties in directly connected brain areas i.e., visual cortex and the optic radiation, and specifically contralateral associations for RNFLT of the nasal sectors. (ii) There are associations beyond directly connected brain areas. Overall, we investigated whether the optically accessible retina might serve as a window into anatomically connected as well as other brain regions associated with cardiovascular risk factors and/or neurodegeneration.

## 2. METHODS

We preregistered the study’s hypotheses, relevant methods, and analysis plan here: https://osf.io/mdgq3/registrations under the ‘eye-brain VBM project’ with https://doi.org/10.17605/OSF.IO/MDGQ3 at Open Science Forum (OSF) platform). The scripts used in this study can be found here: https://github.com/n-ayyildiz/EyeBrain_project.git

### 2.1. Sample and Variables

Data were obtained from the population-based LIFE-Adult-Study, which included deeply phenotyped 10000 human participants recruited from the general population of Leipzig, Germany (Engel et al., 2022; Enzenbach et al., 2019; Loeffler et al., 2015). Here, we used cross-sectional data from the baseline examination (conducted between 2011 and 2014) having OCT, MRI, and cardiovascular risk assessments. We excluded participants with relevant eye, brain, or CNS diseases as well as those with poor OCT and/or MRI data quality. **Figure 1** gives an overview of inclusion/exclusion and the number of participants with OCT and two MRI modalities, T1-weighted and DWI data. Taking the CVRF into account, we included 769 participants for the analysis of OCT and grey matter density using T1-weighted imaging and 550 participants for the analysis of OCT and white matter microstructure using DWI (Results, Table 1).

**Figure 1.**
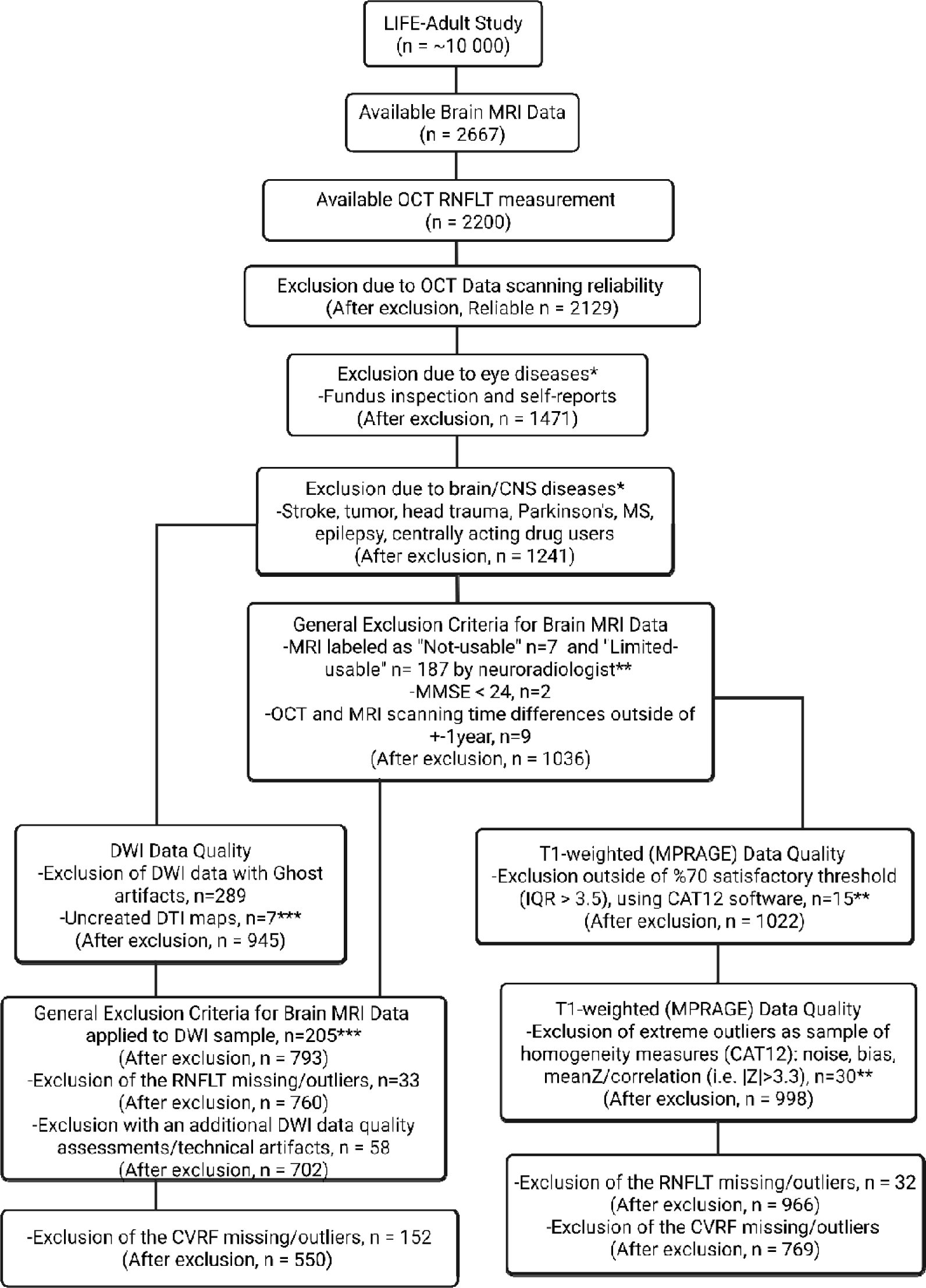
Sample recruitment with implemented inclusion and exclusion criteria. Asterisks show partial overlapping between the subgroups. Final numbers of subjects with usable brain MRI and usable RNFLT of subjects with both eyes healthy were n = 550 for the white matter microstructure and n = 769 for the grey matter density investigations. n: number of participants, MRI: magnetic resonance imaging, OCT: optical coherence tomography, RNFLT: circumpapillary retinal nerve fibre layer thickness, CNS: central nervous system, MS: multiple sclerosis, MMSE: mini-mental state examination, DWI: diffusion-weighted imaging, DTI: diffusion-tensor imaging, MPRAGE: magnetization prepared rapid gradient echo, IQR: index of quality rating, CAT12: computational anatomy toolbox version 12.

**Table 1.**
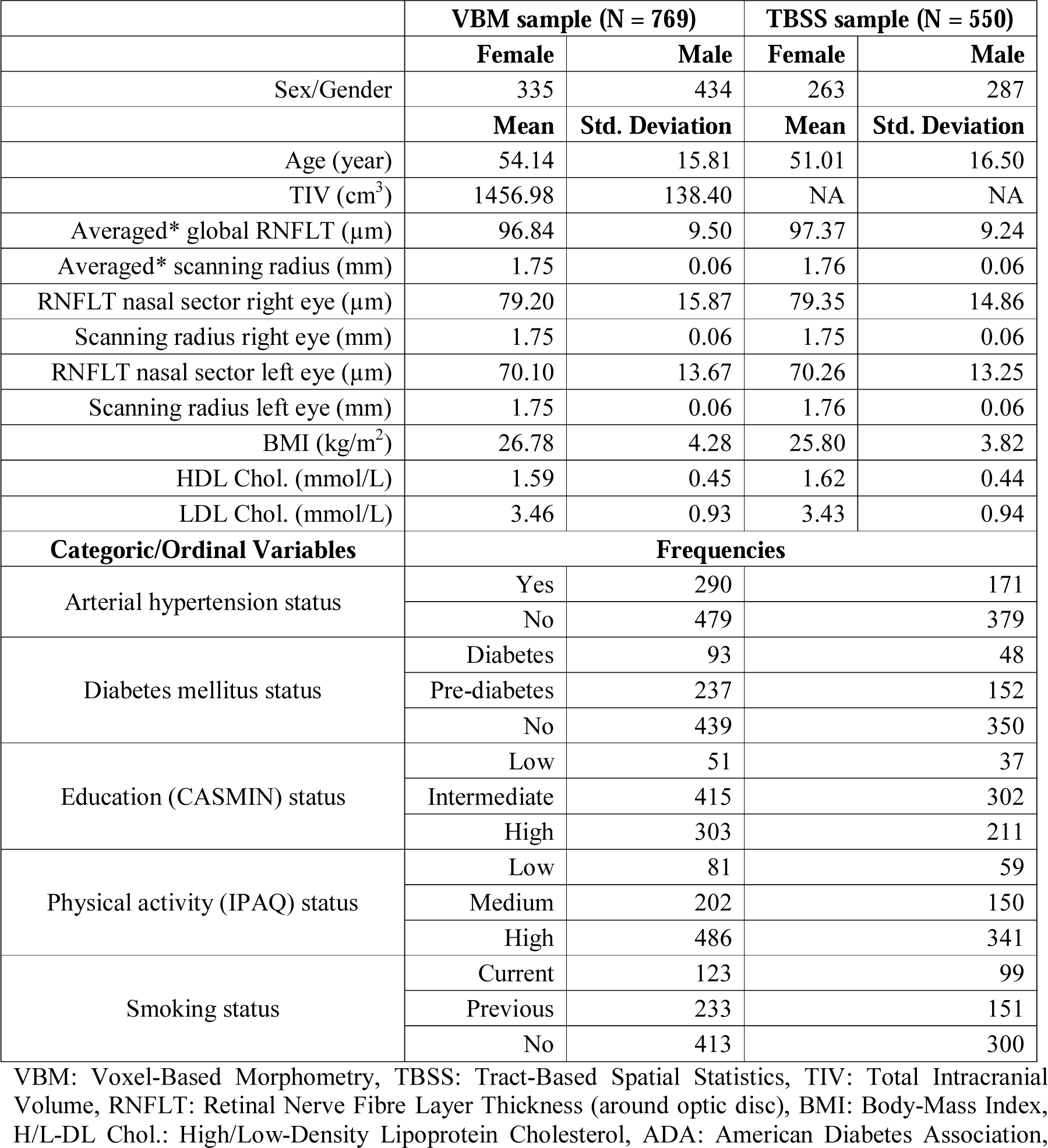
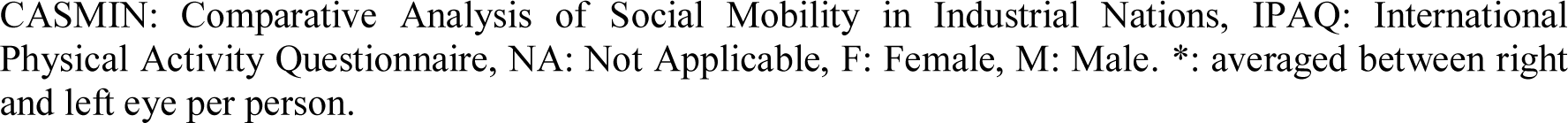
Descriptive statistics of the samples for the whole-brain VBM and TBSS analyses.

|  | VBM sample (N = 769) |  | TBSS sample (N = 550) |  |
| --- | --- | --- | --- | --- |
|  | Female | Male | Female | Male |
| Sex/Gender | 335 | 434 | 263 | 287 |
|  | Mean | Std. Deviation | Mean | Std. Deviation |
| Age (year) | 54.14 | 15.81 | 51.01 | 16.50 |
| TIV (cm<sup>3</sup>) | 1456.98 | 138.40 | NA | NA |
| Averaged* global RNFLT (μm) | 96.84 | 9.50 | 97.37 | 9.24 |
| Averaged* scanning radius (mm) | 1.75 | 0.06 | 1.76 | 0.06 |
| RNFLT nasal sector right eye (μm) | 79.20 | 15.87 | 79.35 | 14.86 |
| Scanning radius right eye (mm) | 1.75 | 0.06 | 1.75 | 0.06 |
| RNFLT nasal sector left eye (μm) | 70.10 | 13.67 | 70.26 | 13.25 |
| Scanning radius left eye (mm) | 1.75 | 0.06 | 1.76 | 0.06 |
| BMI (kg/m<sup>2</sup>) | 26.78 | 4.28 | 25.80 | 3.82 |
| HDL Chol. (mmol/L) | 1.59 | 0.45 | 1.62 | 0.44 |
| LDL Chol. (mmol/L) | 3.46 | 0.93 | 3.43 | 0.94 |
| Categoric/Ordinal Variables | Frequencies |  |  |  |
| Arterial hypertension status | Yes | 290 | 171 |  |
|  | No | 479 | 379 |  |
| Diabetes mellitus status | Diabetes | 93 | 48 |  |
|  | Pre-diabetes | 237 | 152 |  |
|  | No | 439 | 350 |  |
| Education (CASMIN) status | Low | 51 | 37 |  |
|  | Intermediate | 415 | 302 |  |
|  | High | 303 | 211 |  |
| Physical activity (IPAQ) status | Low | 81 | 59 |  |
|  | Medium | 202 | 150 |  |
|  | High | 486 | 341 |  |
| Smoking status | Current | 123 | 99 |  |
|  | Previous | 233 | 151 |  |
|  | No | 413 | 300 |  |

VBM: Voxel-Based Morphometry, TBSS: Tract-Based Spatial Statistics, TIV: Total Intracranial Volume, RNFLT: Retinal Nerve Fibre Layer Thickness (around optic disc), BMI: Body-Mass Index, H/L-DL Chol.: High/Low-Density Lipoprotein Cholesterol, ADA: American Diabetes Association,
CASMIN: Comparative Analysis of Social Mobility in Industrial Nations, IPAQ: International Physical Activity Questionnaire, NA: Not Applicable, F: Female, M: Male. \*: averaged between right and left eye per person.

All variables were measured within the population-based LIFE-Adult-Study and described in detail elsewhere (Engel et al., 2022; Enzenbach et al., 2019; Loeffler et al., 2015). In the following, we briefly describe the variables used in the present study.

Circumpapillary RNFLT (RNFLT around optic nerve head), referred to ‘RNFLT’ in this manuscript, was computed from OCT scans of the retina. Only participants with RNFLT measurements available for both eyes were included in the analyses. We calculated, per participant, an average value between the individual’s left and right ‘global mean of the RNFLT’ measures, referred to ‘averaged global RNFLT’ in this manuscript. Additionally, we calculated, per participant, an average scanning radius value between the individual’s left and right ‘retinal scanning radius’ measures, we termed this value the ‘averaged retinal scanning radius’. For the laterality analysis (see section 2.4.7), we analyzed left and right global RNFLT separately using the respective retinal scanning radius i.e., for left and right eyes.

We furthermore investigated the RNFLT specifically of the nasal sectors to test the laterality of their projection to the brain [see (Girbardt et al., 2021)’s illustration of the ‘Eye to brain projection’ as Figure 2 at the publication]. The nasal sector for either eye allows investigation of the contralateral projection to the brain, as nasal RNFLT marks a retinal area where only decussating fibers in the optic chiasm are located.

**Figure 2.**
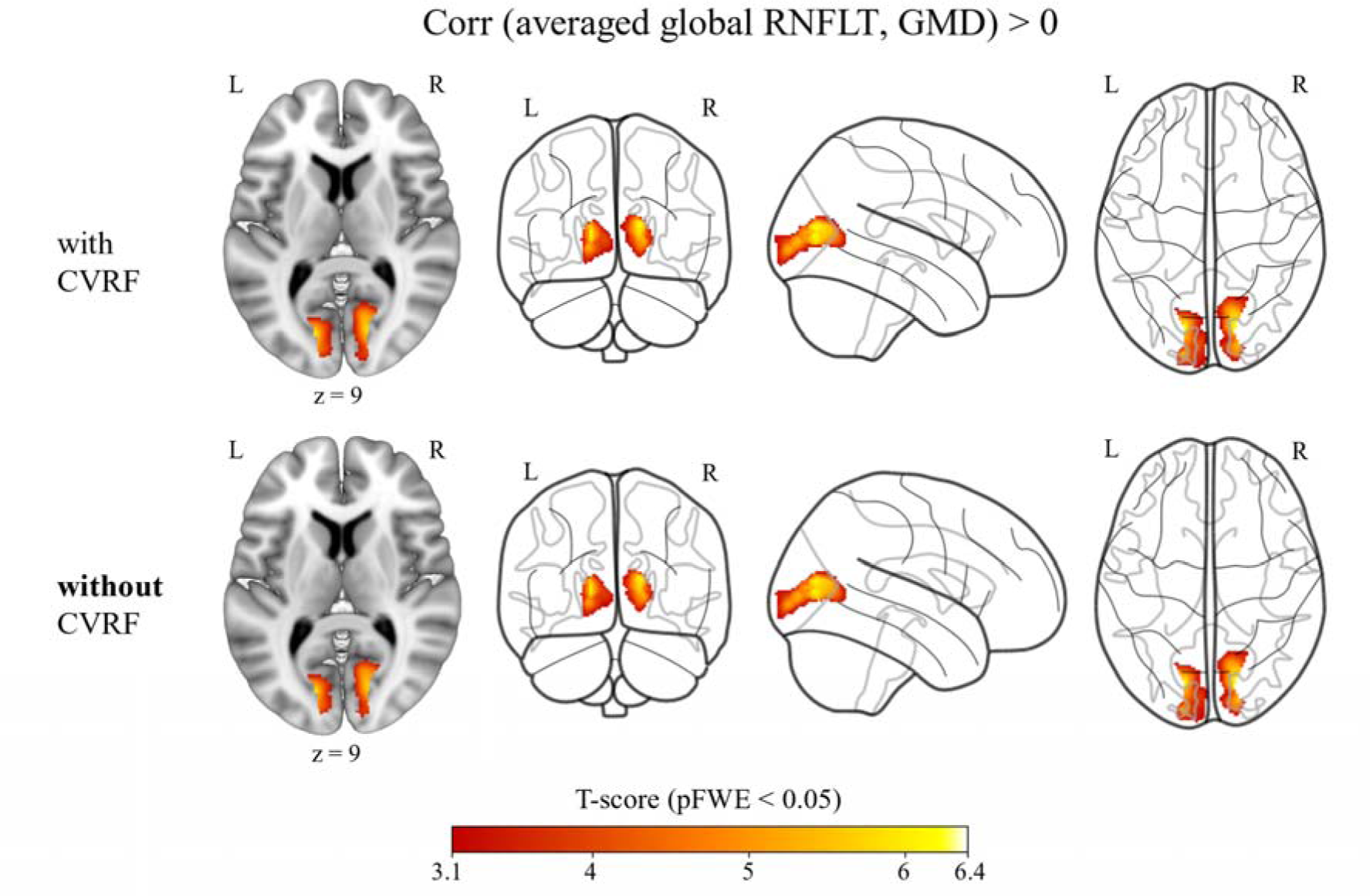
Averaged (between right and left eye per person) global RNFLT showed positive correlations (red overlay) with brain GMD when controlling for age, sex, TIV, and averaged retina scanning radius and CVRF (i.e., BMI, LDL and HDL cholesterol scores, and diabetes mellitus, arterial hypertension, smoking, and physical activity status) and when controlling only for age, sex, TIV, and averaged retina scanning radius **without CVRF** [n = 769]. Results shown on MNI152_T1_1mm standard template and glass brain using Mango and python glass-brain packages, neurological view with the left hemisphere on the left side. Coordinates (12, −72, 9). Color bar shows T statistics corrected at cluster-level pFWE < 0.05 and uncorrected at voxel-level p < 0.001. RNFLT: retinal nerve fibre layer thickness, GMD: grey matter density, TIV: total intracranial volume, CVRF: cardiovascular risk factors, BMI: body-mass index, LDL: low-lipoprotein density, HDL: high-lipoprotein density, MNI: Montreal Neurological Institute, FWE: family-wise error correction.

Grey matter density (GMD) and total intracranial volume (TIV) were computed within the framework of voxel-based morphometry [VBM, (Ashburner & Friston, 2000)] using T1-weighted MRI. We used GMD for ‘modulated grey matter probability’ obtained with VBM.

White matter microstructure (WMM) properties, in particular, fractional anisotropy (FA) and mean diffusivity (MD) values, were computed within tract-based spatial statistics [TBSS, (Smith et al., 2006)] using DWI.

Education status was defined with three ordinal categories as low, intermediate, or high education levels according to (Brauns et al., 2003), coded as 1, 2 and 3, respectively.

#### Cardiovascular Risk Factors (CVRF)

Arterial hypertension status was defined with two categories as: presence or absence of hypertension, coded as 1 and 0, respectively. It was obtained by either a self-reported physician-based diagnosis, intake of antihypertensive drugs, or having a systolic blood pressure > 140 mmHg and a diastolic blood pressure > 90 mmHg measured three times with consecutive 5-minute intervals.

Diabetes mellitus status was defined with three categories as: normal glucose tolerance, prediabetes, or diabetes, coded as 0, 1, and 2, respectively. The categories were defined using fasting blood sugar levels, oral glucose tolerance test, and HbA1C levels according to the America Diabetes Association criteria (American Diabetes, 2021) see more details here (Rauscher et al., 2024).

Smoking status was defined with three categories as: non-smoker, previous smoker, current smoker, coded as 0, 1, and 2, respectively.

Physical activity status was defined with three ordinal categories as: low, medium, and intensive activity from the International Physical Activity Questionnaire (Craig et al., 2003; Meh et al., 2021), coded as 1, 2, and 3, respectively.

Age was defined in years and Body-mass index (BMI) in kg/m^2^ using anthropometric measurement values. Low-density lipoprotein (LDL) and high-density lipoprotein (HDL) levels were acquired with blood sampling. Sex/gender was defined with two categories, female and male, as coded 1 and 2, respectively.

### 2.2. Optical coherence tomography (OCT) imaging of the retina

OCT utilizes laser light, which is differentially reflected by distinct retinal tissue layers, enabling a histology-like, *in vivo* examination of retinal structures. The system employs two simultaneous laser beams i.e., A- and B-scans to generate both a high-resolution retinal cross-sectional scan. In addition, a real-time fundus image is created by confocal scanning laser ophthalmoscopy (cSLO), facilitating precise alignment and measurement during the scanning procedure. OCT imaging was performed using spectral-domain-OCT (Spectralis, Heidelberg Engineering GmbH, Heidelberg, Germany), which incorporates live eye-tracking to enhance scan accuracy. The simultaneous acquisition of the fundus image enabled precise registration of scan locations, while the eye-tracking system minimized motion artifacts by aligning retinal landmarks in real time, thereby ensuring accurate image capture despite involuntary eye movements. The cSLO image was conducted using a wavelength of 815 nm, while OCT scans were acquired using a super luminescent diode with a central wavelength of 870 nm (range: 850–920 nm).

A scanning rate of 40,000 A-scans per second and real-time noise reduction algorithms were employed to enhance the signal-to-noise ratio. Signal processing techniques, including image averaging, were utilized to improve scan clarity. At each scanning location, 100 individual B-scans were acquired and averaged to reduce speckle noise, thereby enhancing the visibility of fine retinal structures. (Circumpapillary) RNFLT was measured with a resolution of 768 equidistant points on a measurement circle of 6-degree visual angle corresponding to an approximate scanning radius of 1.7 mm from the centre of the optic nerve head (See Supplement Figure S19 as an example). The scans were captured in high-speed mode, with each B-scan comprising 496 A-scans, corresponding to 768 pixels by 496 pixels. The Spectralis OCT system achieves an axial imaging depth of 1.9 mm, corresponding to a digital axial resolution of 3.9 μm and an optical resolution of 7 μm. Examination time for RNFLT scan per eye amounted to 1 minute.

Additionally, fundus images (Nidek AFC-230; Oculus, Wetzlar, Germany) of undilated eyes were obtained from all participants. All images were evaluated by two independent, ophthalmologically trained observers who classified retinal or optic nerve abnormalities based on current standards. Ophthalmological and clinical phenotyping, as well as data collection, have been described in more detail elsewhere (Baniasadi et al., 2020; Li et al., 2020; Rauscher et al., 2021).

### 2.3. Magnetic resonance imaging (MRI) of the brain

A 3 Tesla Siemens Verio MRI scanner (Siemens Healthineer, Erlangen, Germany) was used to acquire the brain images. Raw MR images with all modalities were investigated first by an expert neuroradiologist and first exclusions were made according to data usability conditions.

#### 2.3.1. T1-weighted imaging

Anatomical T1-weighted images were acquired using an MPRAGE (magnetization prepared rapid gradient echo) protocol with the following parameters: inversion time [TI]: 900 ms; repetition time [TR]: 2300 ms; echo time [TE]: 2.98 ms; flip angle: 9°; band width: 240 Hz/pixel; field of view [FOV]: 256 mm x 240 mm x 176 mm; image matrix: 256 x 240; sagittal orientation; voxel size: 1 mm x 1 mm x 1 mm, acquisition time: 5 min 6 s.

#### 2.3.2. Diffusion-weighted imaging (DWI)

Diffusion-weighted images were acquired using a double spin-echo encoding sequence with the following parameters: [TR]: 13800 ms; [TE]: 100 ms; [FOV]: 220 mm x 220 mm x 123 mm; image matrix: 128 x 128; voxel size: 1.7 mm x 1.7 mm x 1.7 mm; maximum b-value: 1000 s/mm^2^ in 60 directions, and 7 volumes with b-value of 0 s/mm^2^; GRAPPA 2, 32 channel head coil, acquisition time: 16 min 8 s.

### 2.4. Statistical analysis

#### 2.4.1. Sample description

SPSS 24 (PASW, SPSS, IBM) was used for descriptive statistics of the sample. Outliers i.e., the data points outside of |z| = 3.3 of the distribution and missing data points for each variable were excluded for the respective analysis. For correlation analysis with the CVRF, averaged global RNFLT and averaged retina scanning radius variables were used. Categorical variables were dummy coded as mentioned in the methods section of ‘2.1. Sample and Variables’.

#### 2.4.2. OCT data analysis

Reliability criteria for the OCT scan were based on: (i) image quality with signal-to-noise-ratio ≥20[dB; (ii) average number of B-scans ≥50; and (iii) no more than 5% missing or unreliable RNFLT segmentations among the 768 A-scans, which are the basis of the average sectorial data. All evaluated scans were adequately centered on the optic nerve head.

We used averaged global RNFLT, as the main regressor to predict individual brain grey and white matter measures in multivariable regression models (See sections 2.4.3 and 2.4.5). Additionally, separate analyses for right and left eye of global RNFLT were carried out for the laterality assessment (see section 2.4.7).

As potential confounding variables, we added age, sex, and retina scanning radius in the regression models due to the following considerations: Previous studies have demonstrated strong relationships between RNFLT and age (Wang et al., 2017) and sex (Li et al., 2020). In addition, it has been shown that ocular magnification effects, due to corneal curvature, lens-related myopia, or axial length, have a substantial impact on the true size of the OCT scanning circle of circumpapillary RNFLT (Li et al., 2020). The Spectralis machine estimates the true scanning radius from the individual focus settings of each retina scan. We used this estimated true retina scanning radius to account for refractive error as an additional covariate to age and sex in our analyses.

#### 2.4.3. T1-weighted data analysis using VBM

T1-weighted images were preprocessed using the CAT12 v12.8.2 [computational anatomy toolbox, https://neuro-jena.github.io/cat12-help/] (Gaser et al., 2024) in SPM12 v7771 software [statistical parametric mapping, https://www.fil.ion.ucl.ac.uk/spm/software/spm12/] (Friston et al., 2007) based on MATLAB v9.12 [The MathWorks Inc. (R2022a)] with the settings defined in our preregistration. The main steps were denoise filtering, bias correction, skull-stripping, and initial and refined registration with unified segmentation (Ashburner & Friston, 2005). Subsequently, segmented images were normalized to the MNI (Montreal Neurological Institute) template (Ashburner & Friston, 2011), and, as a final preprocessing step before further analysis, smoothed with 8 mm FWHM (full-width at half maximum). We checked the raw and preprocessed images before and after smoothing with the data quality rating scale provided by CAT12. We first excluded all T1-weighted data rated below “satisfactory (i.e., scored under 3.5 or 70 %)” criterion for the VBM (Gaser et al., 2024). We then used z-scores to create a homogeneous sample for large data sets and excluded the images with |z| = 3.3 as outliers after additional visual inspections. Voxels were masked with an absolute threshold of 0.1 for the VBM analysis.

Whole brain VBM (Ashburner & Friston, 2000) analysis using a multiple regression design for cross-sectional data was performed using the CAT12. GMD values were predicted by RNFLT and/or CVRF, always adjusting for age, sex, TIV, and retina scanning radius in the general linear model: GLM [Brain GMD =[β_0_ + β_1_ (RNFLT) + β_2_ (age) + β_3_ (sex) + β_4_ (retina scanning radius) + β_5_ (TIV) + β_6+_ (CVRF) + error]. Statistical significance was accepted as p < 0.05 with family-wise error (FWE) corrected for multiple comparisons at the cluster level and p < 0.001 uncorrected at voxel level. We performed whole brain VBM with averaged global RNFLT while controlling for all CVRF simultaneously in addition to age, sex, TIV, and retina scanning radius (i.e., ‘models with CVRF’) and controlling for only age, sex, TIV, and retina scanning radius (i.e., ‘models without CVRF’).

#### 2.4.4. Region-of-interest (ROI) analysis using regional and global grey matter volume

We used predefined ROIs (see Supplement Table S20) to test whether we could confirm previous findings from the literature that RNFLT measures show positive correlations with regional and global brain grey matter. We extracted regional grey matter volume (mL) per subject from their segmented and registered native spaces according to the Neuromorphometrics atlas using the CAT12. Then, we used partial correlation analyses between averaged global RNFLT and regional grey matter volume accounting for age, sex, averaged retinal scanning radius, and TIV using SPSS. We used a similar approach to that described in section ‘2.4.3. T1-weighted data analysis using VBM’, performing the partial correlation analysis with and without all CVRF. We corrected p < 0.05 significance level using false-discovery rate (FDR) for multiple comparisons, taking the number of ROIs into account.

#### 2.4.5. DWI data analysis using TBSS

The diffusion-weighted images were visually inspected for ghost artifacts related to the fat suppression settings of the scanner. They were detected in many images resulting in the exclusion of 289 participants due to the lack of suitable preprocessing methods to correct the artifact [cf. (Zhang et al., 2018)]. For the remaining data, raw diffusion-weighted images were preprocessed [i.e., removal of Gibbs-ringing, head motion correction using the ‘eddy’ tool in FSL (FMRIB Software Library, Oxford) with outlier replacement (Andersson et al., 2016; Andersson & Sotiropoulos, 2016), registration to the T1-weighted brain images in AC-PC orientation, tensor model fitting, and FA and MD computation] combining the protocols from Lipsia (Lohmann et al., 2001) and FSL diffusion toolboxes (https://fsl.fmrib.ox.ac.uk/) as previously described (Thomas et al., 2019). We visually checked the preprocessed diffusion-weighted images with the FSL “slicesdir” tool after implementing further inclusion and exclusion criteria (Figure 1). We additionally excluded some of the DWI data (n = 58) with technical problems and strongly enlarged ventricles according to total ventricle volume distribution (i.e., outside of the |z| = 3.3).

We analyzed FA and MD images using whole-brain voxel-wise TBSS (Smith et al., 2006) for the cross-sectional data with FSL software. First, preprocessed images were spatially normalized using linear and non-linear registration to the target image in MNI space (HCP1065-1mm-FA). We used 0.2 as a threshold to create mean FA skeleton image and then all images were projected according to this skeleton to create 4D FA and then MD skeletonized inputs for further analyses. We used permutation testing for GLM with 10000 permutations using the “randomise” tool (Winkler et al., 2014). Statistical significance was accepted as p < 0.05 with FWE correction for multiple comparisons at the cluster-size level and p < 0.001 uncorrected at the voxel level. We preferred cluster inference to deal with the ‘cluster-leakage’ problem (Smith & Nichols, 2009; Spisak et al., 2019). Similar to the VBM analysis, we used multiple regression models, this time predicting WMM properties, i.e., FA or MD values using the RNFLT and/or CVRF as regressors; age, sex, and retinal scan radius were always adjusted in the GLM. Additionally, we also created the models with and without CVRF covariates similar to the VBM analyses as described above (TIV were not included in the TBSS GLM).

#### 2.4.6. Conjunction analysis of global RNFLT and CVRF correlations to brain structure

The regions found using either RNFLT or each CVRF, as regressors in separate models of the VBM analyses, were compared using conjunction analyses (Nichols et al., 2005). Thus, we were able to compare the FWE-corrected correlation maps (i.e., GMD and WMM) of the ‘models with CVRF’ to the ‘models without CVRF’ to identify a possible influence of CVRF on the relationship between RNFLT and brain structure.

#### 2.4.7. Laterality of global RNFLT on brain structure

To test whether the correlations of the right and left global RNFLT with the GMD differ, an ROI analysis was performed. Using Python v3.11.5, we extracted the GLM beta values in bilateral calcarine cortex using Neuromorphometrics atlas in CAT12 toolbox. Then, we compared the regional right global RNFLT beta values with the regional left global RNFLT beta values using a paired-sample t-test with R v 4.4.3 statistical package. We assessed the WMM parameters in the same way using the GLM beta values of the bilateral optic radiation using FSL Juelich histological atlas.

#### 2.4.8. Ethics statement

The LIFE-Adult-Study was approved by the Ethics Committee of the Medical Faculty of Leipzig University in accordance with the Declaration of Helsinki. Written informed consent was obtained from all participants.

## 3. RESULTS

### 3.1. Descriptive statistics

Table 1 shows the study’s sample profile for the VBM (Table 1, left column) and TBSS (Table 1, right column) assessments. At baseline, the LIFE-Adult-Study included mainly participants between 40 and 79 years of age, with a small set of participants below 40 and down to 18 years old (Engel et al., 2022; Enzenbach et al., 2019; Loeffler et al., 2015). This was reflected in our study, furthermore, there were more men than women in our study, especially in the VBM cohort.

### 3.2. Correlations between RNFLT and CVRF

No significant correlation was observed between averaged global RNFLT and any CVRF in either the VBM or TBSS samples, when controlling for averaged retina scanning radius, age, and sex in the partial correlation analyses (Table 2). Thus, there seems to be no partial variance within the averaged global RNFLT that could be explained uniquely by each CVRF, apart from the variance shared by age, sex, and retina scanning radius.

**Table 2.** Partial correlation coefficients between averaged* global RNFLT and CVRF when controlling for averaged scanning radius, age, and sex (n = 769 VBM Sample, n = 550 TBSS Sample)

|  | Diabetes Status | BMI | Hypertension Status | IPAQ Category | LDL | HDL | Smoking Status |
| --- | --- | --- | --- | --- | --- | --- | --- |
| averaged* lobal RNFLT, df = 764 | 0.01 | 0.04 | 0.01 | -0.01 | 0.03 | 0.01 | 0.01 |
| averaged* global RNFLT, df = 545 | -0.04 | 0.07 | 0 | -0.01 | -0.01 | -0.003 | 0.05 |

p > 0.05 for all partial correlations, \*: averaged between right and left eye per person.

### 3.3. Correlations between RNFLT and brain GMD (N = 769 whole-brain VBM)

#### 3.3.1. Correlations with global RNFLT

In the whole-brain VBM multiple regression analysis, averaged global RNFLT showed significant positive correlations with brain GMD in the bilateral occipital cortex when controlling only for age, sex, TIV, and retina scanning radius (i.e., models without CVRF). The effect persisted when controlling for all CVRF in addition to age, sex, TIV, and retina scanning radius (i.e., models with CVRF) (Figure 2 and Table 3). The correlations were similar between the models with CVRF and the models without CVRF (see Supplementary Figures S1-S3 and Table S3, and Overlap section).

**Table 3.** Whole brain VBM results: Positive correlations of averaged** global RNFLT with brain GMD (controlled for all CVRF)

| cluster<br>p(FWE-corr) | cluster<br>size | peak T | peak<br>p(unc) | coordinates mm |  |  | Area* |
| --- | --- | --- | --- | --- | --- | --- | --- |
|  |  |  |  | x | y | z |  |
| 0.006 | 1498 | 6.19 | <0.0001 | 16 | -80 | 14 | Right Calcarine Cortex |
|  |  | 5.36 | <0.0001 | 18 | -90 | 4 |  |
| 0.005 | 1521 | 6.08 | <0.0001 | -15 | -80 | 9 | Left Calcarine Cortex |
|  |  | 5.36 | <0.0001 | -16 | -94 | 0 |  |
|  |  | 4.09 | <0.0001 | -8 | -90 | 4 |  |

\*: CAT12 Neuromorphometric Atlas, \*\*: between right and left eye per person

When correlating global RNFLT of the left and right eyes with brain GMD (Supplementary Figure S4 and Table S3), the right global RNFLT were spatially larger and had higher peak T-statistics in the correlations than the left global RNFLT (see Supplementary Figures S5-S6 and Table S3). These correlation differences between left and right global RNFLT were similar between the models with and without CVRF (see Supplementary Figures S7-S8). Subsequent ROI analysis showed that the right global RNFLT had significantly steeper positive correlations with GMD in bilateral calcarine cortex than the left global RNFLT (paired-sample t-test, see Supplementary Table S4).

#### 3.3.2. Correlations with nasal RNFLT

We did not find significant GMD correlations at p < 0.05 FWE-corrected at cluster level with the nasal RNFLT sector of the left or right eye, when controlling only for age, sex, TIV, and related retina scanning radius as covariates, or additionally controlling for all CVRF. According to relatively small clustered correlations at an uncorrected p < 0.001 at the voxel level (Supplement Figure S20-S21), to further test the contralaterality hypothesis, we decided to perform small volume correction (SVC) using bilateral occipital cortex mask instead of whole brain. There were significant positive correlations at p < 0.05 FWE corrected at cluster level with SVC; the RNFLT of nasal sector of the left eye correlated with the contralateral cortex while the RNFLT of nasal sector of the right eye correlated with the calcarine cortex GMD bilaterally, when CVRF were controlled for in addition to age, sex, TIV, and corresponding retina scanning radius (Figure 3). When CVRF were not used as covariates, similar results were obtained (Supplement Figure S22-S23).

**Figure 3.**
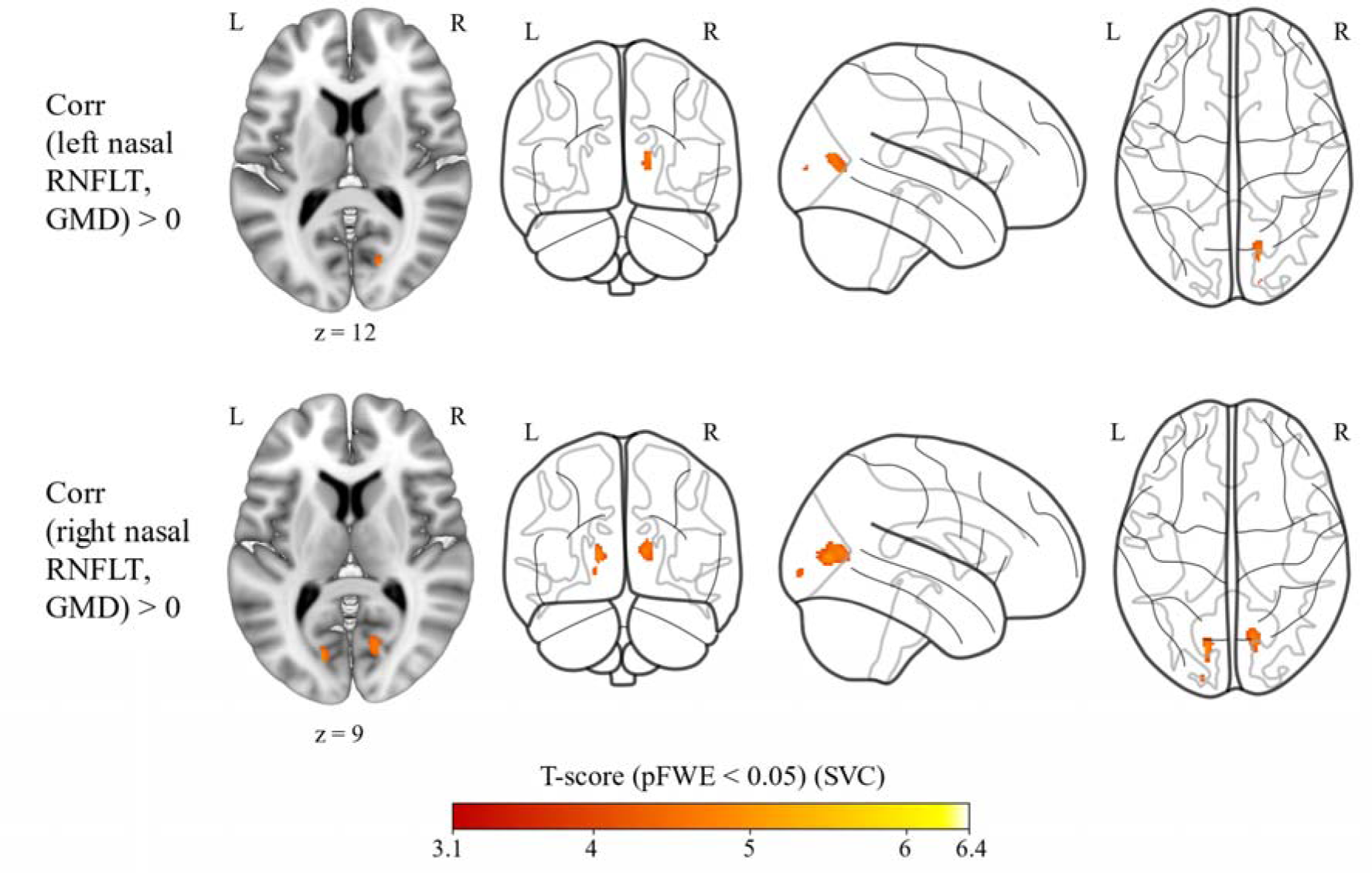
Left nasal and right nasal RNFLT showed positive correlations (red overlay) with brain GMD when controlling for age, sex, TIV, and averaged retina scanning radius and CVRF (i.e., BMI, LDL and HDL cholesterol scores, and diabetes mellitus, arterial hypertension, smoking, and physical activity status) [n = 769]. Results shown on MNI152_T1_1mm standard template and glass brain using Mango and python glass-brain packages, neurological view with the left hemisphere on the left side. Coordinates (16, −72, 12) and (−14, −75, 9), respectively. Color bar shows T statistics at **SVC** with FWE at p < 0.05 at cluster level and uncorrected at voxel-level p < 0.001. RNFLT: retinal nerve fibre layer thickness, GMD: grey matter density, TIV: total intracranial volume, CVRF: cardiovascular risk factors, BMI: body-mass index, LDL: low-lipoprotein density, HDL: high-lipoprotein density, MNI: Montreal Neurological Institute, SVC: small volume correction (i.e., FWE correction at p < 0.05 level was applied masking with bilateral occipital cortex instead of whole-brain), FWE: family-wise error.

We informally tested the presence of stronger contralateral associations with an ROI analysis, namely, whether the right nasal RNFLT had stronger correlations than the left nasal RNFLT with the GMD in the left calcarine cortex and whether the left nasal RNFLT had greater correlations than the right nasal RNFLT with the GMD in the right calcarine cortex. Beta values were extracted from the left and right calcarine cortex correlations using Neuromorphometrics atlas in CAT12. Then the left and right nasal RNFLT beta values were compared with paired-sample t-tests for each calcarine cortex [Supplement Tables S5 and S6, Figure 4 (A and B)]. Finally, the left and right calcarine cortices were compared via independent-samples t-tests for each nasal RNFLT [Supplement Tables S7 and S8 Figure 4 (C and D)]. The paired-sample t-tests showed that the right and left nasal RNFLT had significantly greater positive correlations with the GMD in the contralateral calcarine cortices than the ipsilateral counterparts (see the t-test results in the Supplement Table S5 and S6). While the independent-samples t-test for the left nasal RNFLT showed significantly stronger positive correlations with the contralateral calcarine cortex, the independent t-test for the right nasal RNFLT did not reveal a significant contralateral calcarine cortex relationship (See the t-test results in the Supplement Table S7 and S8).

**Figure 4.**
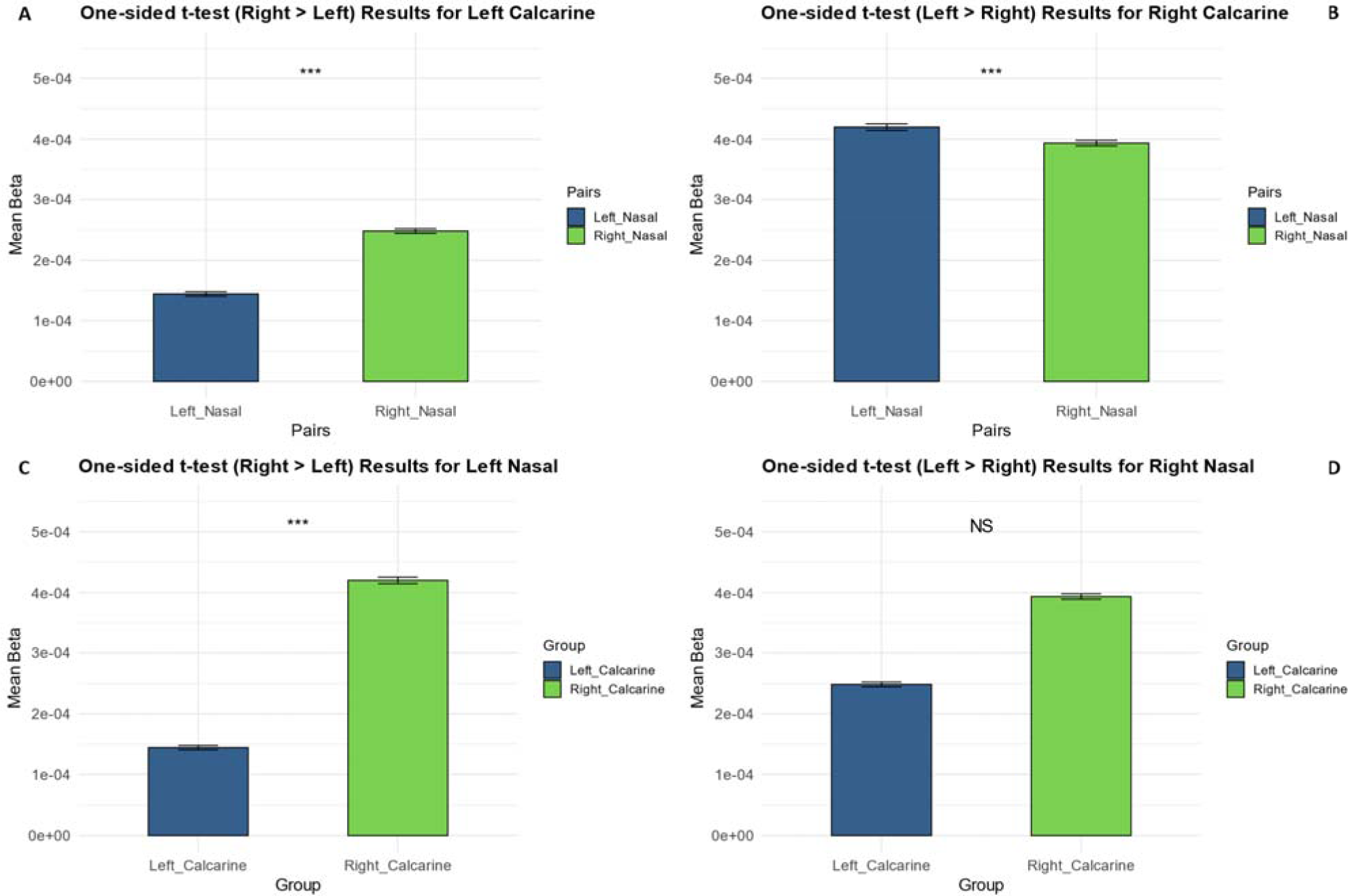
Paired (A and B) and independent (C and D) samples t-test results between the left and right nasal RNFLT correlations with the left and right calcarine cortex GMD. Error bars show standard errors, ***: p < 0.001. RNFLT: retinal nerve fibre layer thickness, GMD: grey matter density.

These results could be explained by possible functional/structural dominance of the right retina over the left and the right calcarine cortex over left calcarine cortex. To test this possible explanation, we additionally performed a paired-sample t-test using the left and right nasal RNFLT for the bilateral (as a whole) calcarine cortex, similar to the global RNFLT comparisons above. Overall, this analysis showed that the right nasal RNFLT had significantly higher positive correlations than the left nasal RNFLT with the bilateral calcarine GMD (Supplementary Figure S18 and Table S9 for the t-test results), supporting the interpretation of right-side dominance.

### 3.4. Correlations between CVRF and brain GMD

Higher BMI and presence of diabetes mellitus and smoking were significantly correlated with lower brain GMD (see Figure 5 and Supplementary Table S10). On the other hand, HDL and LDL cholesterol levels had significant positive correlations with brain GMD (see Supplementary Table S11). We didn’t find any significant correlations between arterial hypertension and physical activity status and brain GMD at p < 0.05 FWE corrected at cluster level and p < 0.001 uncorrected at voxel level. Age, sex, and TIV variables were always controlled for in all multiple regression models.

**Figure 5.**
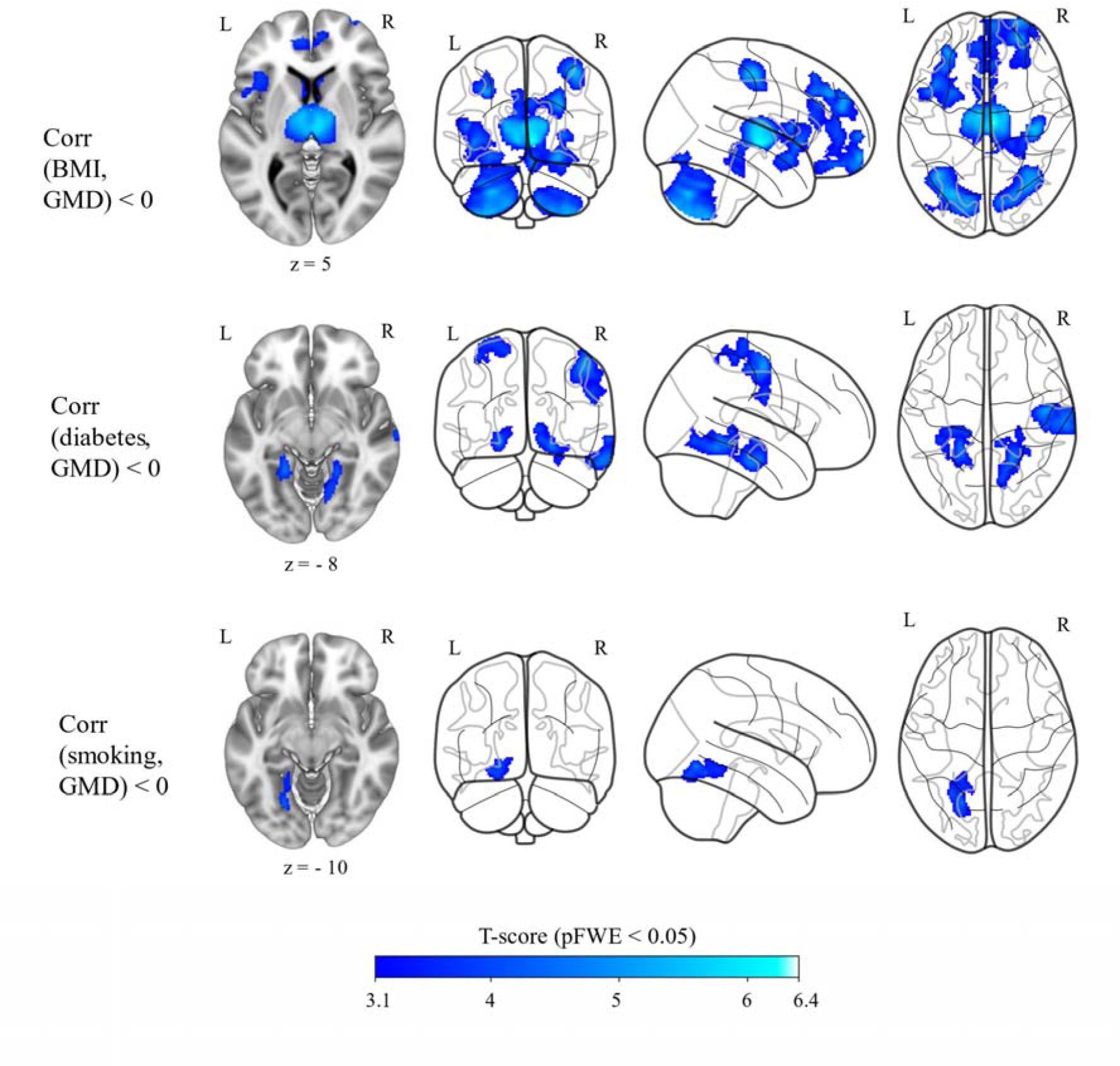
BMI, diabetes mellitus, and smoking status showed negative correlations (blue overlay) with brain GMD when controlling for age, sex, and TIV (n = 769). Results shown on MNI152_T1_1mm standard template and glass brain using Mango and python glass-brain packages, neurological view with the left hemisphere on the left side. Coordinates (15, −18, 5), (−22, −42, −8), and (−22, −64, −10), respectively. Color bar shows T statistics corrected at cluster-level pFWE < 0.05 and uncorrrected at voxel-level p < 0.001. BMI: Body-Mass Index, GMD: grey matter density, TIV: total intracranial volume, MNI: Montreal Neurological Institute, FWE: family-wise error.

### 3.5. Possible overlap between CVRF and RNFLT correlations with brain GMD

The conjunction analysis between correlations of the averaged global RNFLT with the brain GMD and correlations of each CVRF with the brain GMD showed no spatial overlap [Compared correlations were p < 0.05 FWE corrected at cluster level and p < 0.001 uncorrected at voxel level].

### 3.6. Correlations between RNFLT and brain WMM (N = 550 whole-brain TBSS)

#### 3.6.1. Averaged global RNFLT correlations with FA

Averaged global RNFLT had significant positive correlations with FA values, mainly in the bilateral visual pathways (i.e., optic radiation), inferior fronto-occipital fasciculus (IFOF), inferior longitudinal fasciculus (ILF), and the corpus callosum (CC, forceps major)) as but also in the anterior and posterior thalamic radiations (ATR, PTR) (Figure 6 upper, Table 4) when controlling for the CVRF (i.e., BMI, LDL and HDL cholesterol scores, diabetes mellitus, arterial hypertension, smoking, and physical activity status) in addition to age, sex, and averaged retina scanning radius in the multiple regression model using voxel-based whole-brain TBSS. When the CVRF were not included as additional covariates, we also found positive correlations with the bilateral visual pathways (i.e., OR, Figure 6 middle, Supplementary Table S13), but there were some differences compared to the analysis with the CVRF covariates. The differences were found mostly in the right visual pathways. For example, without CVRF covariates, averaged global RNFLT was positively correlated with FA values in the posterior optic radiation (i.e., closer to occipital cortex) while it was positively correlated with FA values in the anterior optic radiation (i.e., closer to the lateral geniculate nucleus (LGN)) with CVRF covariates (Figure 6 lower).

**Figure 6.**
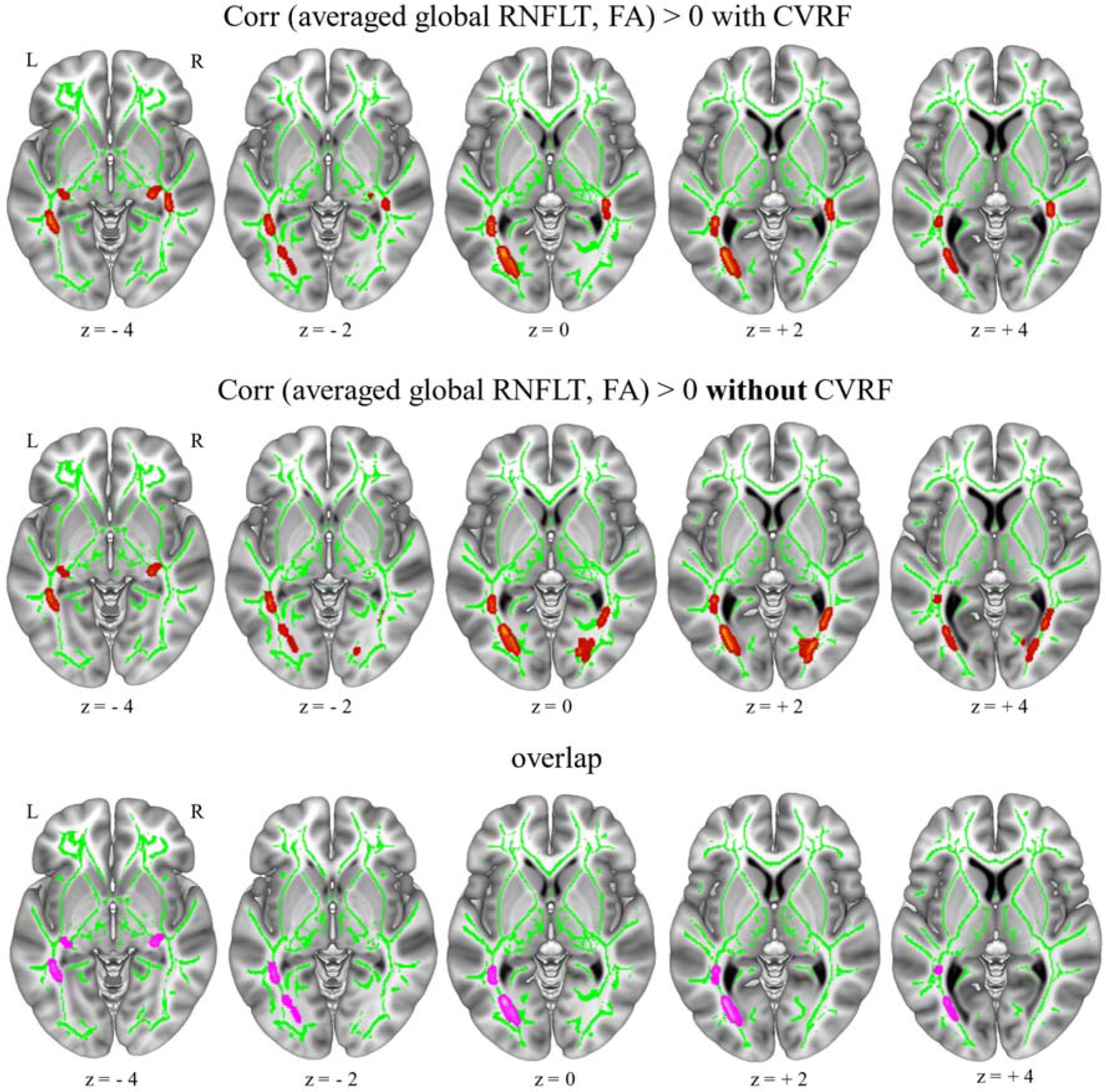
Averaged (between right and left eye per person) global RNFLT showed positive correlations (red overlay) with brain FA with and without controlling for CVRF (n = 550). Upper figure shows the RNFLT correlations when controlling for CVRF (i.e., BMI, LDL and HDL cholesterol scores, and diabetes mellitus, arterial hypertension, smoking, and physical activity status) in addition to age, sex, and averaged retina scanning radius. Middle figure shows the correlations when controlling for only age, sex, and averaged retina scan radius [**i.e., without CVRF].** Lower figure shows the overlap between the correlations of models with and without controlling CVRF (pink). Results shown on MNI152_T1_1mm standard template combined with the cohort’s mean FA skeleton (green) using Mango toolbox, neurological view with the left hemisphere on the left side. Coordinates (−27, −72, 0). Red overlay shows T statistics corrected at cluster-level pFWE < 0.05 and uncorrected at voxel-level p < 0.001, FSL ‘tbss_fill’ was used for visualization purposes. RNFLT: retinal nerve fibre layer thickness, FA: fractional anisotropy, CVRF: cardiovascular risk factors, BMI: body-mass index, LDL: low-lipoprotein density, HDL: high-lipoprotein density, MNI: Montreal Neurological Institute, FWE: family-wise error.

**Table 4.** Whole brain TBSS results: Averaged* global RNFLT correlations with the brain FA (when all CVRF were controlled for)

| Clustersiz | pFWE- | coordinates (mm) |  |  | Tract/Area |
| --- | --- | --- | --- | --- | --- |
|  |  | x | y | z |  |

| e<br>(voxels) | corr |  |  |  |  |
| --- | --- | --- | --- | --- | --- |
| 109 | 0.005 | -38 | -42 | -9 | ILF+IFOF/ Sagittal Stratum [ILF+IFOF]/ OR+CC |
| 108 | 0.005 | -31 | -64 | -1 | IFOF+ILF+*Fmaj*/ PTR[OR]/ OR+CC+Visual Cortex |
| 93 | 0.008 | 34 | -10 | -15 | -/Fornix+Stria Terminalis/ Hippocampus+Amygdala+OR+Fornix+*LGN*+*AcRad* |
| 90 | 0.008 | -30 | -11 | -14 | ILF+ATR/ unc+*Fornix*+*Stria Terminalis*/ OR+Hippocampus+Amygdala+OR |
| 70 | 0.016 | 41 | -36 | -10 | ILF+IFOF/ Sagittal Stratum [ILF+IFOF]/ OR+CC+*Fornix* |
| 45 | 0.043 | 31 | -51 | 13 | *Fmaj*+IFOF/ Tapetum/ CC+OR |
| 43 | 0.049 | 38 | -34 | -1 | IFOF+ILF/ Internal capsule/ OR+CC |

ILF: inferior longitudinal fascicle, IFOF: inferior fronto-occipital fascicle, PTR: posterior thalamic radiation, OR: optic radiation, CC: corpus callosum, LGN: lateral geniculate nucleus, AcRad: acoustic radiation, \*: averaged between right and left eye per person

We additionally tested whether there were signs of laterality (right dominance) regarding the covariance of right and left global RNFLT with FA values. Right and left global RNFLT had significant positive correlations with the FA values in bilateral visual pathways (with and without CVRF as covariates, see Supplement Tables S12-S13). However, the right global RNFLT showed some distinct correlations predominantly in the right optic radiation (Supplement Figure S9) while the left global RNFLT showed some distinct correlations predominantly in the left optic radiation (Supplement Figure S10) when comparisons were made with and without CVRF as covariates in addition to age, sex, and related retina scanning radius variables (see also Supplement Figure S11 for sagittal, coronal, and axial views together).

Interestingly, averaged global RNFLT values were also positively correlated with FA in the limbic pathways (i.e., fornix and stria terminalis including hippocampus and amygdala (independent of using CVRF as covariates)). Notably, right global RNFLT had positive correlations with the FA values of the hippocampus when compared to the left global RNFLT (see the comparisons between the right and left global RNFLT correlations in Supplement Figure S12-S13).

The right global RNFLT seemed to have spatially more extended correlations within bilateral visual tracts than the left global RNFLT correlations (see Supplement Figure S12-S13). To informally test whether the right global RNFLT exhibits significantly stronger correlations with FA compared to the left global RNFLT, we conducted a ROI analysis using the beta values in bilateral optic radiation with a paired-t-test. The results showed that the Right Global RNFLT had significantly higher correlations with the FA in the optic radiation than the Left Global RNFLT (see Supplement Table S14).

#### 3.6.2. Averaged global RNFLT correlations with mean diffusivity (MD)

Averaged global RNFLT had significant negative correlations with the MD values in the left frontal area independent of using CVRF as covariates (Figure 7, Table 5, Supplement Table S15).

**Figure 7.**
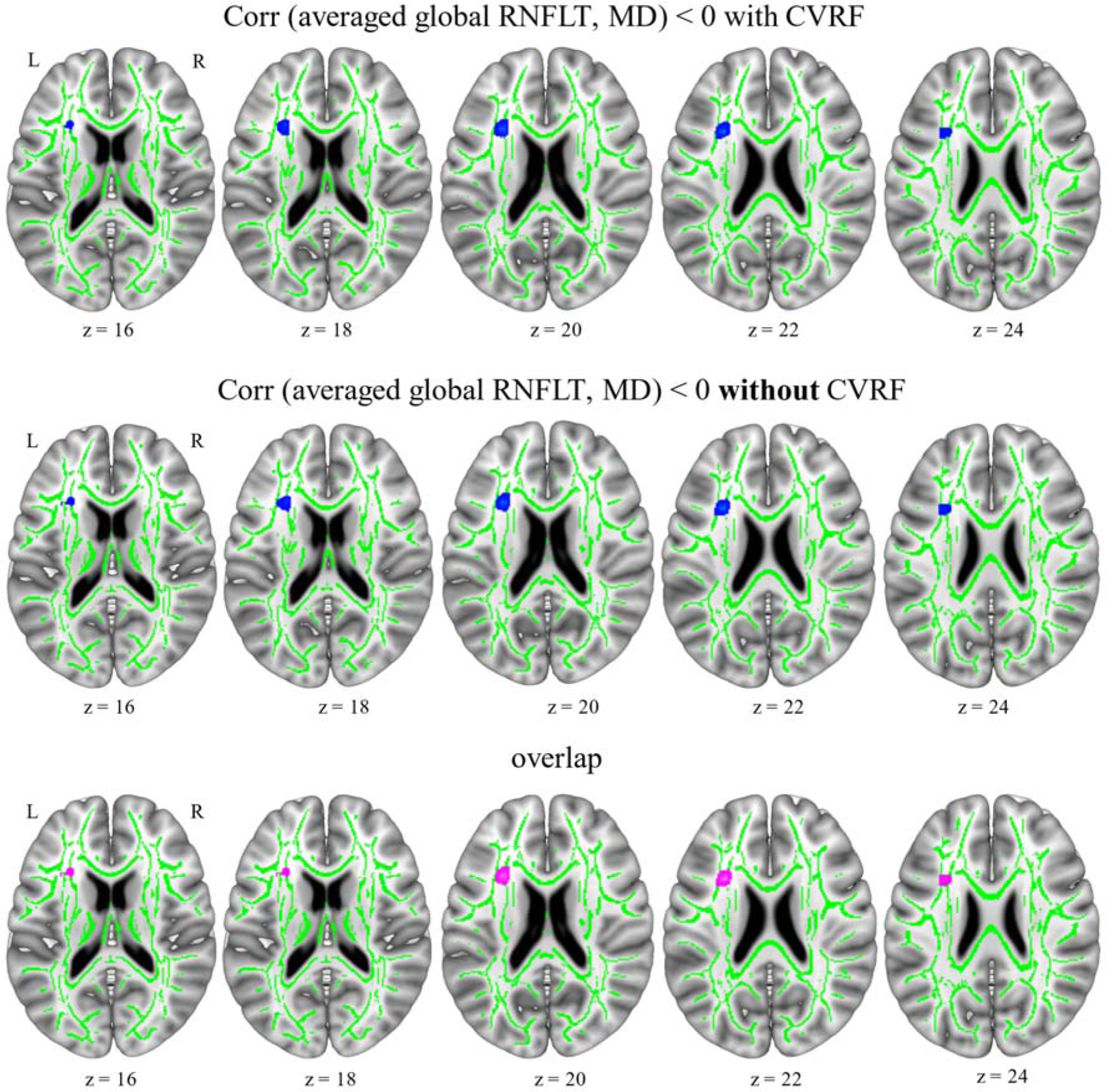
Averaged (between right and left eye per person) global RNFLT showed negative correlations (blue overlay) with brain MD with and without controlling for CVRF (n = 550). Upper figure shows the RNFLT correlations when controlling for CVRF (i.e., BMI, LDL and HDL cholesterol scores, and diabetes mellitus, arterial hypertension, smoking, and physical activity status) in addition to age, sex, and averaged retina scanning radius. Middle figure shows the correlations when controlling for only age, sex, and averaged retina scan radius [**i.e., without CVRF].** Lower figure shows the overlap between the correlations of models with and without controlling CVRF (pink). Results shown on MNI152_T1_1mm standard template combined with the cohort’s mean FA skeleton (green) using Mango toolbox, neurological view with the left hemisphere on the left side. Coordinates (−28, 16, 20). Blue overlay shows T statistics corrected at cluster-level pFWE < 0.05 and uncorrected at voxel-level p < 0.001, FSL ‘tbss_fill’ was used for visualization purposes. RNFLT: retinal nerve fibre layer thickness, MD: mean diffusivity, CVRF: cardiovascular risk factors, BMI: body-mass index, LDL: low-lipoprotein density, HDL: high-lipoprotein density, MNI: Montreal Neurological Institute, FWE: family-wise error.

**Table 5.** Whole brain TBSS results: Averaged*, left and right global RNFLT negative correlations with brain MD.

|  | Cluster | pFWE-corr | coordinates (mm) |  |  | Tract/Area |
| --- | --- | --- | --- | --- | --- | --- |
|  |  |  | x | y | z |  |
| Averaged global RNFLT with MD | 44 | 0.021 | -30 | 18 | 18 | IFG |
| Left global RNFLT with MD without CVRF | 35 | 0.03 | -30 | 18 | 18 | IFG |
|  | 31 | 0.039 | 9 | 4 | -3 | ATR |
| Right global RNFLT with MD without CVRF | 33 | 0.036 | -30 | 18 | 18 | IFG |
|  | 30 | 0.044 | -19 | -66 | 32 | SPL/Precuneus/LOCC |

IFG: inferior frontal gyrus, ATR: anterior thalamic radiation, SPL: superior parietal lobule, LOCC: lateral occipital cortex, CVRF: cardiovascular risk factors, \*: between right and left eye per person.

However, there were some differences for the left and the right global RNFLT negative correlations with MD when accounting for CVRF versus when not. Specifically, the left global RNFLT showed negative correlations with MD in the right ATR (Supplement Figure S14-S17, pink-mid-lower) while the right global RNFLT showed negative correlations with MD in the left superior parietal lobe, SPL, (Supplement Figure S15, pink-mid-lower; Figure S17, blue-top-lower) when CVRF were not controlled, along with age, sex, and respective retina scanning radius variables (Table 5, Supplement Table S16). The left and right global RNFLT both showed negative correlations with MD values in the left inferior frontal gyrus, independent of using CVRF as covariates (Figure S14-S17, bottom, overlap shown in pink). The overlapping correlations seemed to remain spatially similar regardless of whether CVRF was accounted for or not.

#### 3.6.3. Nasal RNFLT correlations with brain WMM

We found no significant correlations at p < 0.05 FWE-corrected at cluster level and p < 0.001 uncorrected at voxel level of the left and right nasal RNFLT, with either FA or MD values, when we controlled only for age, sex, and related retina scanning radius variables, nor when additionally including CVRF.

### 3.7. Correlations between CVRF and brain WMM parameters FA and MD

Arterial hypertension and smoking status had significant negative correlations with brain FA values in the left parietal regions (Figure 8 and see Supplement Table S17). There were no significant correlations between BMI, HDL, LDL scores, or diabetes mellitus and physical activity status and brain FA values when controlling for age and sex. Arterial hypertension and diabetes mellitus status had significant positive correlations with brain MD values in the body of the corpus callosum (CC) and superior longitudinal fasciculus (SLF), respectively (Figure 8 and Supplementary Table S18). BMI values had significant positive correlations with the MD of the forceps minor of the CC and negative correlations with the bilateral visual and cortico-spinal tract MD values (Figure 8 and Supplementary Table S19). There were no significant correlations of HDL and LDL scores, physical activity, or smoking status with MD values when controlling for age and sex.

**Figure 8.**
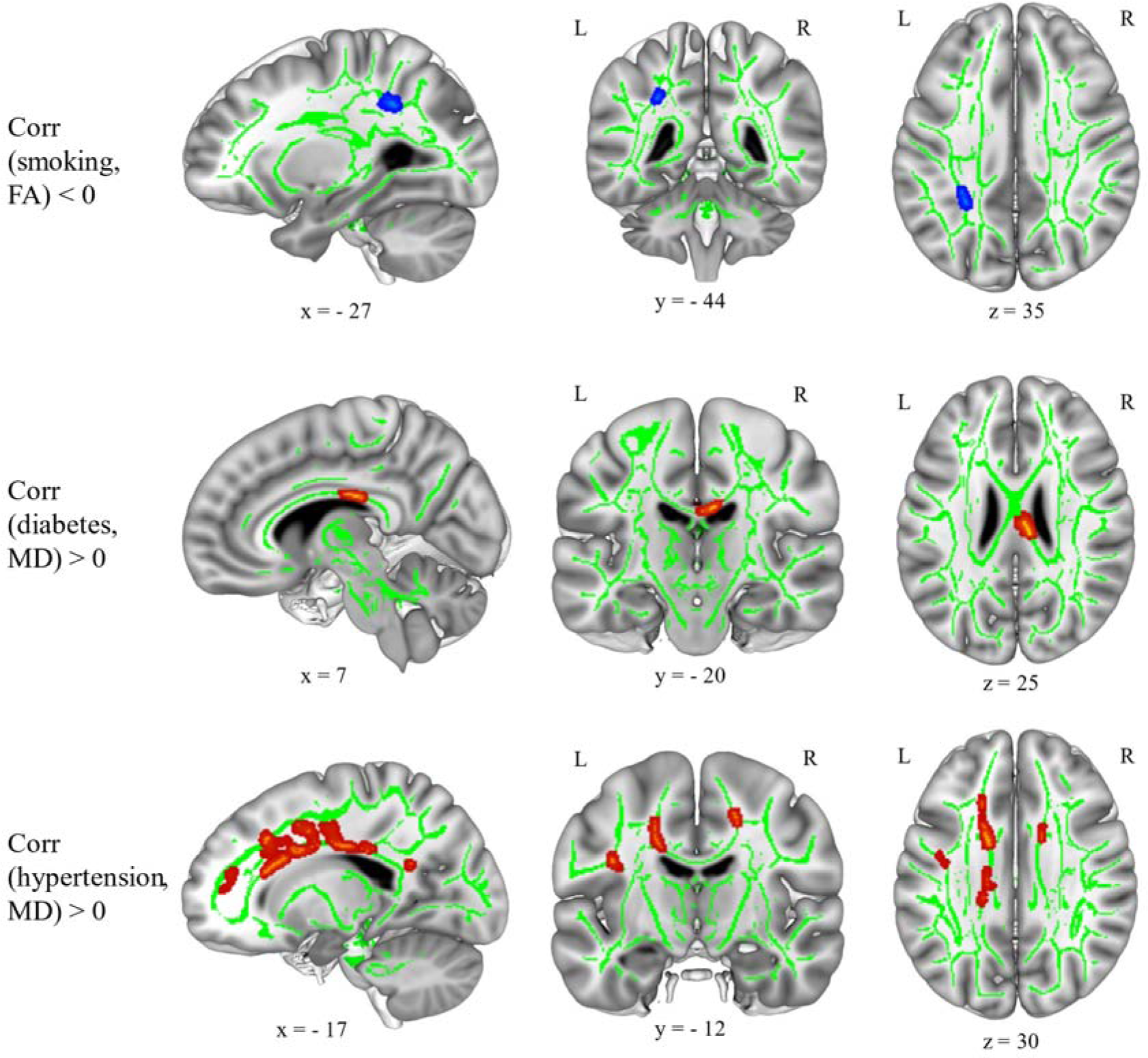
Smoking status showed negative correlations (blue overlay) with brain FA; diabetes mellitus and arterial hypertension status showed positive correlations (red overlay) with brain MD, when controlling for age and sex (n = 550). Results shown on MNI152_T1_1mm standard template combined with the cohort’s mean FA skeleton (green) using Mango toolbox, neurological view with the left hemisphere on the left side. Coordinates (−28, 16, 20). Blue and red overlays show T statistics corrected at cluster-level pFWE < 0.05 and uncorrected at voxel-level p < 0.001, FSL ‘tbss_fill’ was used for visualization purposes. FA: fractional anisotropy, MD: mean diffusivity, MNI: Montreal Neurological Institute, FWE: family-wise error.

### 3.8. Possible overlap between CVRF and RNFLT correlations with Brain WMM

There were no spatial conjunctions between correlations of the averaged global RNFLT and each CVRF with brain FA and MD values [Compared correlations were p < 0.05 FWE corrected at cluster level and p < 0.001 uncorrected at voxel level].

### 3.9. ROI analyses

Separately from the whole-brain voxel-wise analyses, we had confirmational hypotheses based on prior ROIs defined from the literature (Supplement Table S20). Partial correlation analyses were performed between averaged global RNFLT and regional grey matter volume (obtained via CAT12 atlases) measures using age, sex, TIV, and retina scanning radius as control variables. Partial variance of the CVRF was investigated by adding them as additional covariates into the analysis. There were significant positive partial correlations of the averaged global RNFLT with bilateral calcarine cortices, independent of using CVRF as covariates after FDR correction for multiple comparisons (for the number of ROIs (Supplement Table S20)). Additionally, there were significant positive partial correlations of the RNFLT with the whole brain white matter and the left cerebellar volumes after FDR correction and controlling for CVRF. If the FDR correction threshold was waved, whole brain white matter, bilateral cerebellum, bilateral calcarine cortices, left fusiform (when controlling for CVRF), right lingual, left middle cingulate, left occipital pole, and bilateral para-hippocampal gyri were found to be significantly correlated with averaged global RNFLT, both with and without taking CVRF into account (see Supplement Table S20).

## DISCUSSION

Using cross-sectional retinal optical coherence tomography (OCT) and brain magnetic resonance imaging (MRI) data, from a population-based study, we tested two pre-registered hypotheses proposing structural covariations of circumpapillary retinal nerve fibre layer thickness (RNFLT) with: (i) visual pathways and brain areas and (ii) more widespread brain areas related to the influence of cardiovascular risk factors (CVRF). Our results provide general support for the first hypothesis and no evidence for the second. However, we found some evidence for a relationship between RNFLT and brain structure in brain areas possibly of relevance to subclinical Alzheimer’s pathologies.

We clearly demonstrated a structural covariation of global RNFLT with gray matter density (GMD) in primary visual cortices (Brodmann area 17) and white matter microstructure (WMM) properties in optic radiata using whole-brain voxel-wise analyses. This is in line with the results of the Rotterdam and Hisayama studies (Mutlu et al., 2018; Ueda et al., 2022). A novel finding of our work was that left and right nasal RNFLTs partially confirmed structural covariation with the grey matter of contralateral primary visual cortices, albeit with a weaker signal-to-noise ratio, as expected. Interestingly, the structural correlations of the nasal sector RNFLTs with the brain structure were stronger for the right eye than for the left eye, consistent with a possible right-eye dominance in the LIFE-Adult-Study cohort. Eye-dominancy has been suggested to continue in ocular dominance columns (Ip et al., 2021; Ooi & He, 2020), and right-eye dominancy has been observed in previous studies marked by a thicker retinal nerve fibre layer in the right eye and functional dominance of the right visual cortex measured by more cell bodies (Amunts et al., 2007; Baniasadi et al., 2020; Cameron et al., 2017; Choi et al., 2014; Hougaard et al., 2015).

Growing evidence has shown that CVRF have similar effects on retina and brain, i.e., retinal nerve fibre layer thinning (Colijn et al., 2019; Langner et al., 2022; Majithia et al., 2022; Mauschitz et al., 2018; Rauscher et al., 2024; Rauscher et al., 2021) and brain grey and white matter reduction (Beyer, Kharabian Masouleh, et al., 2019; Ebrahimi et al., 2023; Kharabian Masouleh et al., 2016; Schaare et al., 2019; Zhang et al., 2018). Moreover, some studies suggested that neurodegeneration in the retina and the brain might both be related to vascular damage, as an underlying shared pathology (Mirzaei et al., 2020; Ramirez et al., 2017; Ravi Teja et al., 2017; Santos et al., 2017; Shin et al., 2021). Using macular RNFLT of the UK-Biobank longitudinal data over roughly an 8-year period, a recent study suggested retinal thinning as an additional biomarker for developing cardiovascular disease (Chen et al., 2023). In our study, however, we could not provide further evidence regarding the possibility of RNFLT mirroring the impact of CVRF on the brain. While we generally confirmed that CVRF were associated with lower grey and white matter in several brain regions, we did not show additional significant CVRF associations with RNFLT. Furthermore, a conjunction analysis between CVRF and RNFLT covariances on the brain revealed no overlap.

On the other hand, simultaneous inclusion of all CVRF in the models, as done in earlier studies (Chua et al., 2021; Ong et al., 2015), somewhat impacted the RNFLT covariations with the brain grey and white matter. Averaged global RNFLT showed significant correlations with FA in the anterior parts of the optic radiation (close to LGN, with CVRF controlled for) and with the posterior parts of the optic radiation (close to calcarine cortex, without CVRF). This was especially true in the right hemisphere. When examining left and right global RNFLT separately, with CVRF as a covariate, we additionally found spatially more extended FA correlations in ipsilateral optic radiations. These correlations, although relatively small, may be related to the vascular supply (i.e., anterior-posterior circulation) of the optic radiation and/or to right eye dominance related to fiber allocation within the pathways. Furthermore, while averaged global RNFLT showed significant correlations with MD in the left inferior frontal region, regardless of correction for CVRF, there were additional correlations of left and right global RNFLT with MD values of small clusters in the right anterior thalamus and left superior parietal lobe, respectively, when not correcting for CVRF. As a whole, our results suggest there is not enough evidence showing significant variance shared between CVRF and RNFLT on brain structure in our sample. However, this does not rule out the possibility that the retina and brain are similarly affected by neurodegenerative processes with or without the effect of CVRF.

In addition to visual pathways, in our whole-brain WMM analysis, we found associations specifically related to memory- and emotion-related limbic pathways i.e., the fornix and stria terminalis including the hippocampus and amygdala similar to several studies (Barrett-Young et al., 2023; Chua et al., 2021; Mauschitz et al., 2022; Mutlu et al., 2017; Ong et al., 2015; van der Heide et al., 2024). Additionally, meta-analyses and (systematic) reviews have consistently shown retinal nerve fibre layer thinning in Alzheimer’s disease patients (Chan et al., 2019; Coppola et al., 2015; Ge et al., 2021; Thomson et al., 2015). On a cellular level this might be explained by potential retinal ganglion cell and nerve fibre loss through the optic nerve relating to the visual sensation of the information processing in patients with Alzheimer’s disease, compared to non-demented healthy controls (Hinton et al., 1986; Sadun & Bassi, 1990). Thus, we interpret these findings as potential presence of subclinical pathologies related to the development of Alzheimer’s disease in our elderly-weighted sample (Alber et al., 2020; Gaire et al., 2024).

The main **limitation** of our study is its cross-sectional nature. Furthermore, the overall generalizability is limited as the study’s data was drawn from a relatively healthy European population. Larger sample sizes combining multicenter studies to represent balanced subsamples of CVRF, or meta-analyses, will be needed to demonstrate more robust results. The study’s **strengths,** on the other hand, include the availability of high-quality OCT and MRI data in the same subjects at the same approximate time, as well as objective CVRF measures, rigorous statistics, and preregistration. **In conclusion,** our data provide strong evidence that retinal thickness shows structural covariations with associated brain areas, i.e., visual projections and visual cortices, as well as areas beyond, such as limbic pathways. Furthermore, although we could not confirm that RNFLT can reflect the impact of CVRF on the brain, our results suggest that the retina may reflect some neurodegenerative changes in the brain. Longitudinal studies with intra-individual trajectories would allow the effects of various influencing factors such as age, CVRF, neurodegeneration, and others to be disentangled. They would further allow full advantage to be taken of the insights that can be gained from optical access to the retina, an easily accessible part of the brain.

## Data and Code Availability

We used the data obtained within the LIFE-Adult-Study (Engel et al., 2022; Loeffler et al., 2015).

The scripts used for the analyses and visualization of the results can be found here: https://github.com/n-ayyildiz/EyeBrain_project.git

## Author Contributions

Conceptualization: NA, FGR, AV

Data curation: NA, FB, CE, RB, KW, SZ, JG, TE, MW, AVW, FGR, AV

Formal analysis: NA, discussed with [KM, SH, FB, JDH, AA, AAW, FGR, AV], JDH (ROI-based analysis regarding Supplement S20)

Funding acquisition: FGR, AV

Investigation: NA, FB, CE, RB, KW, SZ, JG, TE, MW, AVW, FGR, AV

Methodology: NA, KM, SH, FB, CE, RB, KW, SZ, JG, AA, TE, MW, AVW, FGR, AV

Project administration: NA, FGR, AV

Resources: AVW, FGR, AV

Software: NA, KM, SH, AA

Supervision: NA, FGR, AV

Validation: NA, JDH

Visualization: NA, KM, AA

Writing – original draft: NA, FGR, AV

Writing – review & editing: NA, KM, SH, FB, CE, RB, KW, SZ, JG, JDH, AA, TE, MW, AVW, FGR, AV

## Acknowledgements

The authors would like to thank all researchers and study staff of the LIFE-Adult-Study for their time and efforts ranging from study design to data acquisition. We specifically extend our sincere thanks to all study participants for their time. The authors are grateful to Dr. Joshua Grant from the Max Planck Institute for Human Cognitive and Brain Sciences for his valuable recommendations and proofreading of the manuscript.

Preregistration of the study’s hypotheses, relevant methods, and analysis plan can be found here, ‘eye-brain VBM project’, at Open Science Forum platform:

https://osf.io/mdgq3/registrations (https://doi.org/10.17605/OSF.IO/MDGQ3)

A poster with some of the results of this paper was presented at FENS 2024. The poster can be found here: DOI: 10.17605/OSF.IO/9EWV6

