## supplemental figures and tables for "Retinal nerve fibre layer thickness reflects characteristics of brain grey and white matter": 0_EyeBrain_Supplement_Tables_Nazife_Ayyildiz_8May2025.pdf

### CORRELATIONS

Table S1. Partial Correlations between Average Global RNFLT and CVR factors when controlling for the Average Retina Scan Radius [n=769 (VBM Sample)]

| | Age | Sex | Diabetes Status | BMI | Hypertension Status | IPAQ Category | LDL | HDL | Smoking Status |
|---|---|---|---|---|---|---|---|---|---|
| Average Global RNFLT | **-0,27**** | -0,11 | **-0,12**** | -0,06 | **-0,1*** | -0,04 | -0,06 | 0,05 | 0,02 |
| Age | | 0,07 | **0,41**** | **0,31**** | **0,4**** | **0,09*** | **0,31**** | 0,03 | **-0,12**** |
| Sex | | | **0,14**** | **0,09*** | **0,08*** | 0,04 | **0,07*** | **-0,46**** | **0,13**** |
| Diabetes Status | | | | **0,37**** | **0,34**** | 0,02 | 0,02 | **-0,13**** | -0,02 |
| BMI | | | | | **0,38**** | -0,02 | **0,14**** | **-0,35**** | 0,07 |
| Hypertension Status | | | | | | 0,06 | 0,04 | **-0,16**** | **-0,09*** |
| IPAQ Category | | | | | | | 0,01 | 0,04 | **-0,1*** |
| LDL | | | | | | | | **-0,11*** | 0,03 |
| HDL | | | | | | | | | **-0,11*** |

**: p<=0.001, *: p<0.05

Table S2. Partial Correlations between Average Global RNFLT and CVR factors when controlling for the Average Retina Scan Radius [n=550 (TBSS Sample)]

| | Age | Sex | Diabetes Status | Hypertension Status | BMI | Smoking Status | IPAQ Category | HDL | LDL |
|---|---|---|---|---|---|---|---|---|---|
| Average Global RNFLT | **-0,28**** | -0,07 | **-0,15**** | **-0,11*** | -0,02 | **0,08*** | -0,04 | 0,003 | **-0,11*** |
| Age | | 0,04 | **0,39**** | **0,39**** | **0,31**** | **-0,16**** | **0,1*** | 0,04 | **0,37**** |
| Sex | | | **0,14**** | 0,02 | 0,06 | **0,13*** | 0,04 | **-0,47**** | **0,1*** |
| Diabetes Status | | | | **0,32**** | **0,33**** | -0,02 | 0,04 | **-0,13*** | **0,09*** |
| Hypertension Status | | | | | **0,34**** | **-0,09*** | 0,07 | **-0,11*** | 0,08 |
| BMI | | | | | | **0,09*** | -0,04 | **-0,32**** | **0,2**** |
| Smoking Status | | | | | | | -0,08 | **-0,14**** | 0,02 |
| IPAQ Category | | | | | | | | 0,04 | 0,01 |
| HDL | | | | | | | | | **-0,15**** |

**: p<=0.001, *: p<0.05

**RESULTS: VBM**

Table S3. Whole Brain VBM Results: Average, Left and Right Global Mean RNFLT positive correlations with the brain gray matter density (when only age, sex, TIV and retina scan radius controlled)

| RNFLT | | | | | coordinates mm | | | |
|---|---|---|---|---|---|---|---|---|
| | cluster p(FWE-corr) | cluster size | peak T | peak p(unc) | x | y | z | Area* |
| Average Global | 0,008 | 1421 | 6,07 | 0,000 | 16 | -80 | 14 | R Calcarine Cortex/ V1, 17 CalcS, hOc1 |
| | | | 5,3 | 0,000 | 18 | -90 | 4 | |
| | 0,009 | 1358 | 6 | 0,000 | -15 | -78 | 8 | L Calcarine Cortex/ V1, 17 CalcS, hOc1 |
| | | | 5,15 | 0,000 | -16 | -94 | 0 | |
| Left Global | 0,019 | 1137 | 5,67 | 0,000 | 16 | -78 | 14 | R Calcarine Cortex/ V1, 17 CalcS, hOc1 |
| | | | 5,05 | 0,000 | 18 | -90 | 4 | |
| | 0,038 | 936 | 5,53 | 0,000 | -15 | -78 | 8 | L Calcarine Cortex/ V1, 17 CalcS, hOc1 |
| | | | 4,63 | 0,000 | -16 | -94 | 0 | |
| Right Global | 0,006 | 1500 | 6,02 | 0,000 | 16 | -80 | 14 | R Calcarine Cortex/ V1, 17 CalcS, hOc1 |
| | | | 5,75 | 0,000 | 16 | -70 | 14 | |
| | | | 5,2 | 0,000 | 20 | -90 | 3 | |
| | 0,004 | 1666 | 5,99 | 0,000 | -15 | -80 | 8 | L Calcarine Cortex/ V1, 17 CalcS, hOc1 |
| | | | 5,32 | 0,000 | -16 | -94 | 0 | |
| | | | 3,75 | 0,000 | -6 | -88 | 4 | |

*: CAT12 Neuromorphometrics Atlas

| | | | | | coordinates mm | | |
|---|---|---|---|---|---|---|---|
| | voxel p(FWE-corr) | cluster size | peak T | peak p(unc) | x | y | z |
| Left Nasal | 0,022 | 15 | 4,76 | 0 | 18 | -75 | 14 |
| Right Nasal | 0,01 | 32 | 4,94 | 0 | -15 | -78 | 8 |
| | 0,028 | 34 | 4,69 | 0 | 15 | -72 | 10 |

### T-test Findings

Table S4. Paired-t test results between the Right Global RNFLT correlations with the GMD and the Left Global RNFLT correlations with the GMD in Bilateral Calcarine Cortex

| Mean difference (Right-Left) | T-statistic* | p-value | Degrees of Freedom | %95 CI-low | %95 CI-high | Cohen's_d |
|---|---|---|---|---|---|---|
| 0.0000219 | 24.69 | 1.83e-130 | 8545 | 0.0000205 | inf | 0.27 |

* One sided paired-t-test, Alternative H: The Right RNFLT correlations are greater than the Left RNFLT correlations, RNFLT correlations from the whole-brain VBM when all CVR factors were controlled for the bilateral Calcarine ROI taken from the Neuromorphometrics Atlas.

Table S5. Paired-t test results between the Left Nasal RNFLT correlations and the Right Nasal RNFLT correlations with the GMD in the **Left Calcarine Cortex**

| Mean difference (Left_N_leftCalc-Right_N_leftCalc) | T-statistic* | p-value | Degrees of Freedom | %95 CI-low | %95 CI-high | Cohen's_d |
|---|---|---|---|---|---|---|
| -0.000104376 | -121.87 | < 2.2e-16 | 4927 | -Inf | -0.000102967 | -0.8043228 |

* One sided paired-t-test, Alternative H: The Left Nasal RNFLT correlations are less than the Right Nasal RNFLT correlations with the Left Calcarine GMD, RNFLT correlations from the whole-brain VBM when all CVR factors were controlled.

Table S6. Paired-t test results between the Left Nasal RNFLT correlations and the Right Nasal RNFLT correlations with the GMD in the **Right Calcarine Cortex**

| Mean difference (Left_N_rightCalc-Right_N_rightCalc) | T-statistic* | p-value | Degrees of Freedom | %95 CI-low | %95 CI-high | Cohen's_d |
|---|---|---|---|---|---|---|
| 2.681466e-05 | 16.943 | < 2.2e-16 | 3617 | 2.421083e-05 | Inf | 0.1726788 |

* One sided paired-t-test, Alternative H: The Left Nasal RNFLT correlations are greater than the Right Nasal RNFLT correlations with the Right Calcarine GMD, RNFLT correlations from the whole-brain VBM when all CVR factors were controlled.

Table S7. Independent-sample t-test results between the **Left Nasal** RNFLT correlations with the GMD in the left and the Right Calcarine Cortex

| Mean | Mean | T- | Degree | p-value | %9 | %95 CI-high | Cohen's_ |
|---|---|---|---|---|---|---|---|

| Left_N_leftCalc | Left_N_rightCalc | statistic* | s of Freedom | | 5 CI-low | | d |
|---|---|---|---|---|---|---|---|
| 0.0001442099 | 0.0004201770 | -85.634 | 6438.1 | < 2.2e-16 | -Inf | -0.0002706656 | -1.955888 |

* One sided independent-t-test (Welch’s Two Sample T-test), Alternative H: The Left Nasal RNFLT correlations with the left Calcarine GMD are less than the Left Nasal RNFLT correlations with the right Calcarine GMD, RNFLT correlations from the whole-brain VBM when all CVR factors were controlled.

Table S8. Independent-sample t-test results between the **Right Nasal** RNFLT correlations with the GMD in the left and the Right Calcarine Cortex

| Mean Right_N_leftCalc | Mean Right_N_rightCalc | T-statistic* | Degrees of Freedom | p-value | %95 CI-low | %95 CI-high | Cohen's_d |
|---|---|---|---|---|---|---|---|
| 0.0002485859 | 0.0003933624 | -48.683 | 7639.4 | 1 | -0.0001496686 | Inf | -1.071.475 |

* One sided independent-t-test (Welch’s Two Sample T-test), Alternative H: The Right Nasal RNFLT correlations with the left Calcarine GMD are greater than the Right Nasal RNFLT correlations with the right Calcarine GMD, RNFLT correlations from the whole-brain VBM when all CVR factors were controlled.

Table S9. Paired-t test results between the Left Nasal RNFLT correlations and the Right Nasal RNFLT correlations with the GMD in the **Bilateral Calcarine Cortex**

| Mean difference (Left_N_bilCalc-Right_N_bilCalc) | T-statistic* | p-value | Degrees of Freedom | %95 CI-low | %95 CI-high | Cohen's_d |
|---|---|---|---|---|---|---|
| -4.883567e-05 | -44.873 | < 2.2e-16 | 8545 | -Inf | -0.0000470 | -0.2544713 |

* One sided paired-t-test, Alternative H: The Left Nasal RNFLT correlations are less than the Right Nasal RNFLT correlations with the Bilateral Calcarine Cortex GMD, RNFLT correlations from the whole-brain VBM when all CVR factors were controlled.

### CVRF: VBM Findings

Table S10. Whole Brain VBM Results: **BMI, Diabetes and Smoking** negative correlations with the brain gray matter density (when age, sex and TIV controlled for)

| | | | | | | coordinates (mm) | |
|---|---|---|---|---|---|---|---|
| | cluster p(FWE-corr) | cluster size | peak T | peak p(unc) | x | y | z |
| BMI | 0 | 6794 | 6,29 | 0 | 6 | -15 | -2 |
| | | | 5,73 | 0 | -6 | -10 | 0 |
| | | | 3,99 | 0 | 21 | -32 | -14 |
| | 0,007 | 1467 | 5,51 | 0 | 39 | -20 | 54 |
| | | | 3,91 | 0 | 30 | -30 | 56 |
| | 0 | 7070 | 5,47 | 0 | -27 | -72 | -44 |
| | | | 5,43 | 0 | -39 | -58 | -63 |
| | | | 5,32 | 0 | -30 | -74 | -60 |
| | 0,016 | 1183 | 5,19 | 0 | 28 | 52 | 27 |
| | | | 3,35 | 0 | 10 | 58 | 28 |
| | 0 | 5000 | 5,16 | 0 | 27 | 68 | -15 |
| | | | 4,65 | 0 | 32 | 66 | 3 |
| | | | 4,65 | 0 | 6 | 64 | -20 |
| | 0 | 3057 | 5,16 | 0 | 34 | -64 | -62 |
| | | | 4,91 | 0 | 22 | -75 | -58 |
| | | | 4,56 | 0 | 12 | -80 | -52 |
| | 0,039 | 926 | 5,11 | 0 | -32 | 42 | 38 |
| | | | 4,54 | 0 | -33 | 30 | 40 |
| | | | 4,15 | 0 | -30 | 30 | 51 |
| | 0 | 2598 | 4,69 | 0 | -40 | 16 | -3 |
| | | | 4,66 | 0 | -46 | 10 | 10 |
| | | | 4,48 | 0 | -27 | 9 | -14 |
| Diabetes | 0,001 | 2201 | 5,07 | 0 | 48 | -12 | 54 |
| | | | 4,24 | 0 | 50 | -22 | 51 |
| | | | 3,83 | 0 | 54 | -10 | 32 |
| | 0,005 | 1577 | 4,64 | 0 | 21 | -50 | -12 |
| | | | 4,08 | 0 | 14 | -68 | -6 |
| | | | 4,05 | 0 | 16 | -60 | -9 |
| | 0,003 | 1723 | 4,63 | 0 | 63 | -21 | -24 |
| | | | 4,08 | 0 | 69 | -18 | -6 |
| | 0,043 | 900 | 4,58 | 0 | -20 | -45 | -8 |
| | | | 3,28 | 0,001 | -14 | -38 | 2 |
| | 0,034 | 966 | 3,94 | 0 | -28 | -27 | 74 |
| | | | 3,82 | 0 | -20 | -27 | 68 |
| | | | 3,82 | 0 | -38 | -34 | 60 |
| Smoking | 0,023 | 1086 | 4,57 | 0 | -21 | -66 | -8 |
| | | | 4,22 | 0 | -28 | -60 | -14 |
| | | | 4,05 | 0 | -22 | -72 | -15 |

Table S11. Whole Brain VBM Results: **HDL and LDL cholesterol** positive correlations with the brain gray matter density (when age, sex and TIV controlled for)

| | | | | | | coordinates (mm) | |
|---|---|---|---|---|---|---|---|
| | cluster p(FWE-corr) | cluster size | peak T | peak p(unc) | x | y | z |
| HDL | 0,005 | 1539 | 4,55 | 0 | -46 | -20 | -38 |
| | | | 4,3 | 0 | -58 | -15 | -40 |
| | | | 4,07 | 0 | -40 | -9 | -48 |
| | 0,001 | 2336 | 4,41 | 0 | 34 | -62 | -54 |
| | | | 4,12 | 0 | 27 | -74 | -58 |
| | | | 3,77 | 0 | 33 | -69 | -46 |
| | 0,003 | 1690 | 3,87 | 0 | -21 | -76 | -46 |
| | | | 3,86 | 0 | -18 | -86 | -51 |
| | | | 3,38 | 0 | -27 | -84 | -34 |
| LDL | 0,002 | 1898 | 5,67 | 0 | 32 | -87 | 20 |
| | | | 4,17 | 0 | 28 | -88 | 0 |
| | | | 3,71 | 0 | 28 | -98 | 15 |
| | 0,044 | 893 | 4,46 | 0 | 38 | 38 | -9 |
| | | | 3,48 | 0 | 20 | 39 | -16 |
| | 0 | 2951 | 4,43 | 0 | -39 | -21 | 3 |
| | | | 4,18 | 0 | -20 | -60 | 0 |
| | | | 4 | 0 | -22 | -22 | -10 |
| | 0,002 | 1935 | 4,42 | 0 | 0 | -51 | 28 |
| | | | 3,91 | 0 | 10 | -32 | 34 |
| | | | 3,44 | 0 | 2 | -24 | 28 |
| | 0,015 | 1204 | 4,32 | 0 | -68 | -26 | -6 |
| | | | 4 | 0 | -64 | -34 | 0 |
| | | | 3,37 | 0 | -57 | -46 | -2 |

### RESULTS: TBSS

Table S12. Whole Brain TBSS Results: Left and Right Global Mean RNFLT positive correlations with the brain **Fractional Anisotropy** (**with** CVRF controlling)

| | | | coordinates | | |
|---|---|---|---|---|---|
| | Cluster Size | pFWE -corr | X (mm) | Y (mm) | Z (mm) |

| | | | | | | |
|---|---|---|---|---|---|---|
| Left | 108 | 0,005 | -29 | -63 | -1 | IFOF+ILF+*Fmaj*/ PTR[OR]/ OR+CC+Visual Cortex |
| | 80 | 0,012 | -30 | -11 | -14 | ILF+*ATR+IFOF*/ unc+*Fornix+Stria Terminalis*/ *OR*+Amygdala+Hippocampus+OR |
| | 78 | 0,012 | 34 | -14 | -13 | ILF/ Fornix+Stria Terminalis/ *OR*+Hippocampus+OR+Fornix+*LGN+AcRad* |
| | 49 | 0,039 | 18 | -80 | -1 | IFOF+ILF+FMaj/ unc/ OR+Visual Cortex |
| Right | 140 | 0,002 | -38 | -44 | -8 | ILF+IFOF/ Sagittal Stratum[ILF+IFOF]+*PTR[OR]*/ OR+CC |
| | 107 | 0,004 | -31 | -64 | -1 | IFOF+ILF/ PTR[OR]+*Sagittal Stratum[ILF+IFOF]/* OR+CC+Visual Cortex |
| | 106 | 0,004 | 40 | -36 | -11 | ILF+IFOF/ Sagittal Stratum[ILF+IFOF]/ OR+CC |
| | 100 | 0,004 | -32 | -13 | -14 | -+*ATR*/Fornix+Stria Terminalis/ *OR*+Hippocampus+OR+Amygdala |
| | 90 | 0,006 | 34 | -10 | -15 | -/Fornix+Stria Terminalis/ Hippocampus+Amygdala+OR+Fornix+*AcRad* |
| | 82 | 0,009 | 31 | -51 | 13 | Fmaj+IFOF/ Tapetum/ CC |
| | 74 | 0,011 | 25 | -37 | -8 | Cingulum (hippocampus)/ unc+*Cingulum (hippocampus)*/ Hippocampus+OR+CC+*Cingulum* |

Table S13. Whole Brain TBSS Results: Average, Left and Right Global Mean RNFLT positive correlations with the brain **Fractional Anisotropy** (**without** CVRF controlling)

| | | | coordinates | | | |
|---|---|---|---|---|---|---|
| RNFLT | Cluster size | pFWE -corr | x | y | z | Tract/ Area |
| avG_woCVR | 104 | 0,006 | -31 | -64 | -1 | IFOF+ILF+*Fmaj*/ PTR[OR]/ OR+CC+Visual Cortex |
| | 90 | 0,009 | 34 | -10 | -15 | -/ Fornix+Stria Terminalis/ *OR*+**Hippocampus**+Amydala+OR+Fornix+*LGN+AcRad* |
| | 86 | 0,01 | -34 | -15 | -13 | IFOF+*ATR*/ unc+*Fornix+Stria Terminalis*/ *OR*+OR+IFOF |
| | 79 | 0,012 | -38 | -44 | -8 | IFOF+ILF/ Sagittal Stratum[ILF+IFOF]/ OR+CC |
| | 65 | 0,02 | 18 | -80 | -1 | IFOF+ILF+Fmaj/ unc/ Visual Cortex+OR |
| | 61 | 0,022 | 34 | -59 | 0 | IFOF+ILF/ PTR[OR]/ CC+OR+Visual Cortex |
| | 49 | 0,037 | 31 | -52 | 13 | Fmaj+IFOF/ Tapetum/ CC+Visual Cortex |
| Left_G_woCVR | 86 | 0,01 | -24 | -75 | 0 | IFOF+ILF+Fmaj/ unc+*PTR[OR]*/ OR+Visual Cortex+CC |
| | 71 | 0,015 | 34 | -14 | -13 | IFOF/ Fornix+Stria Terminalis/ Hippocampus+OR+Fornix+*LGN+AcRad* |
| | 65 | 0,019 | -30 | -11 | -14 | ILF+*ATR*/ unc+*Fornix+Stria Terminalis*/ |

| | | | | | | |
|---|---|---|---|---|---|---|
| | | | | | | Amydala+Hippocampus+OR |
| | 50 | 0,039 | 18 | -80 | -1 | IFOF+ILF+Fmaj/ unc/ OR+Visual Cortex+CC |
| Right_G_woCVR | 117 | 0,004 | -38 | -44 | -8 | IFOF+ILF/ Sagittal Stratum[ILF+IFOF]+*PTR[OR]*/ OR+CC |
| | 103 | 0,005 | -31 | -64 | -1 | IFOF+ILF+*FMaj*/ PTR[OR]/ OR+CC+Visual Cortex |
| | 89 | 0,008 | -31 | -14 | -13 | -+*ATR*/ Fornix+Stria Terminalis/ OR+Hippocampus+Amydala |
| | 87 | 0,009 | 34 | -10 | -15 | -/ Fornix+Stria Terminalis/ **Hippocampus**+Amydala+OR+Fornix+*Ac Rad* |
| | 80 | 0,011 | 31 | -51 | 13 | Fmaj+IFOF/ Tapetum/ CC |
| | 78 | 0,011 | 25 | -37 | -8 | Cingulum (hippocampus)/ unc+*Cingulum (hippocampus)/* Hippocamous+OR+CC+*Cingulum* |
| | 72 | 0,014 | 40 | -36 | -11 | ILF+IFOF/ Sagittal Stratum[ILF+IFOF]/ OR+CC+*Fornix* |

### T-Test Finding

Table S14. Paired-t test results between the Right Global RNFLT correlations with the **Fractional Anisotropy** and the Left Global RNFLT correlations with the **Fractional Anisotropy** in Optic Radiata

| Mean difference (Right-Left) | T-statistic* | p-value | Degrees of Freedom | %95 CI-low | %95 CI-high | Cohen's_d |
|---|---|---|---|---|---|---|
| 0.0008869154 | 39.16 | 2.2e-16 | 199583 | 0.0008496655 | inf | 0.09 |

* One sided paired-t-test, Alternative H: The Right global RNFLT correlations are greater than the Left global RNFLT correlations, RNFLT correlations from the whole-brain TBSS when all CVR factors were controlled for the bilateral Optic Radiation ROI obtained from Juelich Histological Atlas multiplied with the mean FA skeleton.

### TBSS: Mean Diffusivity

Table S15. Whole Brain TBSS Results: Average Global Mean RNFLT negative correlations with the brain **Mean Diffusivity** (**without** CVRF controlling)

| | | | coordinates | | | |
|---|---|---|---|---|---|---|
| | Cluster size | pFWE-corr | x | y | z | Tract/Area |

| | | | | | | |
|---|---|---|---|---|---|---|
| avG_MD_woCVR | 44 | 0,021 | -30 | 18 | 18 | -/unc/Broca's Area/IFG(Harvard)/Insula(MNI) |

Table S16. Whole Brain TBSS Results: **Left** and **Right** Global Mean RNFLT negative correlations with the brain **Mean Diffusivity (with** CVRF controlling)

| | Voxels | pFWE-corr | MAX X (mm) | MAX Y (mm) | MAX Z (mm) | |
|---|---|---|---|---|---|---|
| Left_G_MD | 37 | 0,025 | -30 | 18 | 18 | -/unc/Broca's Area/IFG(Harvard)/Insula(MNI) |
| Right_G_MD | 39 | 0,023 | -30 | 18 | 18 | -/unc/Broca's Area/IFG(Harvard)/Insula(MNI) |

**CVRF: TBSS**

Table S17. Whole Brain TBSS Results Hypertension and Smoking negative correlations with the brain Fractional Anisotropy (when controlling for age and sex)

| CVR Factor | | | | coordinates | | |
|---|---|---|---|---|---|---|
| | Cluster size | pFWE-corr | x | y | z | Tract/Area |
| Hypertension_FA | 42 | 0,052 | -42 | -59 | 18 | Inferior Parietal Lobule |
| Smoking_FA | 67 | 0,018 | -28 | -47 | 32 | SLF/SLF/IPS+SLF+*OR+SSC(I)* |

Table S18. Whole Brain TBSS Results Hypertension and Diabetes positive correlations with the brain Mean Diffusivity (when controlling for age and sex)

| CVR Factor | Cluster Size | pFWE-corr | x | y | z | Tract/Area |
|---|---|---|---|---|---|---|
| Hypertension_ MD_pos | 157 | 0,001 | -15 | 26 | 17 | Forceps minor+*Cingulum(%1)*/CC/CC |
| | 109 | 0,003 | -13 | -24 | 28 | -/SCR/CC+CST |
| | 99 | 0,004 | -10 | 2 | 27 | -/CC/CC+Cingulum |
| | 87 | 0,005 | 30 | 32 | 6 | IFOF/-/-SLF(%2) |
| | 76 | 0,007 | -20 | 23 | 27 | SLF(%2)+UF(%1)/ACR/CC |
| | 65 | 0,011 | -18 | -2 | 35 | -/SCR/CC+*PremotorC*(%2) |
| | 64 | 0,011 | -34 | -8 | 20 | SLF/SLF/SSC(II)[Parietal Operculum]+CST(%2)+SSC(I)+*SSC(II)%1 +PMC%1* |
| | 54 | 0,015 | -20 | 44 | 7 | Forceps minor+ATR+*IFOF(%4)+Cingulum(%2)+ UF(%1)*/-/CC |

| | | | | | | |
|---|---|---|---|---|---|---|
| | 38 | 0,031 | -33 | -1 | 7 | SLF/ExtCapsule/- |
| | 36 | 0,035 | -17 | -29 | 29 | -/CC/CC |
| | 35 | 0,037 | -19 | -52 | 20 | Fmajor/CC/CC+*OR* |
| | 33 | 0,041 | 14 | 11 | 26 | -/CC/CC+*Cingulum(%1)* |
| | 32 | 0,044 | -35 | 23 | 15 | *IFOF+ILF(%1)+UF(%1)*/-/Broca44+*45* |
| | 31 | 0,048 | 19 | -13 | 43 | -/-/CST+PremotorC |
| Diabetes_MD_pos | 52 | 0,016 | 7 | -23 | 24 | -/CC/**CC**+Fornix+*Cingulum* |

Table S19. Whole Brain TBSS Results BMI positive and negative correlations with the brain Mean Diffusivity (when controlling for age and sex)

| | | | coordinates | | | |
|---|---|---|---|---|---|---|
| | Cluster size | pFWE _corr | x | y | z | Tract/Area |
| MD_pos | 40 | 0,032 | -14 | 33 | 6 | Fminor+*Cingulum*/CC/CC |
| MD_neg | 306 | 0 | 6 | -20 | -30 | CST+*ATR*(%2)/CST/- |
| | 73 | 0,005 | 41 | -60 | -10 | ILF(%2)/-/VisualC(%2)+OR |
| | 65 | 0,007 | -5 | -19 | -29 | CST/MiddleCerebellar Pedincle/- |
| | 56 | 0,01 | 28 | -62 | 24 | IFOF+[*SLF%1+ILF%1+FMajor%2*]/-/OR+CC+*SPL*(%1) |
| | 53 | 0,011 | 33 | -57 | 29 | ILF(%2)+*SLF*(%2)/-/OR+IPS+*OR*(%3) |
| | 52 | 0,011 | 38 | -19 | -9 | IFOF+ILF/Sagittal Stratum[ILF+IFOF]/OR+Insula+AR+Hipp(%2)+*IFOF(%1)+CC(%1)* |
| | 43 | 0,017 | -30 | -85 | -6 | IFOF(%2)+*ILF*(%1)/-/VisualCortex+OR |
| | 42 | 0,019 | 32 | -66 | -9 | ILF+IFOF+*Cingulum(Hipp%1)*/-/VisualCortex+OR+*CC*(%4) |
| | 38 | 0,025 | 36 | -31 | -21 | -*ILF*(%4)+*Cingulum(Hipp%1)*/-/Hippocampus+**OR** |
| | 31 | 0,036 | 6 | -17 | -7 | ATR/-/- |
| | 29 | 0,041 | 33 | 25 | -14 | ---*IFOF*(%1) |
| | 29 | 0,041 | -30 | -59 | -12 | ILF+IFOF/-/OR |
| | 29 | 0,041 | 33 | -57 | 18 | ILF+IFOF+FMajor(%4)+SLF(%1)/PTR[OR]/OR+CC |
| | 27 | 0,049 | 29 | -16 | -26 | Cingulum(Hipp)/Hippocampus/**Hippocampus** |

Table S20. Partial correlation coefficients [for pre-chosen **ROIs** below, from the literature, the number of ROIs was 49 in total] between the average Global RNFLT and brain regional gray/white matter volume measures (age, sex and retina scan radius were always controlled for)

| **RNFLT correlations** | |
|---|---|
| **without CVRF correction** | **with CVRF correction** |

| | raw | FDR corrected | | raw | FDR corrected |
|---|---|---|---|---|---|
| Whole-brain GM | r=0.044, p=0.226 | | Whole-brain GM | r=0.049, p=0.179 | |
| Whole-brain WM | r=0.100, p=0.006** | p=0.058 | Whole-brain WM | r=0.101, p=0.005** | p=0.048* |
| Right Caudate | r=0.035, p=0.337 | | Right Caudate | r=0.035, p=0.297 | |
| Left Caudate | r=0.040, p=0.265 | | Left Caudate | r=0.044, p=0.228 | |
| Right Cerebellum | r=0.081, p=0.025* | p=0.113 | Right Cerebellum | r=0.086, p=0.018* | p=0.096 |
| Left Cerebellum | r=0.102, p=0.005** | p=0.058 | Left Cerebellum | r=0.107, p=0.003** | p=0.036* |
| Right Cerebellar WM | r=-0.033, p=0.358 | | Right Cerebellar WM | r=-0.032, p=0.384 | |
| Left Cerebellar WM | r=-0.021, p=0.569 | | Left Cerebellar WM | r=-0.018, p=0.625 | |
| Right Hippocampus | r=0.069, p=0.057 | | Right Hippocampus | r=0.067, p=0.064 | |
| Left Hippocampus | r=0.046, p=0.207 | | Left Hippocampus | r=0.044, p=0.220 | |
| Right Pallidum | r=0.065, p=0.074 | | Right Pallidum | r=0.062, p=0.086 | |
| Left Pallidum | r=0.068, p=0.059 | | Left Pallidum | r=0.066, p=0.070 | |
| Right Putamen | r=0.045, p=0.214 | | Right Putamen | r=0.045, p=0.217 | |
| Left Putamen | r=0.057, p=0.113 | | Left Putamen | r=0.058, p=0.110 | |
| Right Thalamus | r=0.010, p=0.779 | | Right Thalamus | r=0.015, p=0.671 | |
| Left Thalamus | r=-0.008, p=0.816 | | Left Thalamus | r=-0.004, p=0.908 | |
| Optic Chiasm | r=-0.050, p=0.165 | | Optic Chiasm | r=-0.055, p=0.128 | |
| Right Ant. Cingulate | r=0.035, p=0.330 | | Right Ant. Cingulate | r=0.039, p=0.285 | |
| Left Ant. Cingulate | r=-0.021, p=0.558 | | Left Ant. Cingulate | r=-0.018, p=0.614 | |

| | | | | | |
|---|---|---|---|---|---|
| Right Calcarine | r=0.186, p=0.000*** | p=0.000*** | Right Calcarine | r=0.190, p=0.000*** | p=0.000*** |
| Left Calcarine | r=0.165, p=0.000*** | p=0.000*** | Left Calcarine | r=0.171, p=0.000*** | p=0.000*** |
| Right Cuneus | r=0.041, p=0.263 | | Right Cuneus | r=0.040, p=0.273 | |
| Left Cuneus | r=0.041, p=0.253 | | Left Cuneus | r=0.042, p=0.248 | |
| Right Entorhinal | r=0.026, p=0.468 | | Right Entorhinal | r=0.024, p=0.500 | |
| Left Entorhinal | r=0.057, p=0.116 | | Left Entorhinal | r=0.056, p=0.121 | |
| Right Fusiform | r=0.031, p=0.390 | | Right Fusiform | r=0.033, p=0.368 | |
| Left Fusiform | r=0.067, p=0.064 | p=0.219 | Left Fusiform | r=0.072, p=0.047* | p=0.188 |
| Right Lingual | r=0.081, p=0.026* | p=0.113 | Right Lingual | r=0.083, p=0.022* | p=0.096 |
| Left Lingual | r=0.041, p=0.255 | | Left Lingual | r=0.043, p=0.236 | |
| Right Inf. Occipital | r=-0.029, p=0.429 | | Right Inf. Occipital | r=-0.028, p=0.445 | |
| Left Inf. Occipital | r=0.022, p=0.535 | | Left Inf. Occipital | r=0.023, p=0.519 | |
| Right Mid. Cingulate | r=-0.021, p=0.561 | | Right Mid. Cingulate | r=-0.020, p=0.588 | |
| Left Mid. Cingulate | r=-0.086, p=0.017* | p=0.101 | Left Mid. Cingulate | r=-0.083, p=0.022* | p=0.096 |
| Right Mid. Occipital | r=0.024, p=0.499 | | Right Mid. Occipital | r=0.024, p=0.514 | |
| Left Mid. Occipital | r=-0.006, p=0.862 | | Left Mid. Occipital | r=-0.006, p=0.876 | |
| Right Mid. Temporal | r=0.033, p=0.363 | | Right Mid. Temporal | r=0.036, p=0.325 | |
| Left Mid. Temporal | r=0.011, p=0.763 | | Left Mid. Temporal | r=0.013, p=0.717 | |
| Right Occipital Pole | r=0.039, p=0.279 | | Right Occipital Pole | r=0.039, p=0.283 | |
| Left Occipital Pole | r=0.085, p=0.019* | p=0.101 | Left Occipital Pole | r=0.085, p=0.019* | p=0.096 |
| Right Occ. Fusiform | r=0.043, p=0.237 | | Right Occ. Fusiform | r=0.044, p=0.224 | |
| Left Occ. Fusiform | r=0.028, p=0.436 | | Left Occ. Fusiform | r=0.031, p=0.393 | |
| Right Sup. Occipital | r=0.060, p=0.097 | | Right Sup. Occipital | r=0.060, p=0.098 | |

| | | | | | |
|---|---|---|---|---|---|
| Left Sup. Occipital | r=0.034, p=0.354 | | Left Sup. Occipital | r=0.034, p=0.342 | |
| Right LGN | r=0.027, p=0.464 | | Right LGN | r=0.029, p=0.423 | |
| Left LGN | r=-0.002, p=0.945 | | Left LGN | r=0.000, p=0.997 | |
| Right Post. Cingulate | r=-0.005, p=0.892 | | Right Post. Cingulate | r=-0.007, p=0.870 | |
| Left Post. Cingulate | r=-0.050, p=0.168 | | Left Post. Cingulate | r=-0.052, p=0.154 | |
| Right Parahippocampal | r=0.086, p=0.017* | p=0.101 | Right Parahippocampal | r=0.087, p=0.017* | p=0.096 |
| Left Parahippocampal | r=0.084, p=0.019* | p=0.101 | Left Parahippocampal | r=0.087, p=0.016* | p=0.096 |

***
