## supplemental figures and tables for "Retinal nerve fibre layer thickness reflects characteristics of brain grey and white matter": 1_Supplement_Figures_final_Nazife_Ayyildiz_8MAY2025.pdf

### SUPPLEMENTARY FIGURES

#### VBM

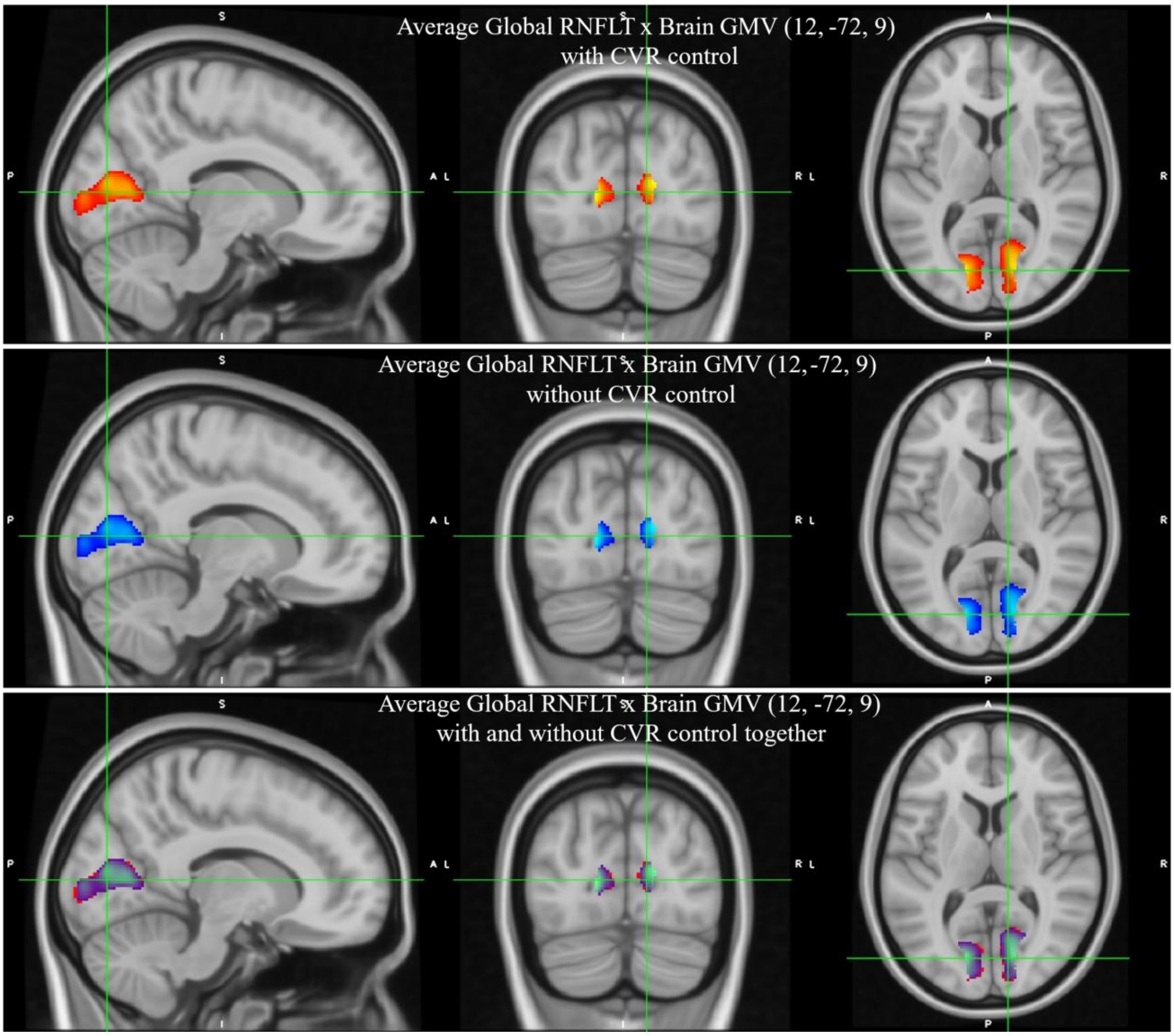


Figure S1. Average Global Mean RNFLT positive correlations with the brain GMV with and without CVR factors controlling (n=769). **Upper** figure shows the correlations when controlling for the CVR factors i.e., BMI, LDL and HDL Cholesterol scores and Diabetes, Hypertension, Smoking and Physical Activity status (red) in addition to age, sex, total intracranial volume and retina scan radius. **Middle** figure shows the correlations without CVR factor controlling, only age, sex, total intracranial volume and retina scan radius were controlled (blue). Lower figure shows both correlations together (overlap, purple). Results shown on MNI152_T1_05.mm template, corrected at cluster-level pFWE<0.05 and uncorrected at voxel-level p<0.001. RNFLT: Retinal Nerve Fiber Layer Thickness, GMV: Gray Matter Volume.

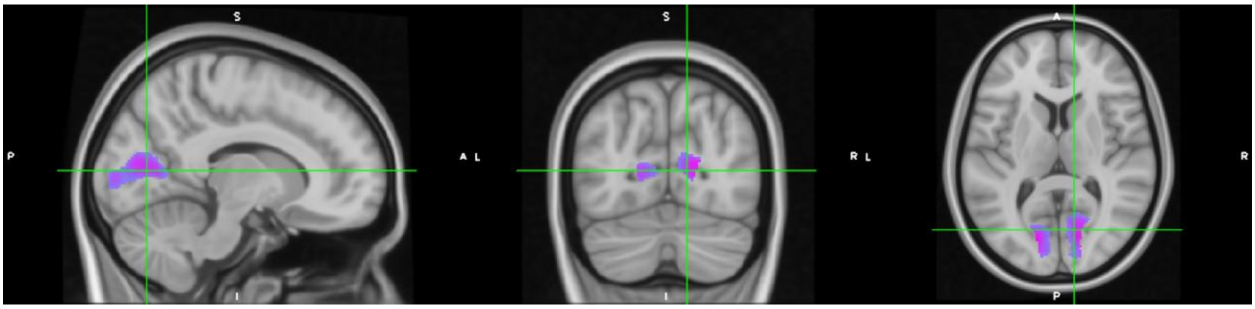


Figure S2. Conjunction (overlap) between Average Global Mean RNFLT positive correlations with and without CVR when age, sex, TIV and average retina scan radius were controlled

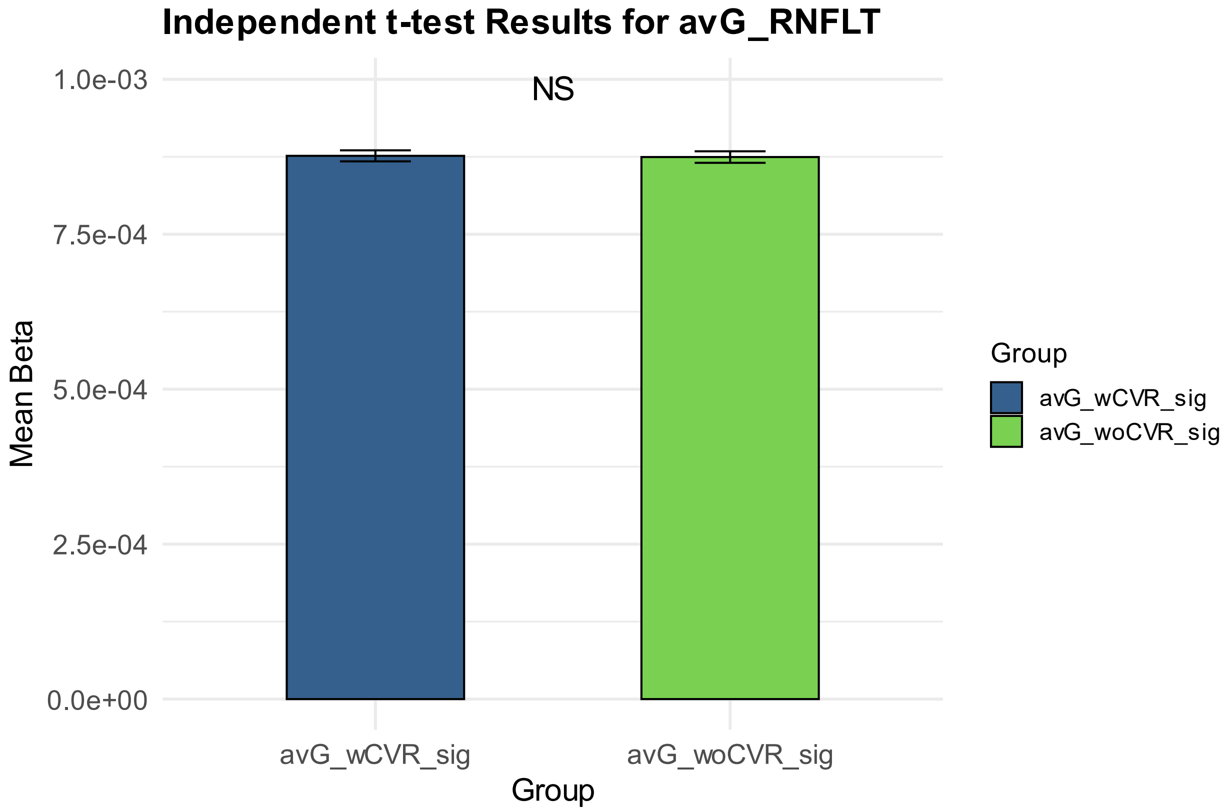


Figure S3. Comparison of the Beta values of the regions between Average Global Mean RNFLT positive correlations **with** and **without** controlling for the CVR factors in addition to age, sex, TIV and average retina scan radius variables. The independent t-test results were not statistically significant between the beta values [T (5796) = 0.31, p=0.76, cohen's_d = (0.0081)].

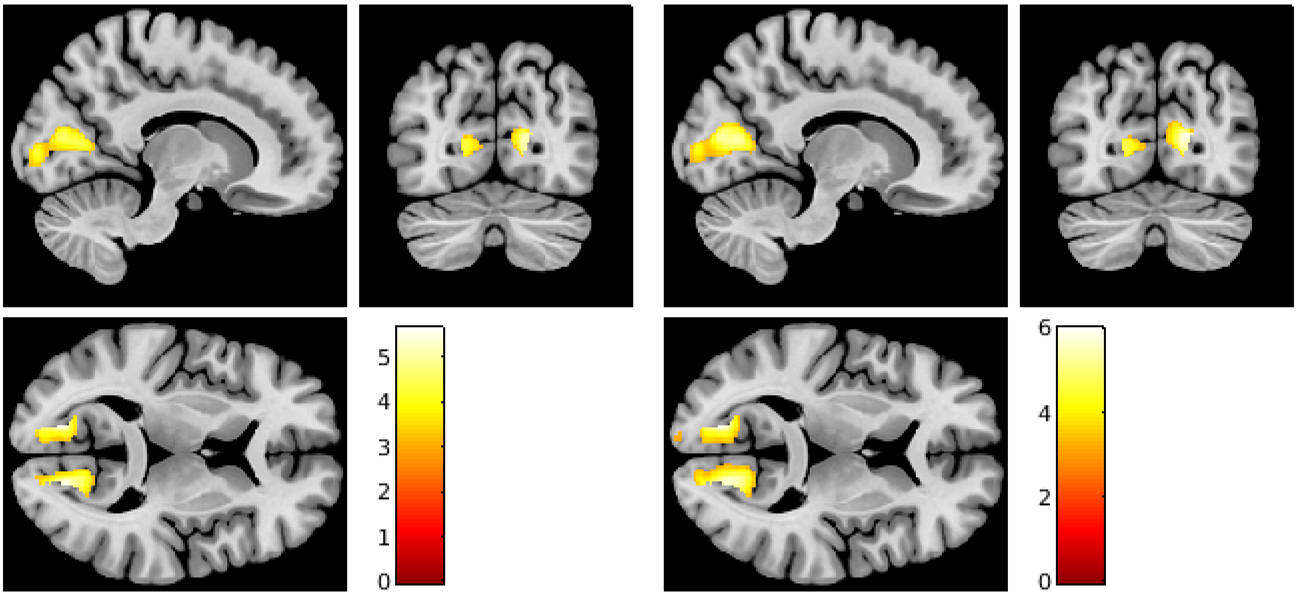


Figure S4. **Left** (left side) **and Right** (right side) **Global Mean RNFLT** positive correlations with the brain GMV when controlling for age, sex, total intracranial volume, retina scan radius and CVR factors i.e., BMI, LDL and HDL Cholesterol scores and Diabetes, Hypertension, Smoking and Physical Activity status **(n=769).** Results shown on MNI registered CAT-T1_IXI 555 GS standard atlas, Neurological View. Coordinates (12, -72, 9). Color bar shows T statistics corrected at cluster-level pFWE<0.05 and uncorrected at voxel-level p<0.001. RNFLT: Retinal Nerve Fiber Layer Thickness, GMV: Gray Matter Volume.

**Comparison** of the Left Global RNFLT Correlations to the Right Global RNFLT Correlations when CVR factors were and were not taken into account in addition to age, sex, TIV and Retina Scan Radius variables:

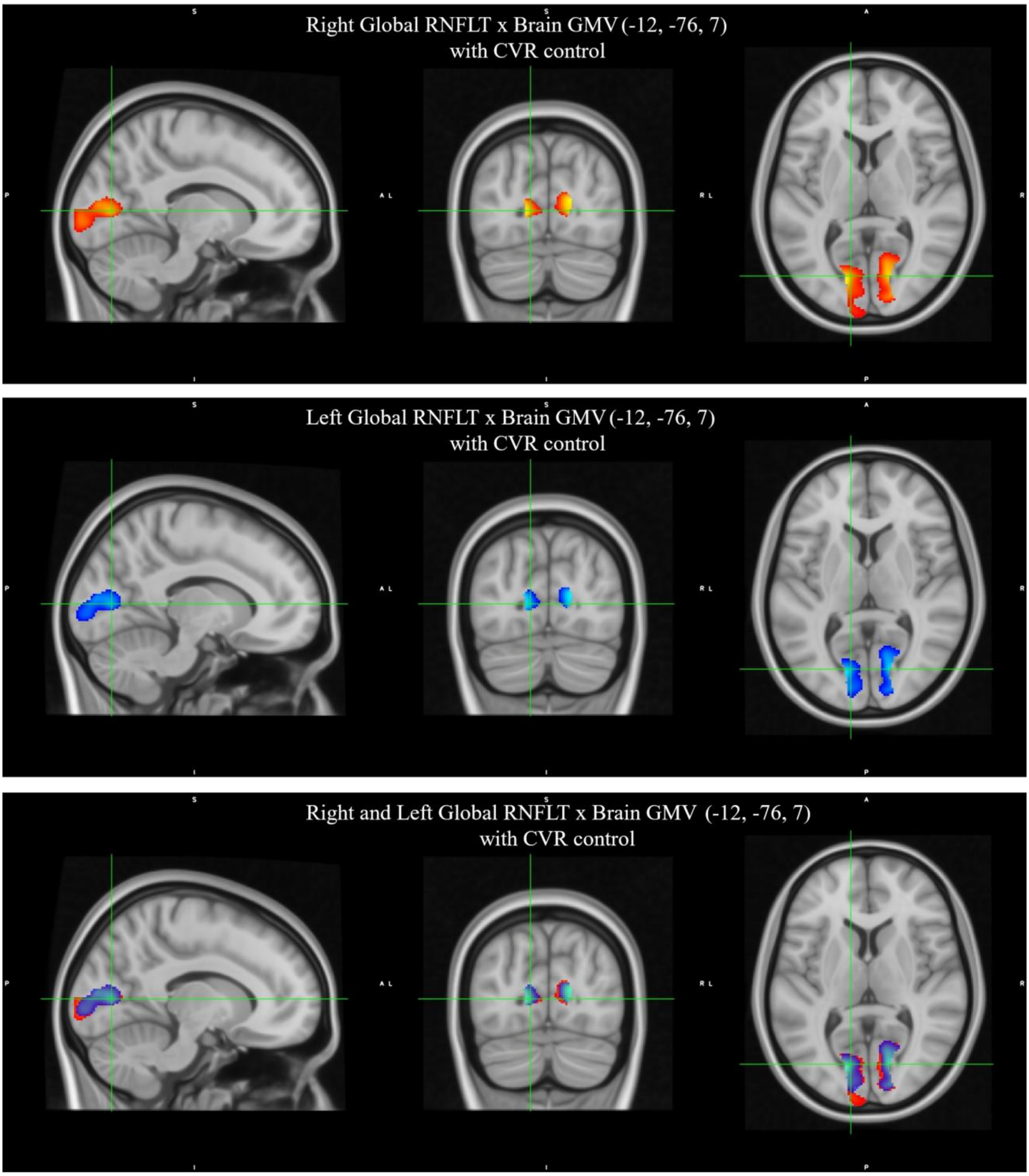


Figure S5. Comparison of the **Right** and **Left Global Mean RNFLT** positive correlations with the brain GMV when controlling for age, sex, total intracranial volume, retina scan radius and **CVR** factors i.e., BMI, LDL and HDL Cholesterol scores and Diabetes, Hypertension, Smoking and Physical Activity status **(n=769).** Results shown, at cluster-level corrected p< 0.05 for FWER with an uncorrected p<0.001 voxel-level clustering threshold, on MNI152_T1_0.5mm standard atlas, Neurological View. RNFLT: Retinal Nerve Fiber Layer Thickness

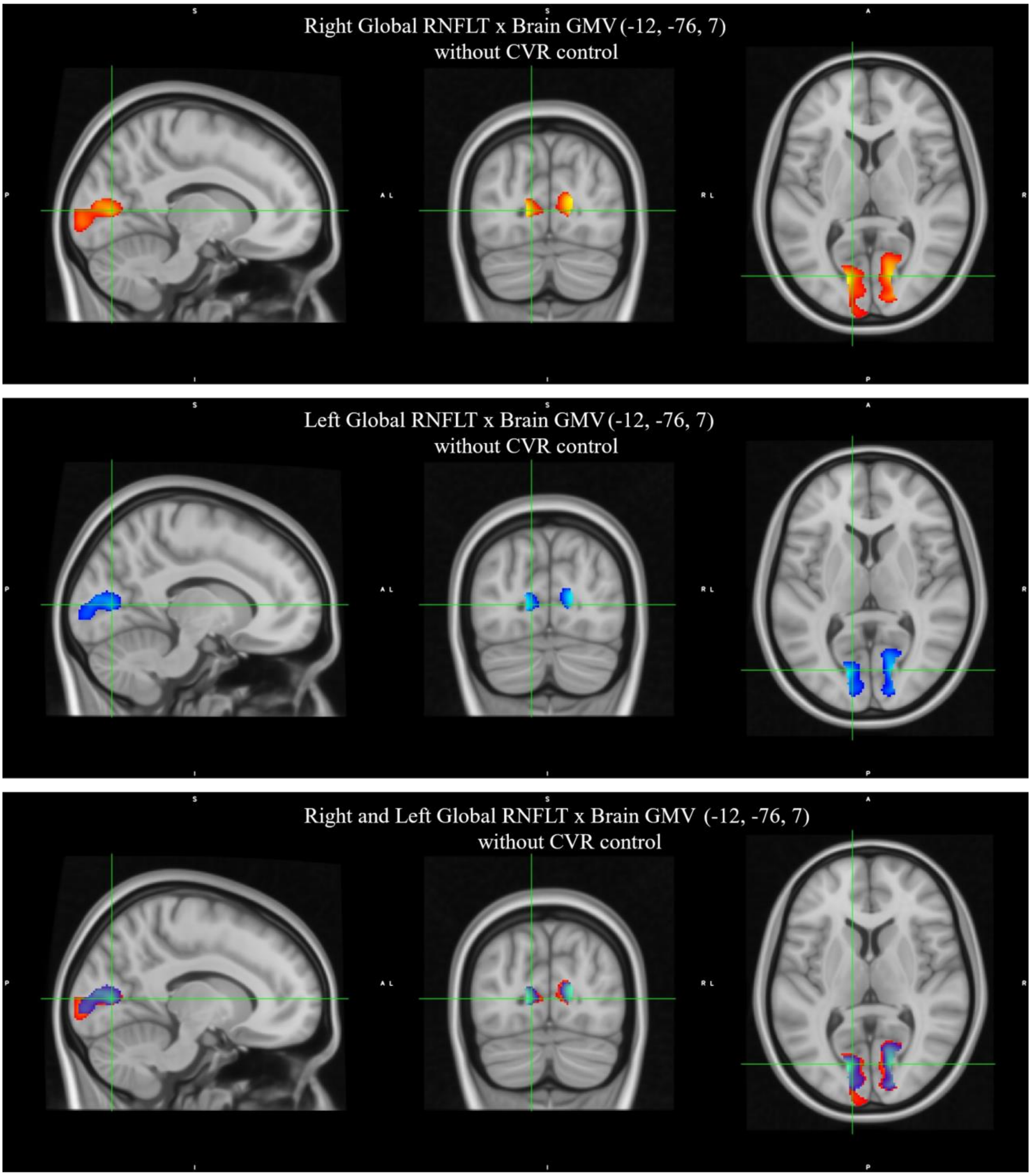


Figure S6. Comparison of the **Right** and **Left Global Mean RNFLT** positive correlations with the brain GMV when controlling for only age, sex, total intracranial volume and retina scan radius **(n=769).** Results shown, at cluster-level corrected $p < 0.05$ for FWER with an uncorrected $p < 0.001$ voxel-level clustering threshold, on MNI152_T1_0.5mm standard atlas, Neurological View. RNFLT: Retinal Nerve Fiber Layer Thickness

**Comparison** of the Right Global RNFLT Correlations when CVR factors were and were not taken into account in addition to age, sex, TIV and Retina Scan Radius variables:

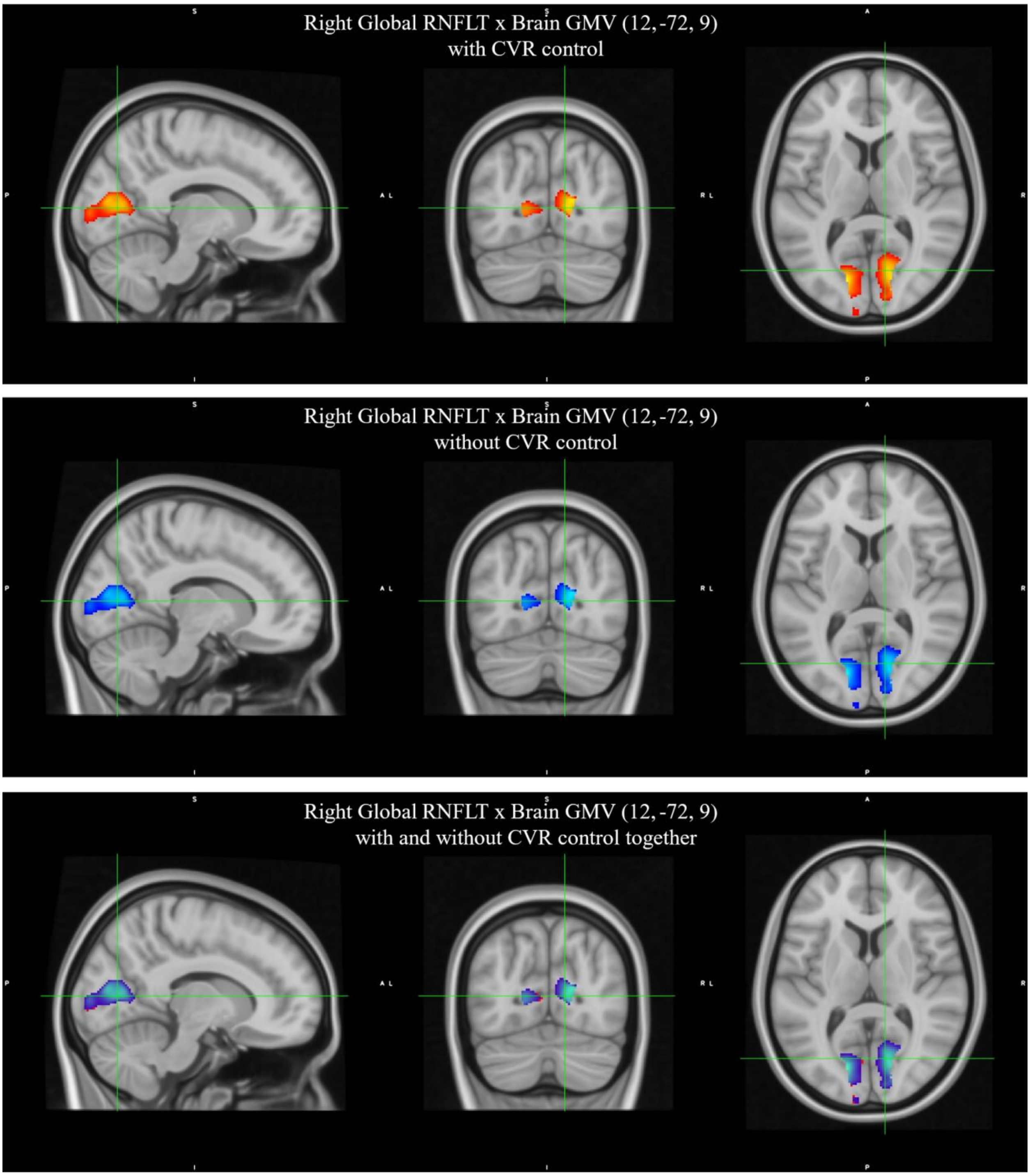


Figure S7. **Right** Global Mean RNFLT positive correlations with the brain GMV **with and without CVR factors** controlling (n=769). **Upper** figure shows the correlations when controlling for the CVR factors i.e., BMI, LDL and HDL Cholesterol scores and Diabetes, Hypertension, Smoking and Physical Activity status (red) in addition to age, sex, total intracranial volume and retina scan radius. **Middle** figure shows the correlations when controlling for only age, sex, total intracranial volume and related retina scan radius (blue)**.** Lower figure shows both correlations together (overlap, purple). Results shown on MNI152_T1_0.5mm template, corrected at cluster-level pFWE<0.05 and uncorrected at voxel-level p<0.001. RNFLT: Retinal Nerve Fiber Layer Thickness, GMV: Gray Matter Volume

**Comparison** of the Right Global RNFLT Correlations when CVR factors were and were not taken into account in addition to age, sex, TIV and Retina Scan Radius variables:

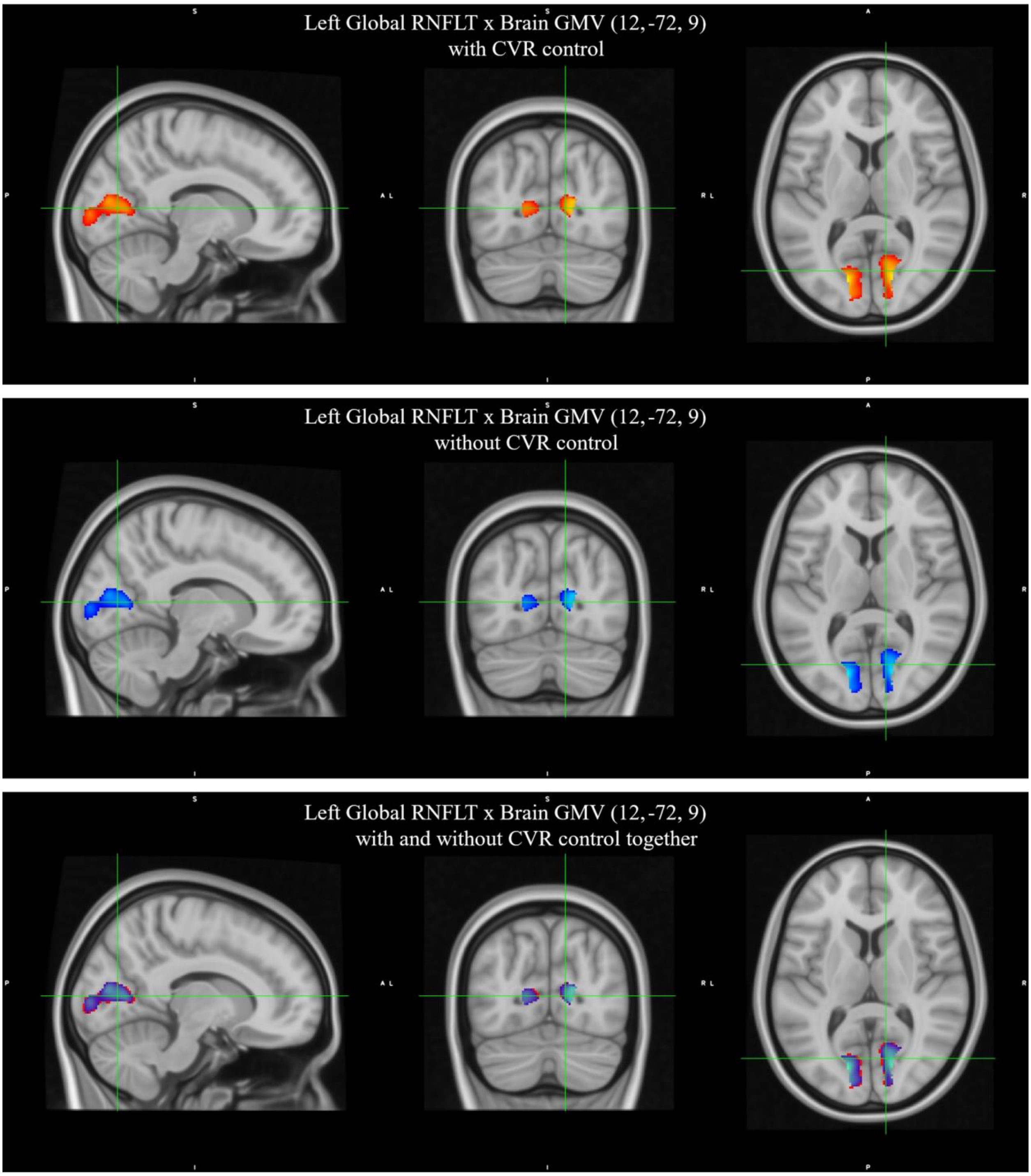


Figure S8. **Left** Global Mean RNFLT positive correlations with the brain GMV **with and without CVR factors** controlling (n=769). **Upper** figure shows the correlations when controlling for the CVR factors i.e., BMI, LDL and HDL Cholesterol scores and Diabetes, Hypertension, Smoking and Physical Activity status (red) in addition to age, sex, total intracranial volume and retina scan radius. **Middle** figure shows the correlations when controlling for only age, sex, total intracranial volume and related retina scan radius (blue)**.** Lower figure shows both correlations together (overlap, purple). Results shown on MNI152_T1_0.5mm template, corrected at cluster-level pFWE<0.05 and uncorrected at voxel-level p<0.001. RNFLT: Retinal Nerve Fiber Layer Thickness, GMV: Gray Matter Volume

#### TBSS: FRACTIONAL ANISOTROPY

Right Global RNFLT x Brain FA (39, 42, 5) with CVR control

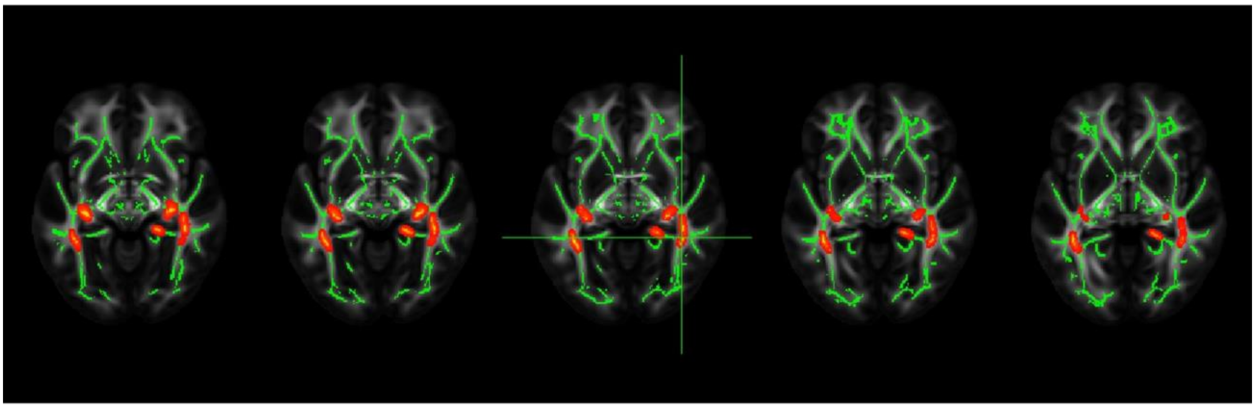

Right Global RNFLT x Brain FA (39, 42, 5) without CVR control

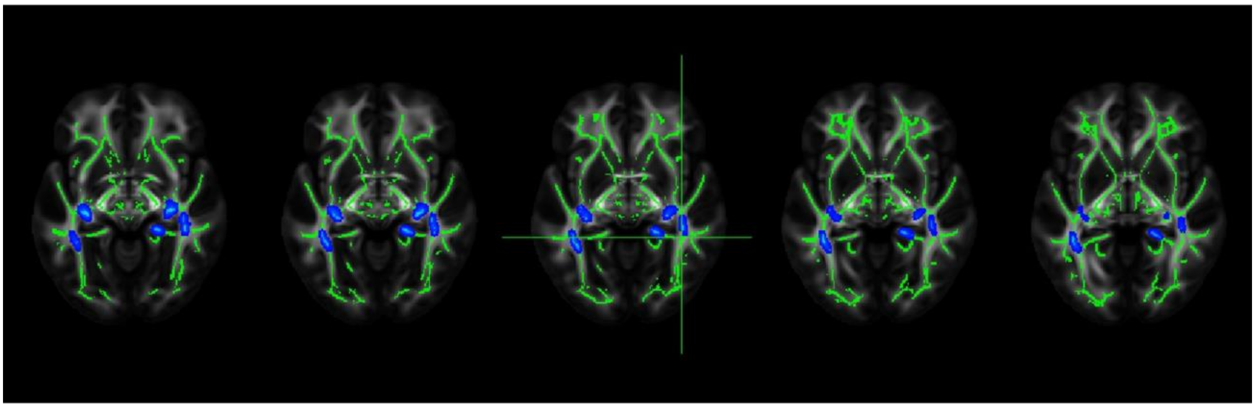

Right Global RNFLT x Brain FA (39, 42, 5) with and without CVR control (together)

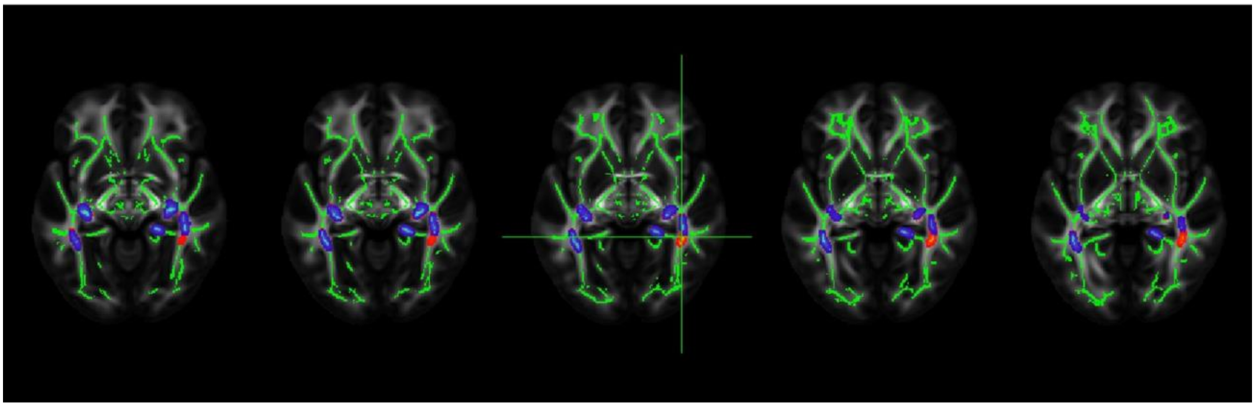

Figure S9. **Right** Global Mean RNFLT positive correlations with brain Fractional Anisotropy when controlling for the CVR factors i.e., BMI, LDL and HDL Cholesterol scores and Diabetes, Hypertension, Smoking and Physical Activity status (**i.e., with CVR**) in addition to age, sex and retina scan radius (**i.e., without CVR**), n=550. Results shown, at cluster-level corrected p< 0.05 for FWER with an uncorrected p<0.001 voxel-level clustering threshold, on MNI-registered FSL_HCP1065_FA_1mm standard atlas, Neurological View. RNFLT: Retinal Nerve Fiber Layer Thickness

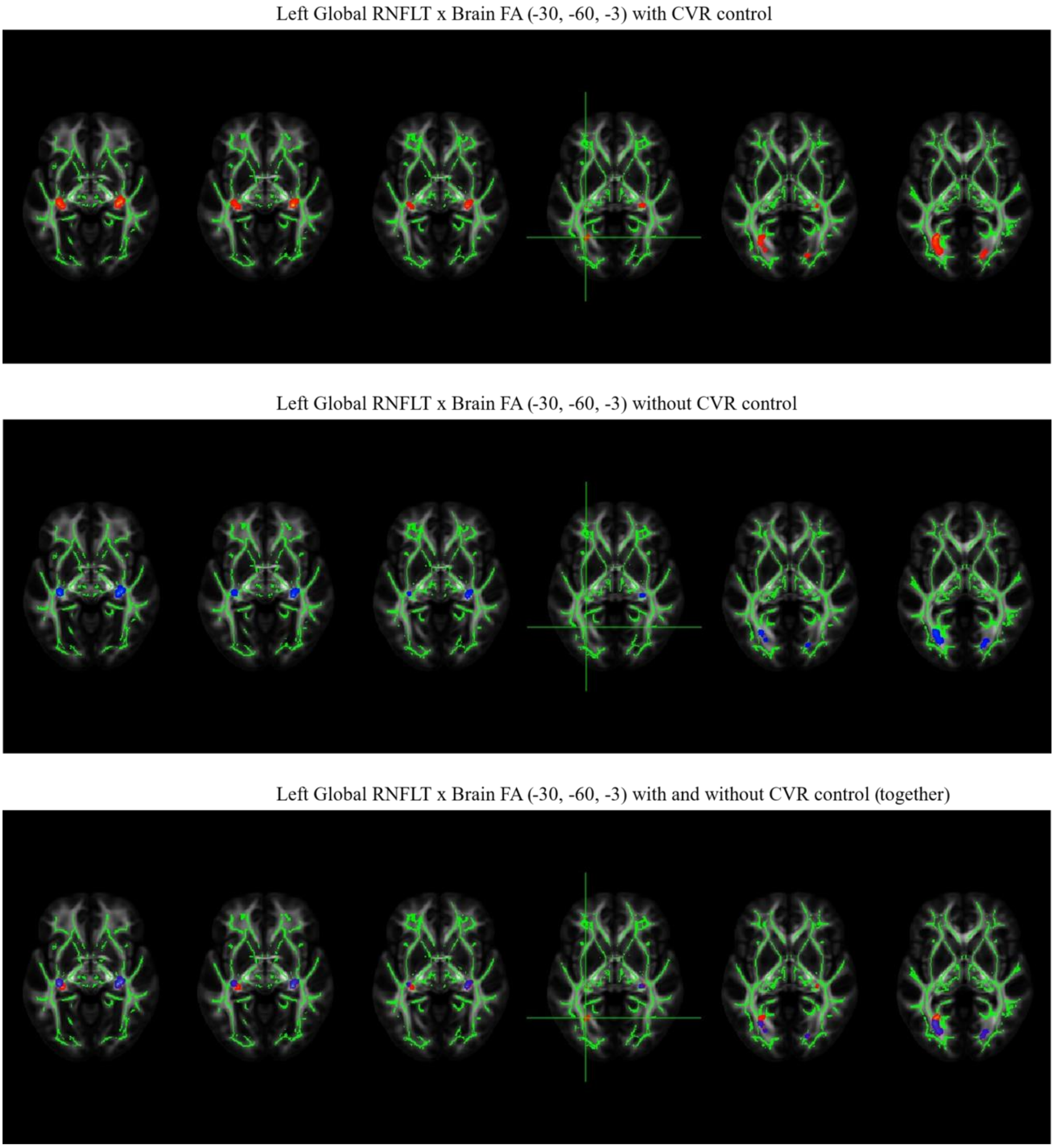


Figure S10. **Left** Global Mean RNFLT positive correlations with brain Fractional Anisotropy when controlling for the CVR factors i.e., BMI, LDL and HDL Cholesterol scores and Diabetes, Hypertension, Smoking and Physical Activity status (**i.e., with CVR**) in addition to age, sex and retina scan radius (**i.e., without CVR**), n=550. Results shown, at cluster-level corrected $p < 0.05$ for FWER with an uncorrected $p < 0.001$ voxel-level clustering threshold, on MNI-registered FSL_HCP1065_FA_1mm standard atlas, Neurological View. RNFLT: Retinal Nerve Fiber Layer Thickness

Left Global RNFLT x Brain FA (-29, -58, -1)

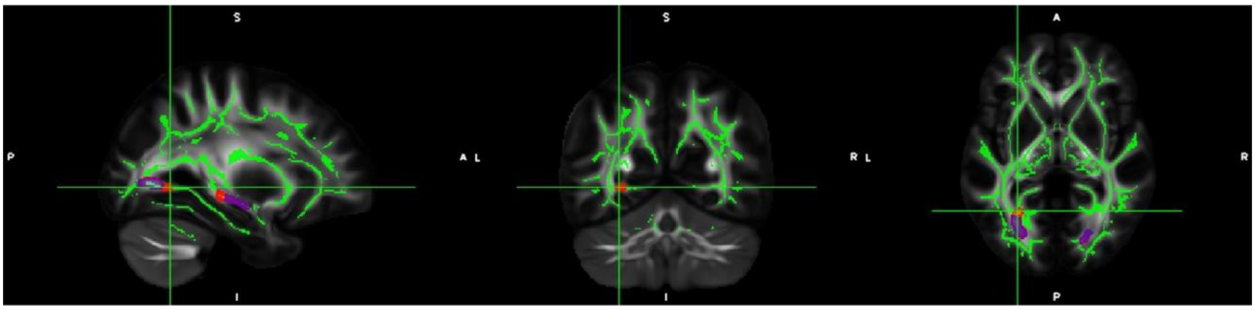


Right Global RNFLT x Brain FA (40, -42, -5)

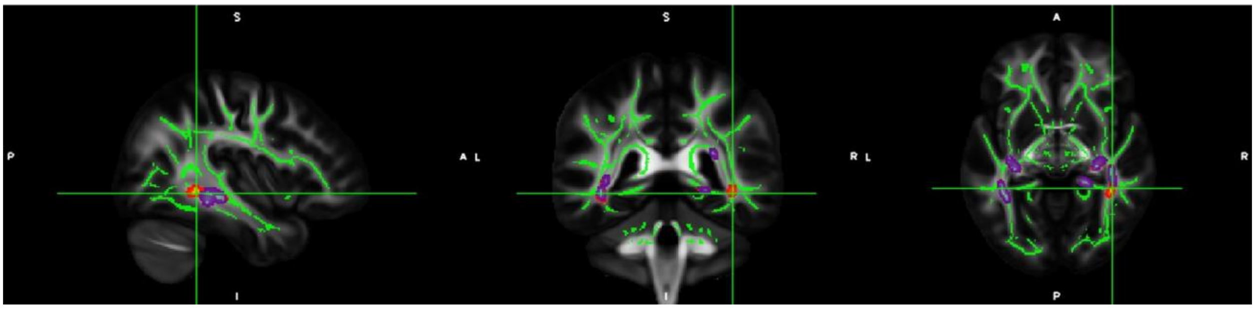


Figure S11. Left Global (upper side) and Right Global (lower side) Mean RNFLT positive correlations with brain Fractional Anisotropy. **Blue** shows the results when controlling for only age, sex, retina scan radius (**i.e., without CVR**) and **red** shows the results when controlling CVR factors additionally (**i.e., with CVR**), and **purple** shows the overlap between blue and red colors (n=550). Results shown, at cluster-level corrected p< 0.05 for FWER with an uncorrected p<0.001 voxel-level clustering threshold, on MNI-registered FSL_HCP1065_FA_1mm standard atlas, Neurological View. RNFLT: Retinal Nerve Fiber Layer Thickness

**Comparison** of the Left and Right Global RNFLT positive correlations with the Brain Fractional Anisotropy, n=550, with and without CVR factors:

Right Global RNFLT x Brain FA (20, -40, -4) with CVR control

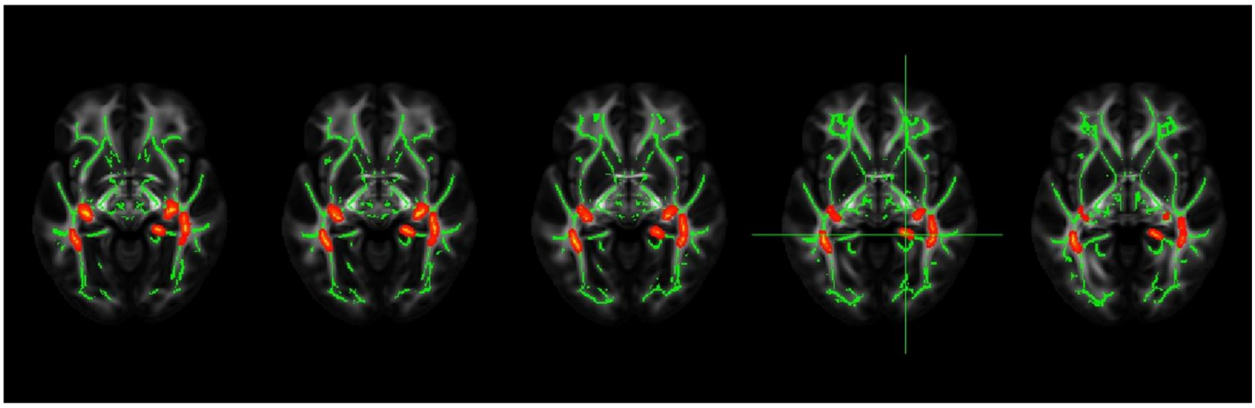

Left Global RNFLT x Brain FA (20, -40, -4) with CVR control

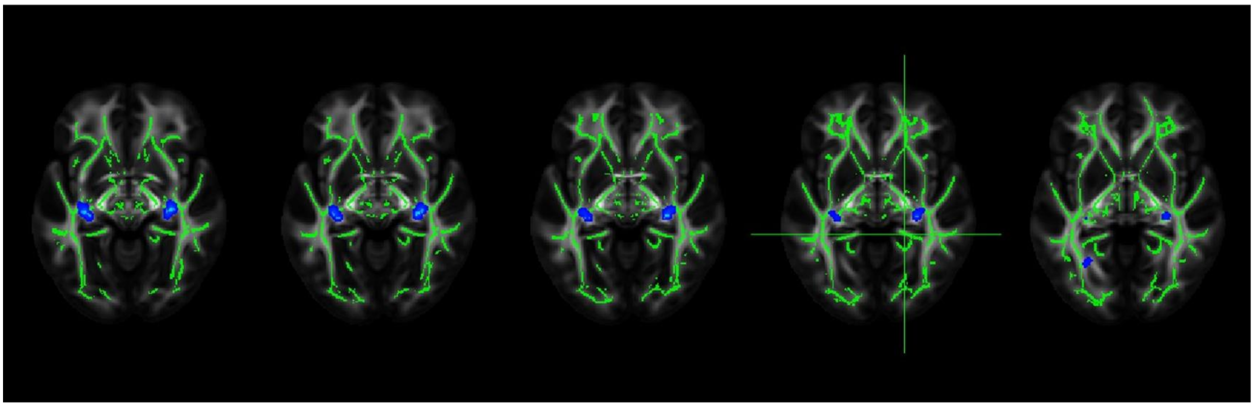

Right and Left Global RNFLT x Brain FA (20, -40, -4) with CVR control (together)

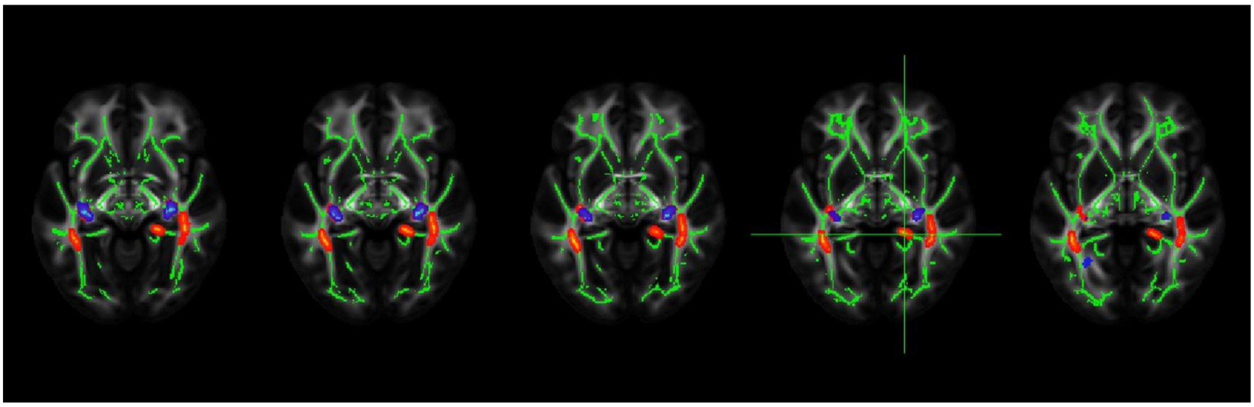

Right and Left Global RNFLT x Brain FA (24, -48, 20) with CVR control (together)

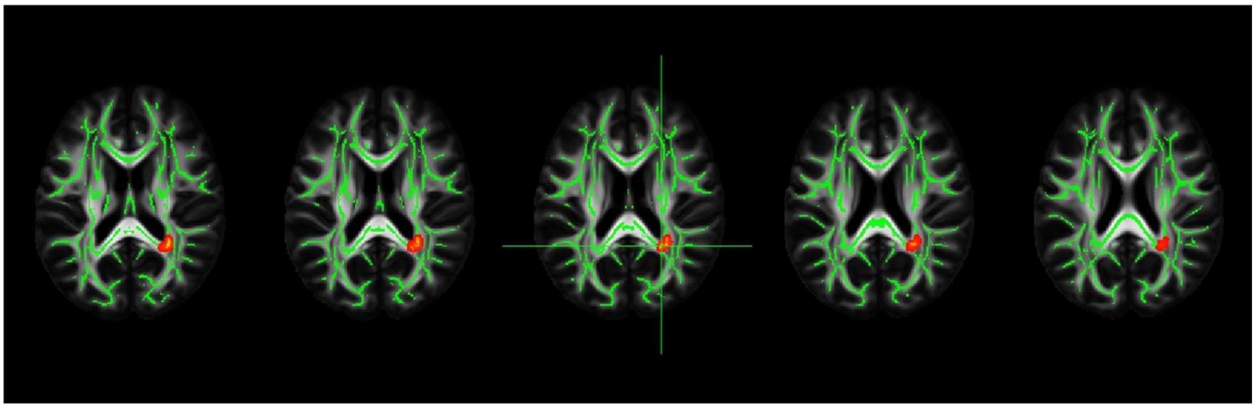

Figure S12. Comparison of the **Right** and **Left Global Mean RNFLT** positive correlations with the brain FA when controlling for the CVR factors i.e., BMI, LDL and HDL Cholesterol scores and Diabetes, Hypertension, Smoking

and Physical Activity status (**i.e., with CVR**) in addition to age, sex and regarding retina scan radius **(n=550).** Results shown, at cluster-level corrected p< 0.05 for FWER with an uncorrected p<0.001 voxel-level clustering threshold, on MNI-registered FSL_HCP1065_FA_1mm standard atlas, Neurological View. RNFLT: Retinal Nerve Fiber Layer Thickness, FA: Fractional Anisotropy

Right Global RNFLT x Brain FA (20, -40, -4) without CVR control

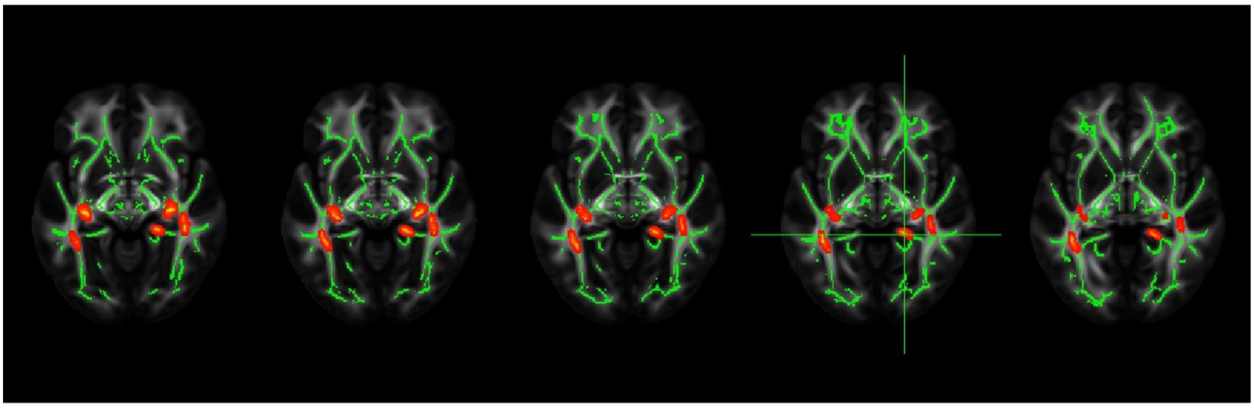

Left Global RNFLT x Brain FA (20, -40, -4) without CVR control

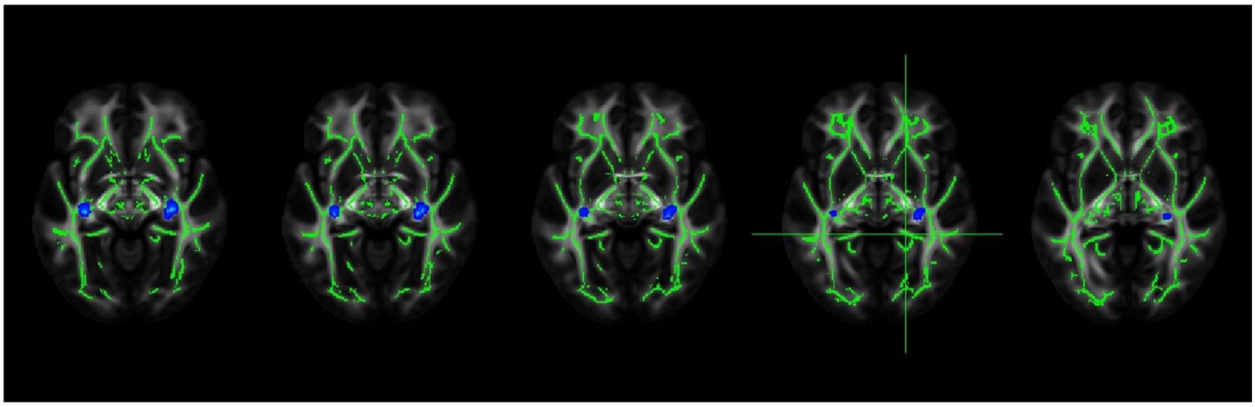

Right and Left Global RNFLT x Brain FA (20, -40, -4) without CVR control (together)

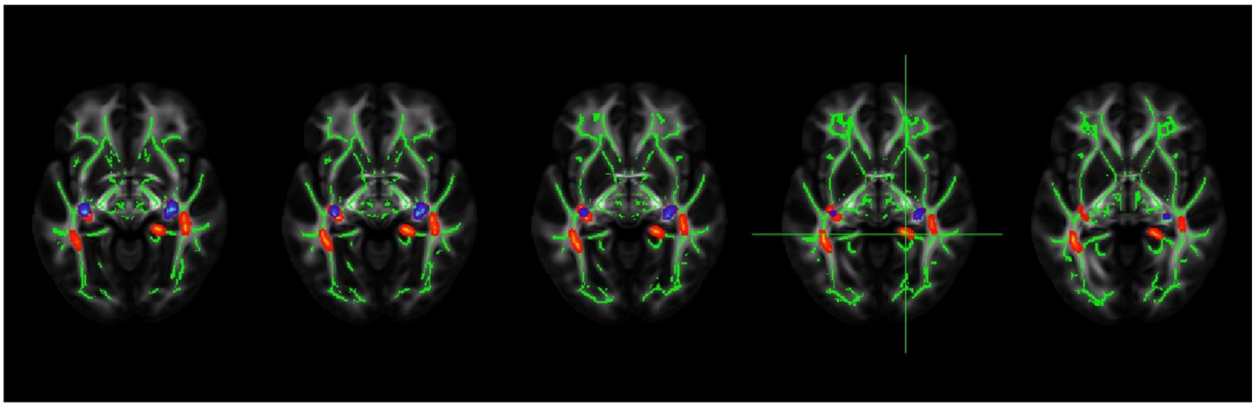

Right and Left Global RNFLT x Brain FA (24, -48, 20) without CVR control (together)

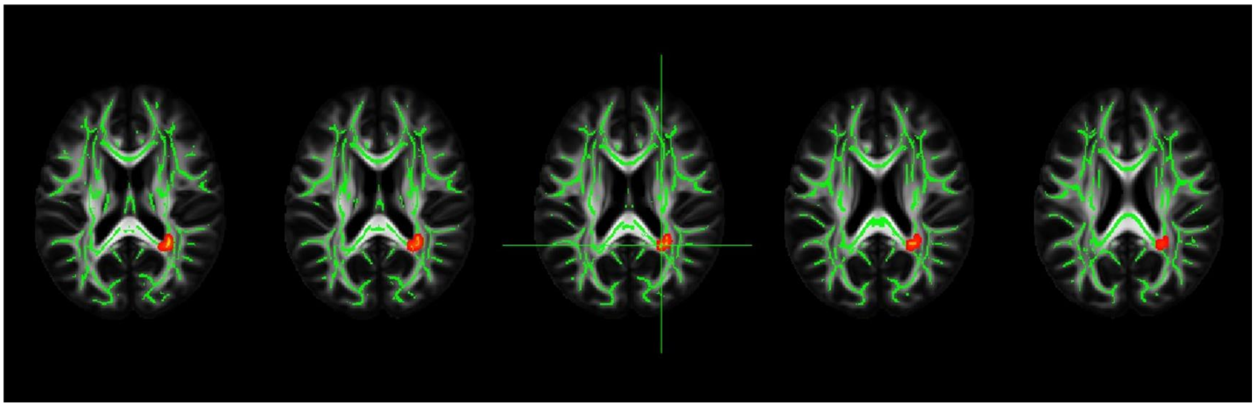

Figure S13. Comparison of the **Right** and **Left Global Mean RNFLT** positive correlations with the brain FA when controlling for only age, sex and regarding retina scan radius [(**i.e., without CVR, (n=550)].** Results shown, at cluster-level corrected p< 0.05 for FWER with an uncorrected p<0.001 voxel-level clustering threshold, on MNI-registered FSL_HCP1065_FA_1mm standard atlas, Neurological View. RNFLT: Retinal Nerve Fiber Layer Thickness, FA: Fractional Anisotropy

#### TBSS: MEAN DIFFUSIVITY

Left Global RNFLT x Brain MD (-28, 16, 20) with CVR control

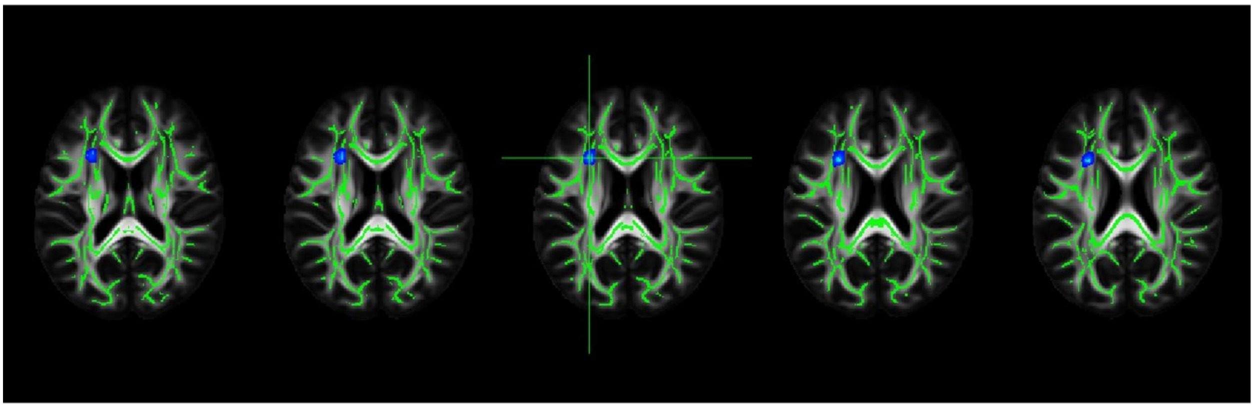

Left Global RNFLT x Brain MD (-28, 16, 20) without CVR control

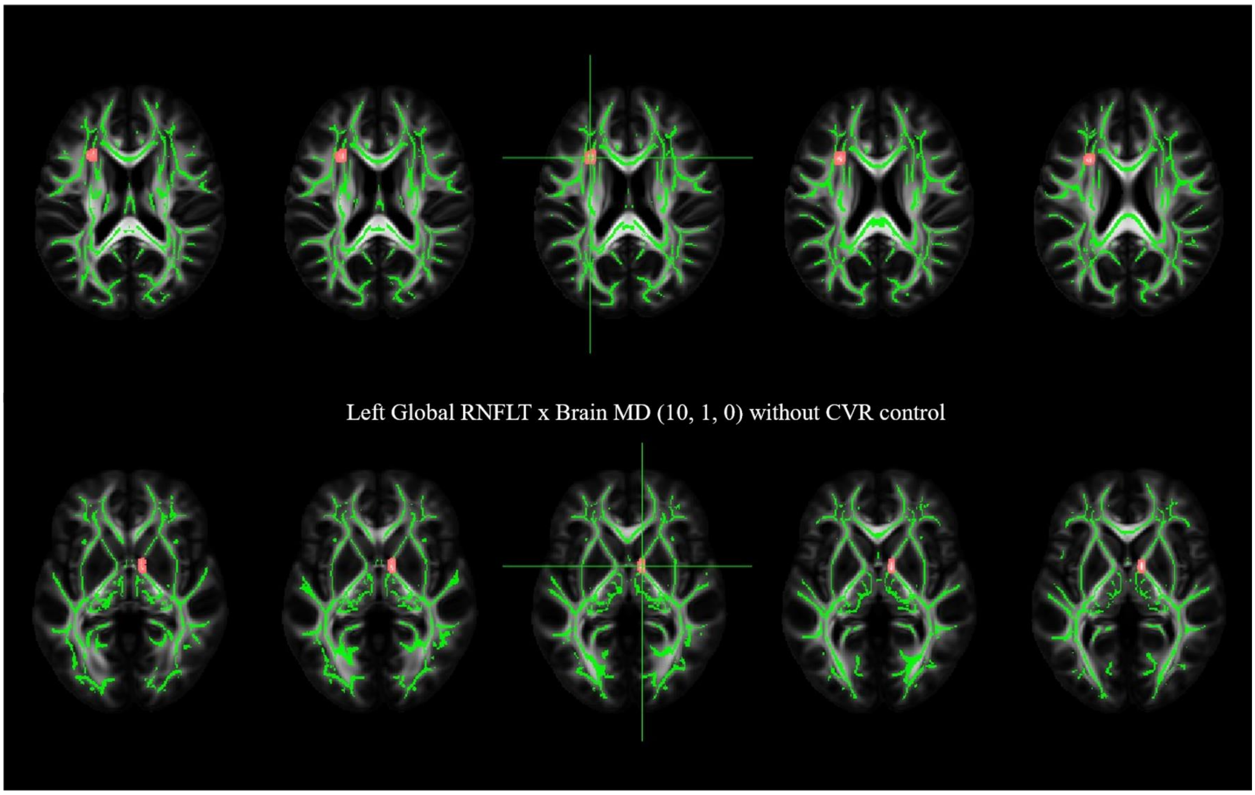


Left Global RNFLT x Brain MD (-28, 16, 20) with and without CVR control (together)

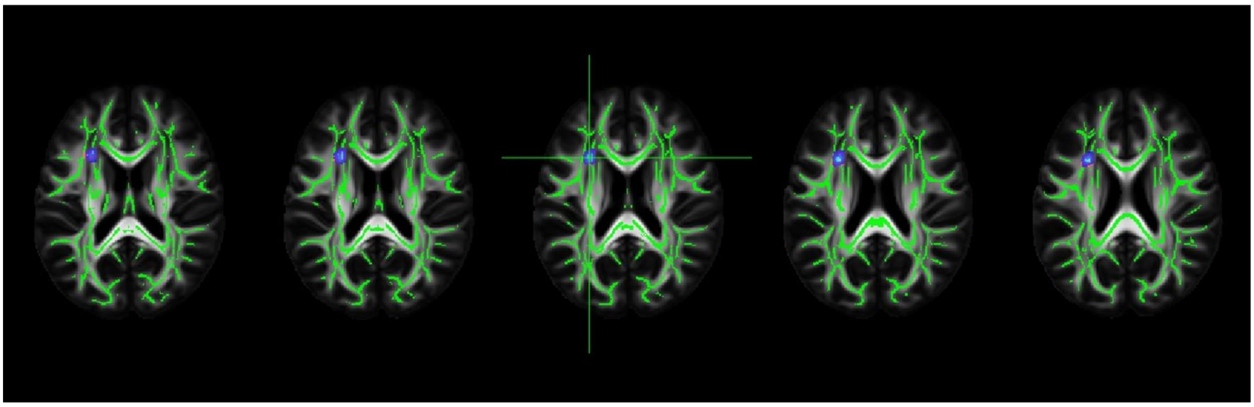

Figure S14. Left Global Mean RNFLT negative correlations with Mean Diffusivity **with and without CVR factors** controlling (n=550). **Upper** figure shows the correlations when controlling for the CVR factors i.e., BMI, LDL and HDL Cholesterol scores and Diabetes, Hypertension, Smoking and Physical Activity status (blue) in addition to age, sex, total intracranial volume and retina scan radius. **Middle** figures show the correlations when controlling for only age, sex and related retina scan radius (pink)**. Lower** figure shows both correlations together (overlap, purple). Results shown, at cluster-level corrected p< 0.05 for FWER with an uncorrected p<0.001 voxel-level clustering threshold, on MNI-registered FSL_HCP1065_FA_1mm standard atlas, Neurological View. RNFLT: Retinal Nerve Fiber Layer Thickness

Right Global RNFLT x Brain MD (-28, 16, 20) with CVR control

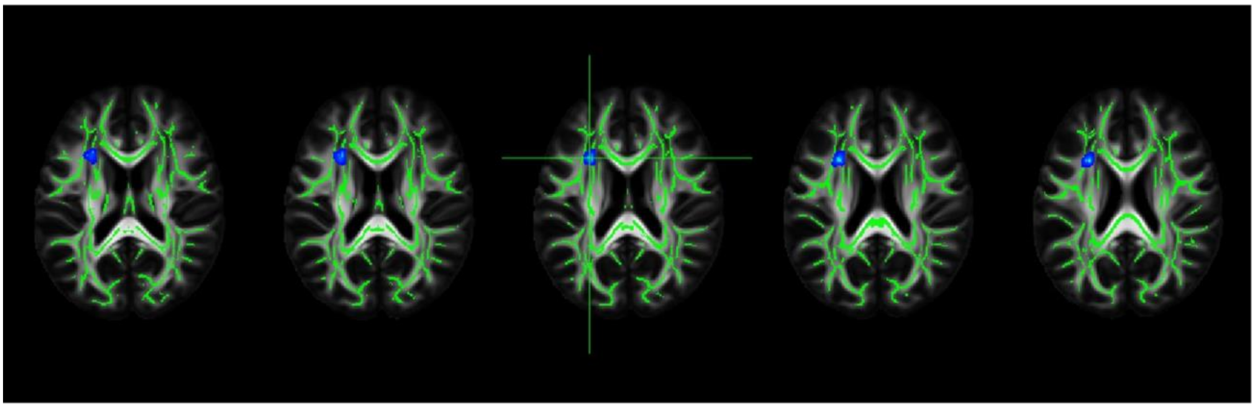

Right Global RNFLT x Brain MD (-28, 16, 20) without CVR control

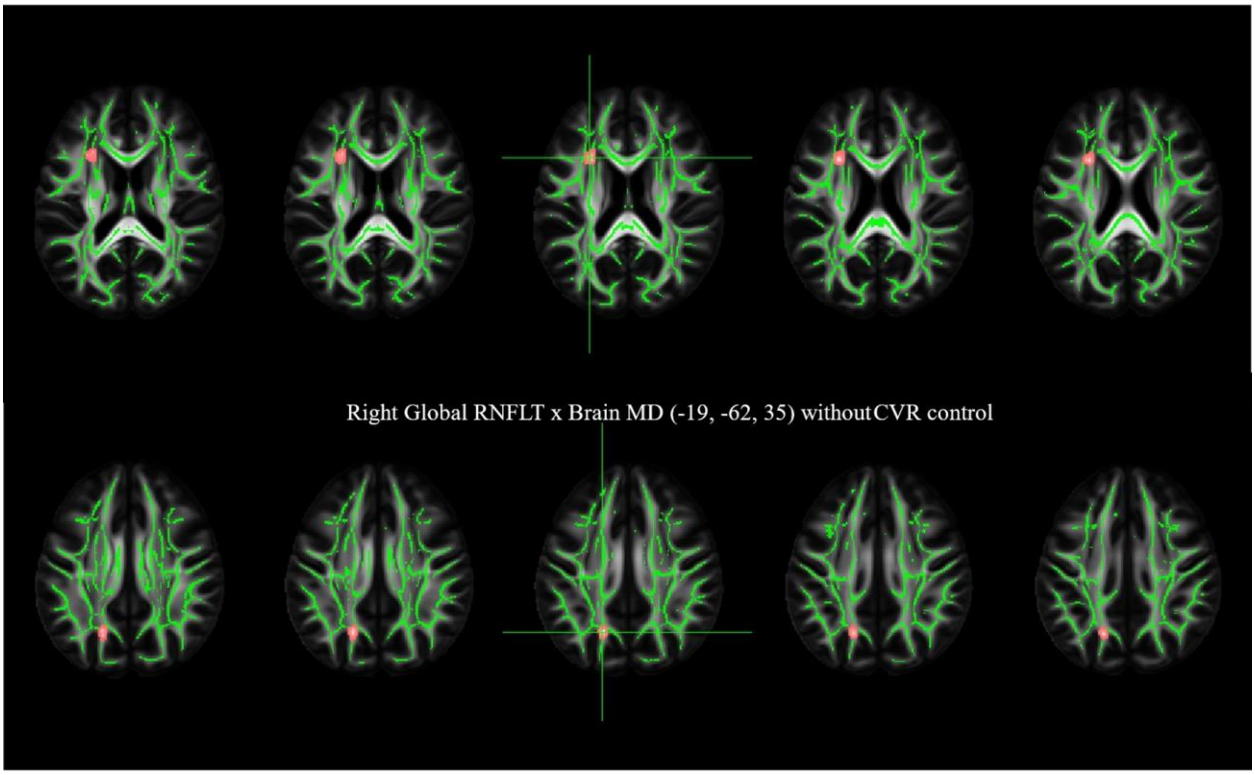


Right Global RNFLT x Brain MD (-28, 16, 20) with and without CVR control (together)

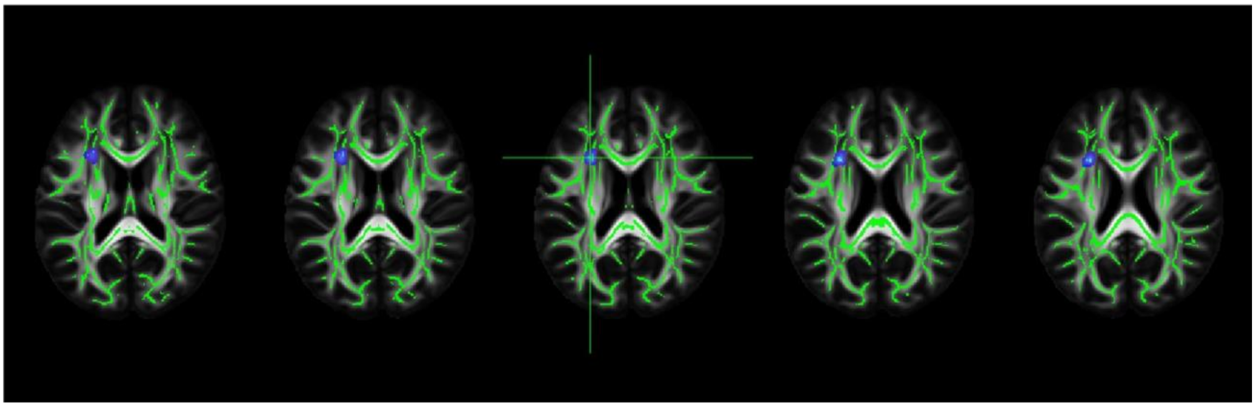

Figure S15. Right Global Mean RNFLT negative correlations with Mean Diffusivity **with and without CVR factors** controlling (n=550). **Upper** figure shows the correlations when controlling for the CVR factors i.e., BMI, LDL and HDL Cholesterol scores and Diabetes, Hypertension, Smoking and Physical Activity status (blue) in addition to age, sex, total intracranial volume and retina scan radius. **Middle** figures show the correlations when controlling for only age, sex and related retina scan radius (pink)**. Lower** figure shows both correlations together (overlap, purple). Results shown, at cluster-level corrected p< 0.05 for FWER with an uncorrected p<0.001 voxel-level clustering threshold, on MNI-registered FSL_HCP1065_FA_1mm standard atlas, Neurological View. RNFLT: Retinal Nerve Fiber Layer Thickness

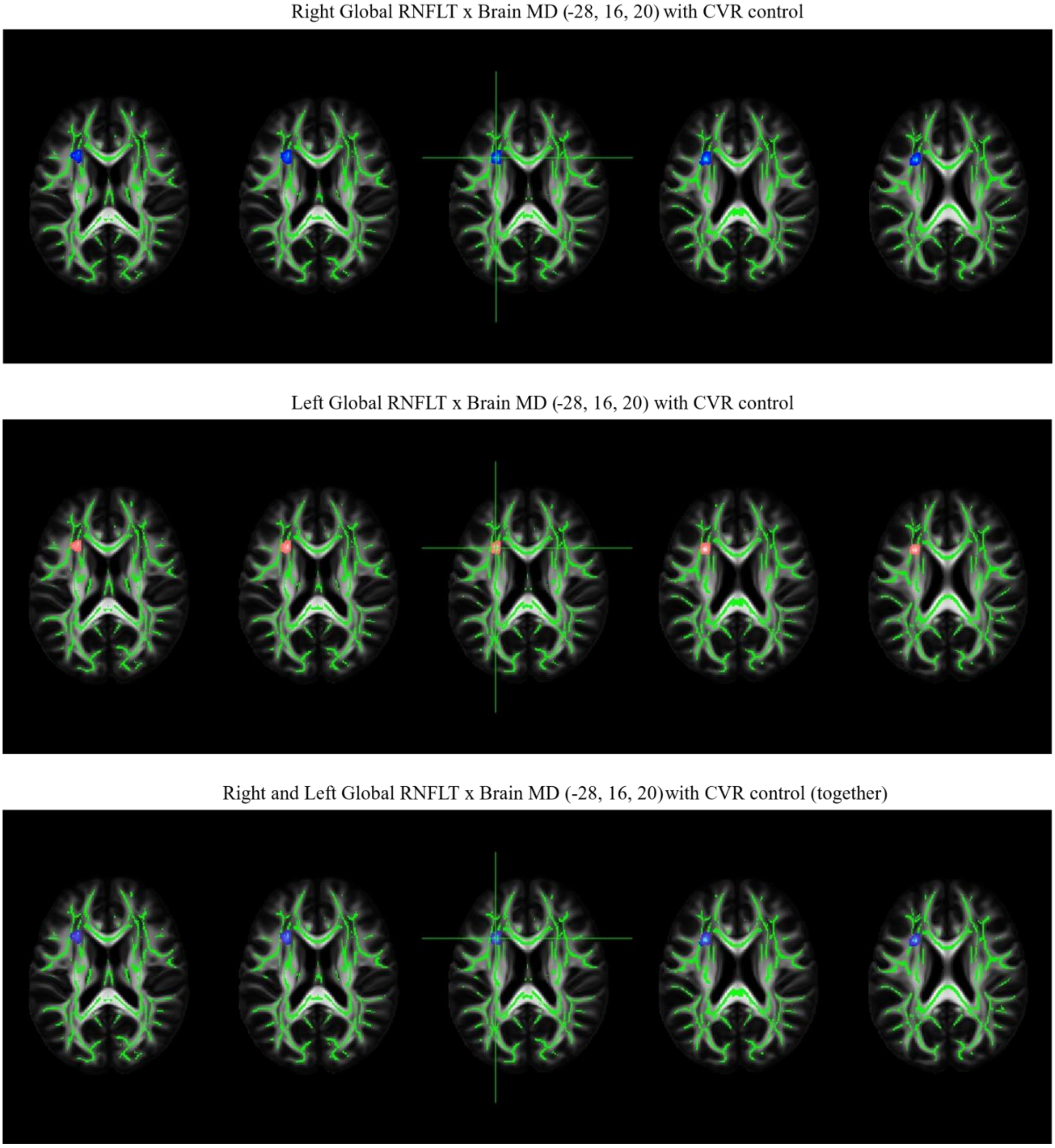


Figure S16. Comparison of the **Right** and **Left Global Mean RNFLT** negative correlations with the brain MD when controlling for the CVR factors i.e., BMI, LDL and HDL Cholesterol scores and Diabetes, Hypertension, Smoking and Physical Activity status (**i.e., with CVR**) in addition to age, sex and related retina scan radius **(n=550).** Results shown, at cluster-level corrected p< 0.05 for FWER with an uncorrected p<0.001 voxel-level clustering threshold, on MNI-registered FSL_HCP1065_FA_1mm standard atlas, Neurological View. RNFLT: Retinal Nerve Fiber Layer Thickness, MD: Mean Diffusivity

Right Global RNFLT x Brain MD (-28, 16, 20) without CVR control

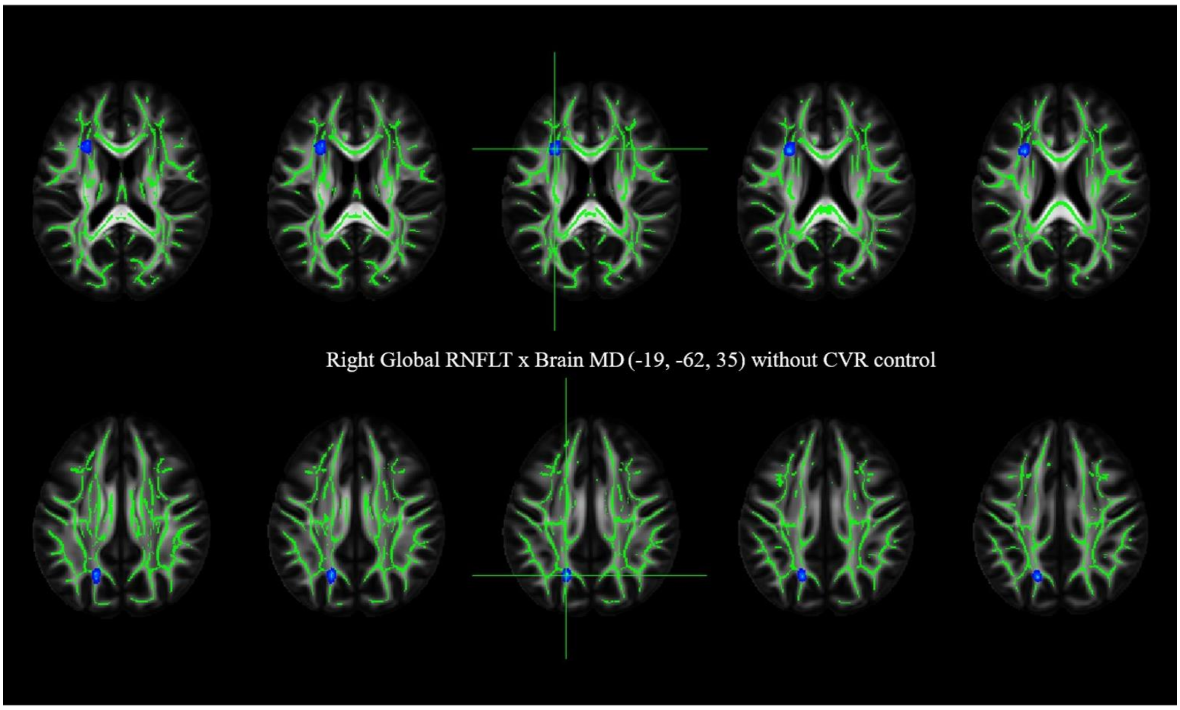


Left Global RNFLT x Brain MD (-28, 16, 20) without CVR control

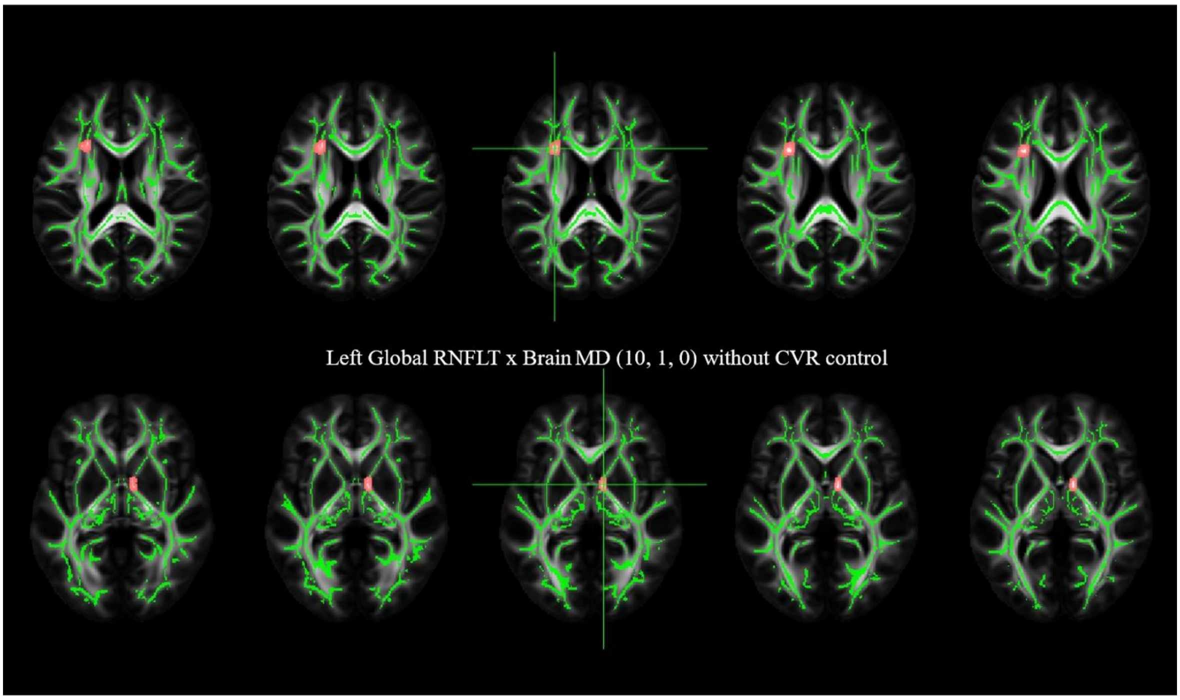


Right and Left Global RNFLT x Brain MD (-28, 16, 20) without CVR control (together)

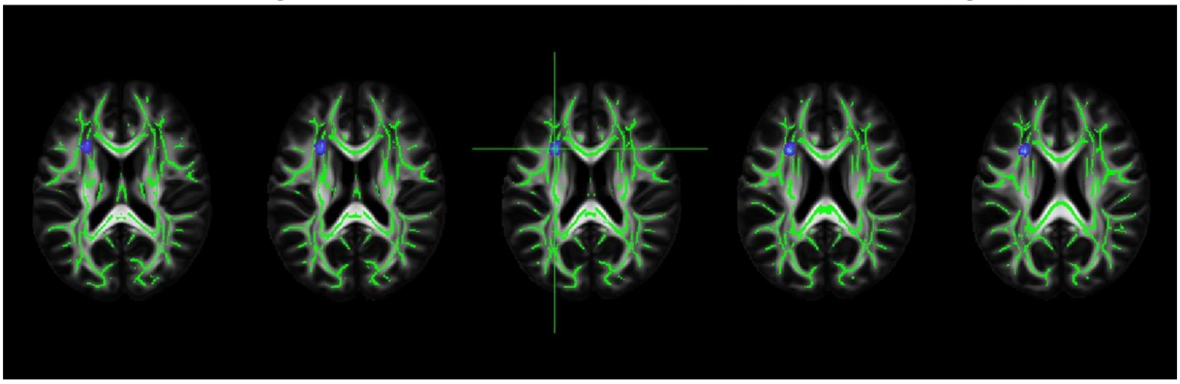

Figure S17. Comparison of the **Right** and **Left Global Mean RNFLT** negative correlations with the brain MD when controlling for only age, sex and regarding retina scan radius [(**i.e., without CVR, (n=550)]. ** Results shown, at cluster-level corrected p< 0.05 for FWER with an uncorrected p<0.001 voxel-level clustering threshold, on MNI-registered FSL_HCP1065_FA_1mm standard atlas, Neurological View. RNFLT: Retinal Nerve Fiber Layer Thickness, MD: Mean Diffusivity

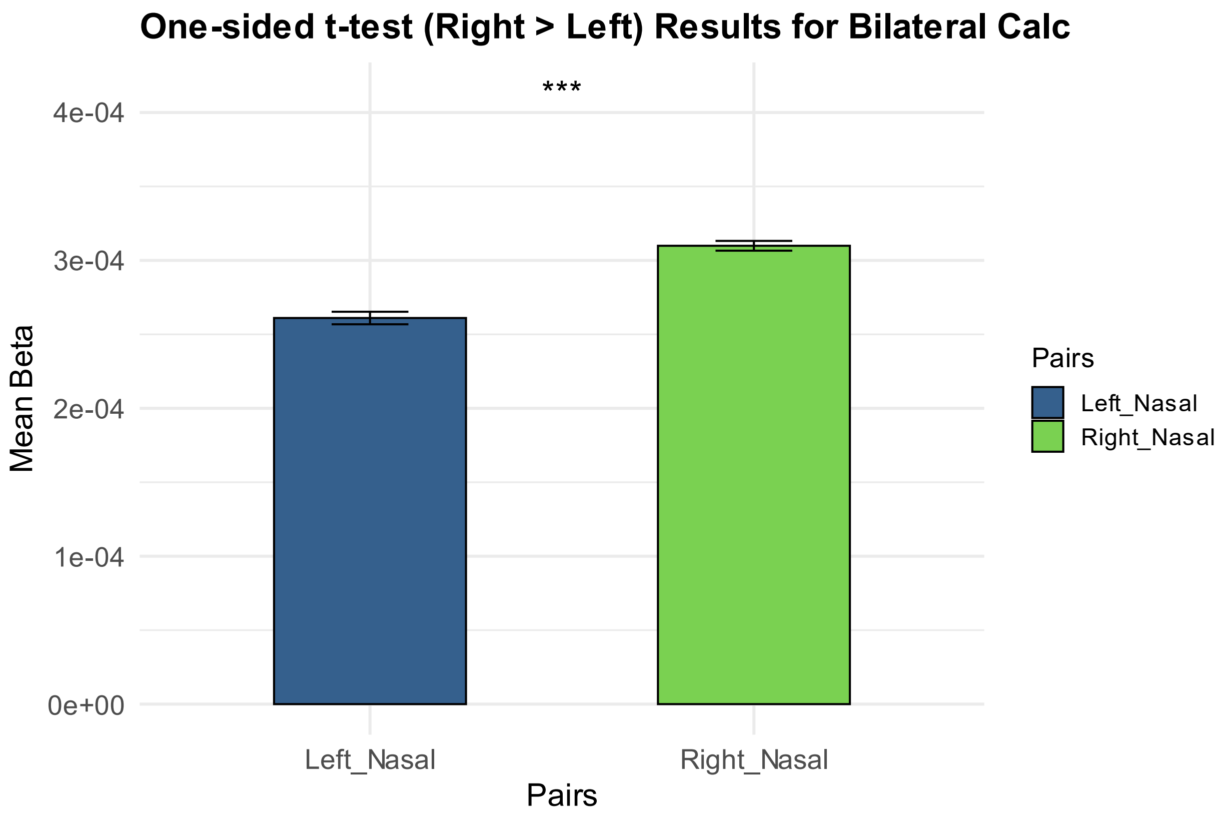


Figure S18. Paired sample t-test results between the Left and Right Nasal RNFLT correlations with the GMD for the Bilateral Calcarine Cortex. RNFLT: Retinal Nerve Fiber Layer Thickness, GMD: gray matter density.

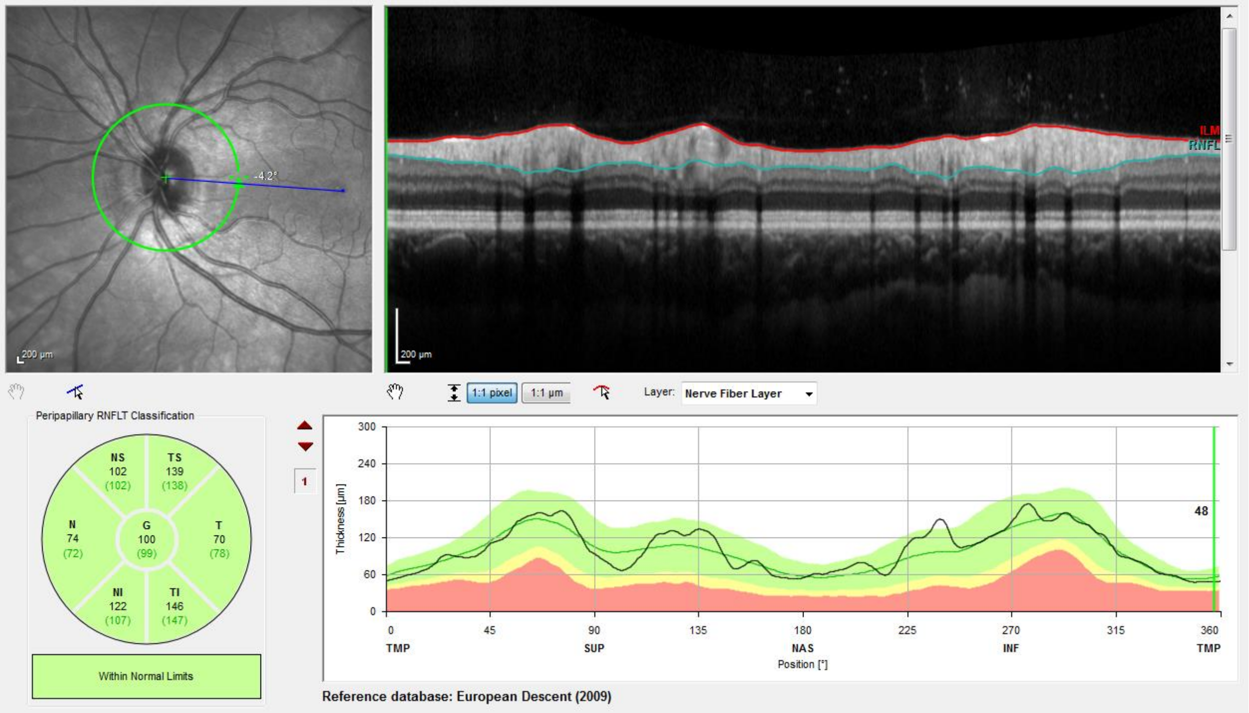


Figure S19. An example image of a circumpapillary RNFLT from an OCT scan (by Franziska G. Rauscher). RNFLT: retinal nerve fibre layer thickness, OCT: optical coherence tomography.

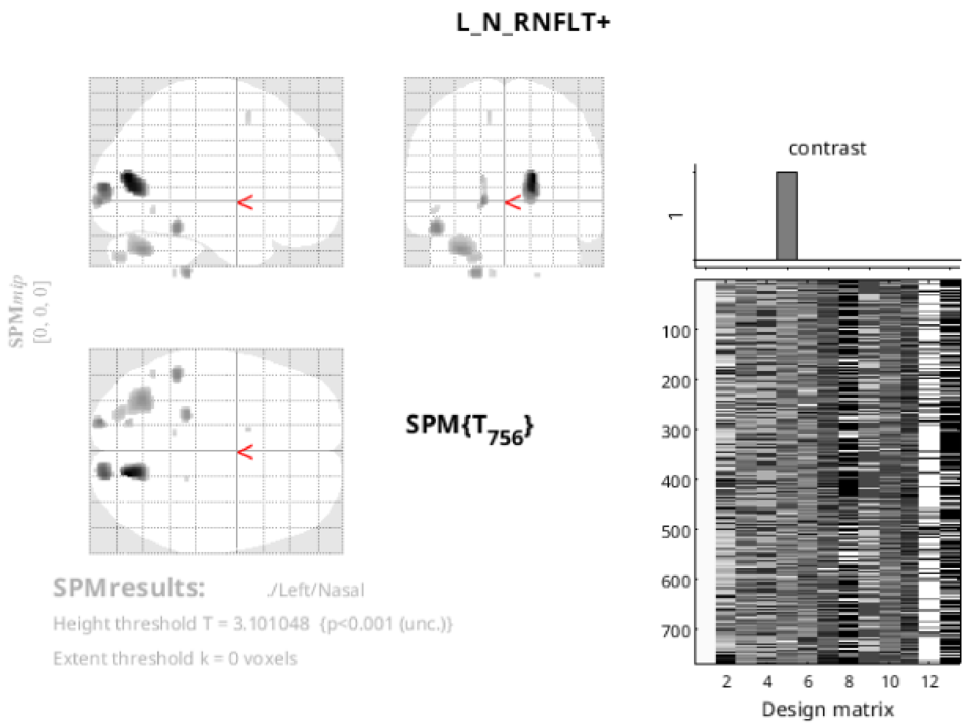


**Statistics:** ***p-values adjusted for search volume***

| set-level | | cluster-level | | | | peak-level | | | | | mm mm mm |
|---|---|---|---|---|---|---|---|---|---|---|---|
| $p$ | $c$ | $p_{FWE-corr}$ | $q_{FDR-corr}$ | $k_E$ | $p_{uncorr}$ | $p_{FWE-corr}$ | $q_{FDR-corr}$ | $T$ | $(Z_E)$ | $p_{uncorr}$ | |
| 0.021 | 12 | 0.378 | 0.472 | 333 | 0.079 | 0.022 | 0.047 | 4.76 | 4.72 | 0.000 | 18 -75 14 |
| | | 0.552 | 0.533 | 235 | 0.133 | 0.270 | 0.339 | 4.07 | 4.05 | 0.000 | 18 -92 8 |
| | | 0.949 | 0.844 | 49 | 0.492 | 0.593 | 0.534 | 3.77 | 3.75 | 0.000 | -16 -94 -2 |
| | | 0.806 | 0.816 | 122 | 0.272 | 0.631 | 0.534 | 3.73 | 3.72 | 0.000 | -48 -42 -18 |
| | | 0.187 | 0.413 | 508 | 0.034 | 0.753 | 0.534 | 3.62 | 3.61 | 0.000 | -32 -66 -33 |
| | | 0.923 | 0.844 | 65 | 0.425 | 0.774 | 0.534 | 3.60 | 3.59 | 0.000 | -21 -36 -48 |
| | | 0.975 | 0.919 | 28 | 0.614 | 0.958 | 0.879 | 3.34 | 3.33 | 0.000 | -16 -76 14 |
| | | 0.897 | 0.844 | 79 | 0.377 | 0.970 | 0.879 | 3.31 | 3.30 | 0.000 | -22 -81 -39 |
| | | 0.992 | 0.919 | 8 | 0.809 | 0.988 | 0.879 | 3.22 | 3.21 | 0.001 | -12 6 56 |
| | | 0.994 | 0.919 | 5 | 0.857 | 0.990 | 0.879 | 3.21 | 3.20 | 0.001 | -9 -45 -48 |
| | | 0.992 | 0.919 | 8 | 0.809 | 0.994 | 0.879 | 3.16 | 3.15 | 0.001 | -48 -58 -8 |
| | | 0.996 | 0.919 | 2 | 0.919 | 0.995 | 0.879 | 3.15 | 3.14 | 0.001 | 21 -36 -50 |

*table shows 3 local maxima more than 8.0mm apart*

Height threshold: T = 3.10, p = 0.001 (0.998)
Extent threshold: k = 0 voxels
Expected voxels per cluster, <k> = 109.162
Expected number of clusters, <c> = 6.03
FWEp: 4.553, FDRp: 4.759, FWEc: Inf, FDRc: Inf

Degrees of freedom = [1.0, 756.0]
FWHM = 14.8 14.6 14.5 mm mm mm; 9.9 9.8 9.6 {voxels}
Volume: 1489158 = 441232 voxels = 419.0 resels
Voxel size: 1.5 1.5 1.5 mm mm mm; (resel = 929.93 voxels)

Figure S20. Left nasal RNFLT result map at uncorrected p < 0.001 level when all CVRF controlled for in addition to age, sex, TIV and left retina scanning radius in the VBM multiple regression model.

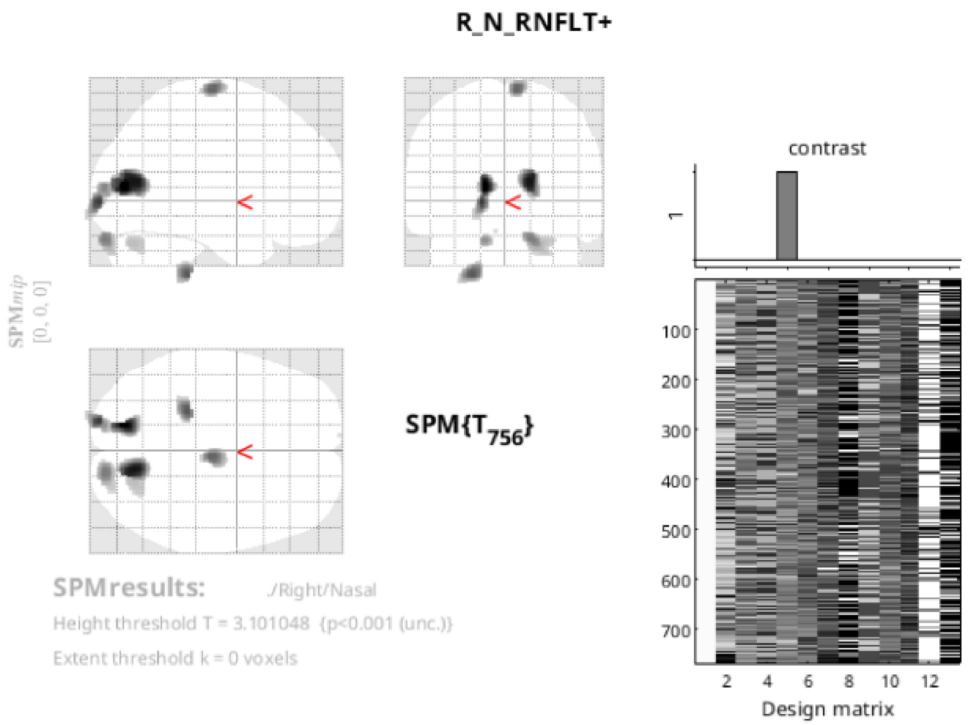


**Statistics:** ***p-values adjusted for search volume***

| set-level | | cluster-level | | | | peak-level | | | | | mm mm mm | | |
|---|---|---|---|---|---|---|---|---|---|---|---|---|---|
| $p$ | $c$ | $p_{FWE-corr}$ | $q_{FDR-corr}$ | $k_E$ | $p_{uncorr}$ | $p_{FWE-corr}$ | $q_{FDR-corr}$ | $T$ | $(Z_E)$ | $p_{uncorr}$ | | | |
| 0.087 | 10 | 0.363 | 0.264 | 343 | 0.075 | 0.010 | 0.020 | 4.94 | 4.90 | 0.000 | -15 | -78 | 8 |
| | | 0.141 | 0.252 | 579 | 0.025 | 0.028 | 0.028 | 4.69 | 4.66 | 0.000 | 15 | -72 | 10 |
| | | 0.688 | 0.276 | 173 | 0.193 | 0.098 | 0.051 | 4.38 | 4.35 | 0.000 | -18 | -96 | -3 |
| | | 0.510 | 0.264 | 256 | 0.118 | 0.219 | 0.098 | 4.14 | 4.12 | 0.000 | -24 | -36 | -50 |
| | | 0.616 | 0.264 | 205 | 0.159 | 0.257 | 0.098 | 4.09 | 4.07 | 0.000 | 6 | -20 | 72 |
| | | 0.482 | 0.264 | 271 | 0.109 | 0.466 | 0.178 | 3.87 | 3.85 | 0.000 | 16 | -92 | -26 |
| | | | | | | 0.978 | 0.686 | 3.28 | 3.27 | 0.001 | 26 | -87 | -34 |
| | | 0.756 | 0.292 | 144 | 0.233 | 0.574 | 0.212 | 3.78 | 3.76 | 0.000 | 18 | -92 | 8 |
| | | 0.930 | 0.489 | 61 | 0.440 | 0.627 | 0.218 | 3.74 | 3.72 | 0.000 | -15 | -93 | -27 |
| | | 0.571 | 0.264 | 226 | 0.140 | 0.947 | 0.584 | 3.37 | 3.36 | 0.000 | 21 | -72 | -30 |
| | | 0.997 | 0.948 | 1 | 0.948 | 0.997 | 0.967 | 3.11 | 3.10 | 0.001 | -8 | 68 | -21 |

*table shows 3 local maxima more than 8.0mm apart*

Height threshold: T = 3.10, p = 0.001 (0.998)
Extent threshold: k = 0 voxels
Expected voxels per cluster, <k> = 109.023
Expected number of clusters, <c> = 6.04
FWEp: 4.553, FDRp: 4.597, FWEc: Inf, FDRc: Inf

Degrees of freedom = [1.0, 756.0]
FWHM = 14.8 14.6 14.5 mm mm mm; 9.9 9.8 9.6 {voxels}
Volume: 1489158 = 441232 voxels = 419.6 resels
Voxel size: 1.5 1.5 1.5 mm mm mm; (resel = 928.74 voxels)

Figure S21. Right nasal RNFLT result map at uncorrected p < 0.001 level when all CVRF controlled for in addition to age, sex, TIV and right retina scanning radius in the VBM multiple regression model.

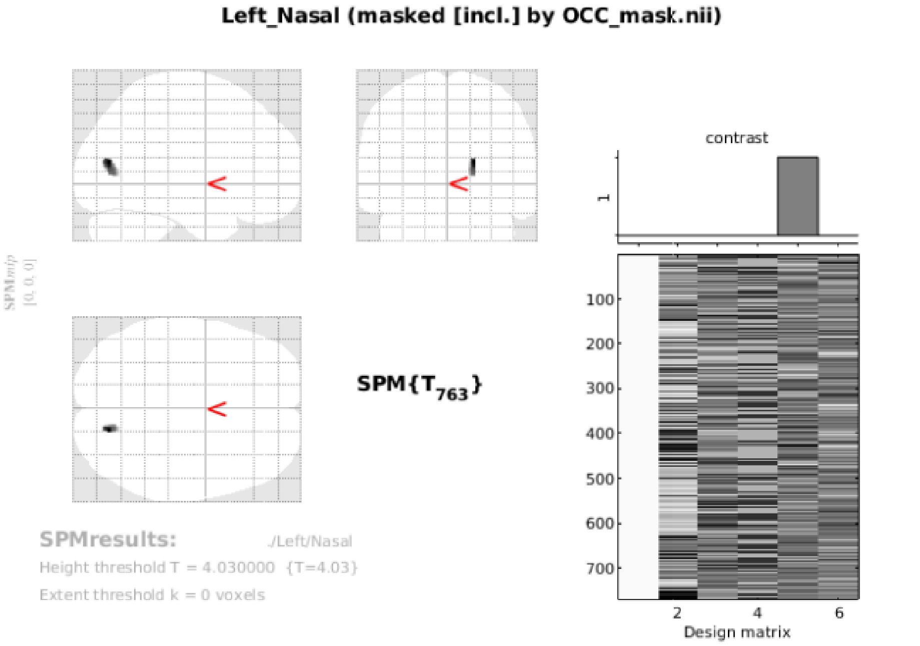


**Statistics:** ***search volume: image mask: ./OCC_mask.nii***

| cluster-level | | | | peak-level | | | | | mm mm mm |
|---|---|---|---|---|---|---|---|---|---|
| $p_{FWE-corr}$ | $q_{FDR-corr}$ | $k_E$ | $p_{uncorr}$ | $p_{FWE-corr}$ | $q_{FDR-corr}$ | $T$ | $(Z_E)$ | $p_{uncorr}$ | |
| 0.015 | 0.286 | 52 | 0.286 | 0.007 | 0.141 | 4.55 | 4.51 | 0.000 | 18 -75 14 |

*table shows 16 local maxima more than 4.0mm apart*

Height threshold: T = 4.03, p = 0.000 (0.050)
Extent threshold: k = 0 voxels
Expected voxels per cluster, <k> = 49.424
Expected number of clusters, <c> = 0.05
FWEp: 4.029, FDRp: Inf, FWEc: 52, FDRc: Inf

Degrees of freedom = [1.0, 763.0]
FWHM = 14.9 14.7 14.5 mm mm mm; 9.9 9.8 9.7 {voxels}
Volume: 156158 = 46269 voxels = 48.4 resels
Voxel size: 1.5 1.5 1.5 mm mm mm; (resel = 938.25 voxels)

Figure S22. Left nasal RNFLT result map with small volume correction at cluster-level pFWE < 0.05 and at uncorrected voxel-level p < 0.001 when all CVRF controlled for in addition to age, sex, TIV and left retina scanning radius in the VBM multiple regression model.

**Right_Nasal+ (masked [incl.] by OCC_mask.nii)**

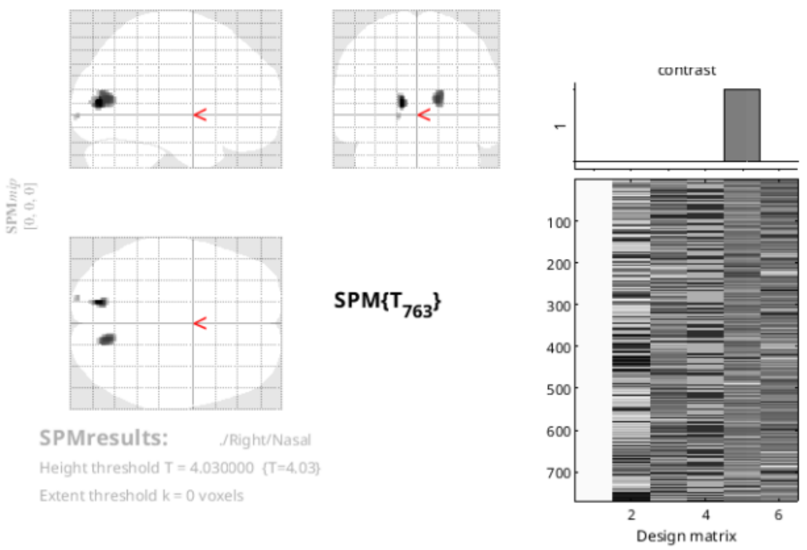


**Statistics:** ***search volume: image mask: ./OCC_mask.nii***

| set-level | | cluster-level | | | | peak-level | | | | | mm mm mm |
|---|---|---|---|---|---|---|---|---|---|---|---|
| $p$ | $c$ | $p_{FWE-corr}$ | $q_{FDR-corr}$ | $k_E$ | $p_{uncorr}$ | $p_{FWE-corr}$ | $q_{FDR-corr}$ | $T$ | $(Z_E)$ | $p_{uncorr}$ | |
| 0.000 | 3 | 0.007 | 0.198 | 107 | 0.132 | 0.002 | 0.154 | 4.85 | 4.81 | 0.000 | -15 -78 8 |
| | | 0.003 | 0.154 | 190 | 0.051 | 0.005 | 0.200 | 4.61 | 4.58 | 0.000 | 15 -72 10 |
| | | | | | | 0.008 | 0.200 | 4.53 | 4.50 | 0.000 | 16 -75 14 |
| | | 0.037 | 0.743 | 6 | 0.743 | 0.027 | 0.534 | 4.20 | 4.18 | 0.000 | -18 -96 -3 |

*table shows 16 local maxima more than 4.0mm apart*

Height threshold: T = 4.03, p = 0.000 (0.050)
Extent threshold: k = 0 voxels
Expected voxels per cluster, <k> = 49.364
Expected number of clusters, <c> = 0.05
FWEp: 4.030, FDRp: Inf, FWEc: 6, FDRc: Inf

Degrees of freedom = [1.0, 763.0]
FWHM = 14.9 14.7 14.5 mm mm mm; 9.9 9.8 9.7 {voxels}
Volume: 156158 = 46269 voxels = 48.4 resels
Voxel size: 1.5 1.5 1.5 mm mm mm; (resel = 937.11 voxels)

Figure S23. Right nasal RNFLT result map with small volume correction at cluster-level pFWE < 0.05 and at uncorrected voxel-level p < 0.001 when only age, sex, TIV and right retina scanning radius were controlled for in the VBM multiple regression model.

***
